# Robust Organ Shape During Growth Requires Local Morphogen Signaling and Global Curvature Feedback

**DOI:** 10.64898/2026.04.02.716206

**Authors:** Kristoffer Jonsson, Amir Porat, Viraj Alimchandani, Sahil Haque, Émile Julien, Henrik Jönsson, Yasmine Meroz, Anne-Lise Routier-Kierzkowska

**Author notes:** These authors contributed equally (Kristoffer Jonsson, Amir Porat, Viraj Alimchandani).

## Abstract

Precise organ shape during development requires feedback between organ-scale geometry and cell-level growth dynamics. Here, we use the apical hook of dicotyledonous plants as a model to study this multiscale control. This protective structure forms a ∼180° bend that is stably maintained despite rapid elongation. Combining quantitative growth mapping, multiscale modeling, and hormone analysis, we identify the cellular dynamics underlying hook maintenance. Contrary to the prevailing model, hook stability does not arise from sustained inner–outer growth asymmetry. Instead, stable curvature emerges from a biphasic, self-similar growth pattern that remains spatially fixed despite continuous cell flux. Auxin-driven growth asymmetry induces bending but is spatially restricted and insufficient to maintain shape. By contrast, a graded, auxin-independent autotropic response provides curvature-dependent growth regulation that counteracts bending and stabilizes shape against perturbations. Together, these findings establish a general principle in which local morphogen-driven bending and global geometry-dependent feedback jointly maintain robust organ shape during growth.

**Highlights:**

- Self-similar growth maintains apical hook curvature despite continuous cell flux
- Curvature stability emerges from coordinated axial and differential growth
- Auxin asymmetry is spatially restricted and insufficient for shape maintenance
- Curvature-dependent feedback ensures robust and optimized hook shape

## Introduction

How living systems generate precise and robust organ shapes, despite intrinsic variability and environmental perturbations remains a central challenge in developmental biology (Lander 2011; Hong et al. 2018; Moghe et al. 2026). Addressing this problem requires linking local cellular growth to global organ form—a connection that remains poorly understood. Slender, rod-like organs provide a tractable system, as their shape can be largely reduced to curvature along a single axis. Such curvature control is widespread, spanning systems from bacteria and neurons to multicellular structures such as the Drosophila hindgut and the *C. elegans* embryo (Wong et al. 2017; Oliveri and Goriely 2022; Seale et al. 2018; Inaki et al. 2018; Dai and Ben Amar 2024).

Plants offer powerful systems to investigate shape control. As cells are fixed within rigid walls, organ shape emerges solely from spatial patterns of cellular expansion, without the confounding effects of cell migration or rearrangements present in animal development (Coen and Cosgrove 2023; Kierzkowski et al. 2019; Silveira et al. 2025). Elongated organs such as roots and shoots bend in response to external signals, such as light or gravity, due to faster cell expansion on the outer (convex) side of the curve (Gilroy and Masson 2008; Jonsson et al. 2023, Harmer and Brooks 2018). This growth asymmetry is widely accepted as a reflection of the asymmetric distribution of the growth hormone auxin (Vanneste et al. 2025; Ottenschläger et al. 2003; Rakusová et al. 2011; Evans et al. 1994). However, a sustained growth asymmetry alone would lead to runaway curvature. To maintain stable shapes, bending must be counteracted by feedback. In organ-scale models, this is captured by autotropism—a curvature-dependent attenuation of growth asymmetry that acts as a damping term (Moulia et al. 2019). Yet how this geometric feedback emerges from cellular growth dynamics remains unknown.

The apical hook has been central to elucidating the molecular signaling of growth and shape control over the last several decades (Cao et al. 2019; Zádníková et al. 2010; Žádníková et al. 2016; Vandenbussche et al. 2010; Wang et al. 2020; Béziat et al. 2017; Du et al. 2022; Zhu et al. 2019; Abbas et al. 2018; Willige et al. 2012; H. Li et al. 2004; Gallego-Bartolomé et al. 2011; Mazzella et al. 2014; Vanneste et al. 2025; Walia et al. 2024; Stepanova et al. 2008; Lehman et al. 1996; Hua and Meyerowitz 1998; Baral et al. 2021; Guzmán and Ecker 1990; Shen et al. 2016; King et al. 1995; Kieber et al. 1993; Raz and Ecker 1999; Vriezen et al. 2004; Vain et al. 2019; Zhang et al. 2014). This transient ∼180° bend of the hypocotyl protects the shoot meristem during soil emergence (Shen et al. 2016; Abbas et al. 2013). Unlike responses to light or gravity, hook formation is developmentally programmed (Miyamoto et al. 2014; Jonsson et al. 2025). During the maintenance phase, the hook angle remains remarkably stable, offering a reproducible and quantifiable organ-scale phenotype (Vandenbussche et al. 2010; Mazzella et al. 2014).

Current models interpret both hook formation and maintenance through a sustained inner– outer growth asymmetry, in which auxin accumulates on the inner side to repress growth while the outer side elongates faster (Vandenbussche et al. 2010; Zádníková et al. 2010; Abbas et al. 2013; Gallego-Bartolomé et al. 2011). However, classical kinematic studies show that during maintenance, cells elongate continuously and are displaced along the hook (Silk and Erickson 1978; Raz and Ecker 1999). This creates a fundamental paradox: how can a stable organ-scale curvature be maintained in a tissue undergoing continuous growth and material flux?

Here, integrating live cell imaging, quantitative 3D analysis, and organ-scale modeling, we establish a framework for understanding the cellular basis of curvature control. We show that stable curvature does not arise from a static and uniform growth asymmetry, but from a dynamic steady state defined by self-similar growth, in which cells move through an asymmetric growth pattern that reverses along the organ. Quantitative auxin measurements and molecular perturbations show that, while auxin asymmetry is necessary for bending, it is insufficient for maintenance. Instead, curvature-dependent feedback directly couples organ geometry to local cell growth, stabilizing shape independently of auxin.

Together, these results overturn the prevailing model of hook maintenance, revealing a generalizable multiscale mechanism: stable form in growing organs emerges from dynamic feedback between local growth and global geometry.

## Results

### Cell-Level Growth Dynamics of Hook Maintenance Reveal a Biphasic Differential Growth Pattern

In darkness, germinating seedlings bend their young stem (hypocotyl) into an apical hook, shielding the stem cell niche during soil emergence. In *Arabidopsis*, the hypocotyl elongates several-fold while maintaining a nearly constant hook angle over the ∼48 h maintenance phase (**Fig. 1A–B**). As previously reported, the auxin signaling reporter DR5 accumulates along the inner side of the hook (Friml et al. 2002; Zádníková et al. 2010; Vandenbussche et al. 2010) (**Fig. 1C**). Auxin is a central plant growth hormone that regulates cell expansion, and current models propose that its asymmetric distribution represses growth on the inner side to maintain curvature (Vandenbussche et al. 2010; Žádníková et al. 2016; Abbas et al. 2013; Béziat and Kleine-Vehn 2018; Du et al. 2022) (**Fig. 1D**).

**Figure 1.**
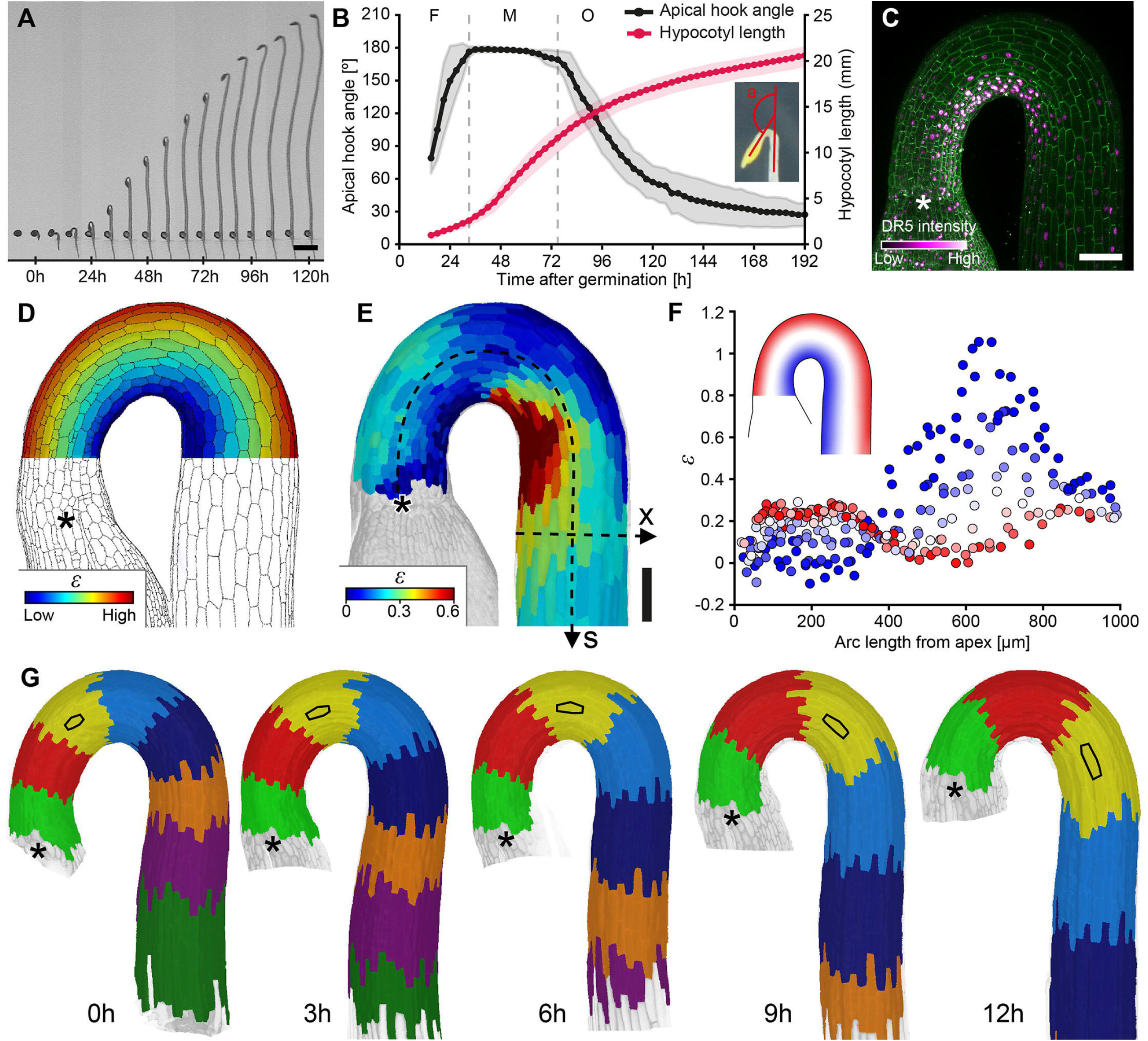
Direct growth measurements reveal complex dynamics underlying apical hook maintenance. **(A)** Time-lapse series of a dark-grown *Arabidopsis* wild-type (WT) seedling, exhibiting apical hook formation, maintenance and opening occurring concomitantly with rapid hypocotyl elongation. Depicted at 8 h intervals. Scale bar, 5 mm. **(B)** Kinematics of apical hook angle and hypocotyl length in WT seedlings. Measurements were performed at 3 h intervals. The inset depicts the definition of the apical hook angle (a). Vertical dotted lines demarcate hook developmental phases: formation (F), maintenance (M), and opening (O). Shaded areas indicate mean ±SD. *n* = 12 **(C)** DR5::Venus reporter indicating higher auxin response along the inner side of the hook at onset of hook maintenance (∼24 h after germination) (DR5, magenta; tdTomato, green). *n* = 16. Scale bar, 100 μm. **(D)** Schematic heatmap depicting binary strain (*ε*) pattern commonly inferred from DR5 observations. **(E)** Live cell tracking and spatiotemporal strain quantification in WT at onset of hook maintenance reveal a coordinated, spatially structured growth field inconsistent with a simple binary inner–outer asymmetry. Dotted line arrows indicate longitudinal (*s*) and radial (*x*) coordinates and direction. Scale bar, 100 μm. Additional biological replicates are shown in **Fig. S2** (*n* = 10). **(F)** Quantification of individual cell axial strain (*ε*) across the apical hook, from **(E)**. Data points are color-coded according to lateral position (outer side: red; inner side: blue; intermediate positions: white). Inset: schematic indicating the lateral red-to-blue gradient across the hook. **(G)** Kinematic tracking of organ regions from onset of hook maintenance (designated *t*= 0h), depicting the continuous flow of segments passing through the static hook region. Black contours highlight one individual cell over time. All asterisks indicate hypocotyl apex. Scale bar, 100 μm.

To test this prediction, we quantified hook growth dynamics at cellular resolution during the maintenance phase. Time-lapse confocal imaging of *Arabidopsis* seedlings (3 h intervals over 12 h) was performed under physiological dark conditions (561 nm illumination) (Mazzella et al. 2014). Using MorphoGraphX, we reconstructed the curved epidermal surface in 3D and quantified longitudinal strain (, relative length increase) for individual cells (Barbier de Reuille et al. 2015; Strauss et al. 2022) (**Fig. S1**). Unexpectedly, differential growth followed a biphasic pattern rather than a simple outer–inner gradient (**Fig. 1E–F, Fig. S2**). Cells elongated faster on the outer side near the apex, but this gradient reversed basally, where inner-side cells grew faster. This reversal challenges the prevailing model in which auxin-mediated growth inhibition along the inner side maintains hook curvature.

Tracking cells across the full 12 h sequence revealed dynamics over longer time scales. Despite a stable hook shape, cells and tissue segments progressively moved through the hook from apex to base (**Fig. 1G, Fig. S3, Movie S1–S2**), consistent with early organ-level descriptions of hook growth (Silk and Erickson 1978; Raz and Ecker 1999). As plant cells do not migrate nor intercalate, and cell division is negligible at this stage (Gendreau et al. 2020; Bou Daher et al. 2018; Raz and Koornneef 2001), this displacement reflects cumulative cell elongation along the hypocotyl. Consequently, material segments undergo a characteristic curvature trajectory as they transit through the hook: initially straight near the apex, maximally curved in the mid-hook region, and straightening again as they exit the hook. To determine how complex cell growth patterns give rise to stable hook shape, we next established a quantitative framework linking cell-scale expansion to organ-scale curvature dynamics.

### Self-Similar Hook Shape Dynamics Results from Coordinated Axial and Differential Growth Patterns

Despite continuous cellular displacement, the spatial pattern of growth remained stationary over 12 h, while hook curvature was preserved during elongation, indicating self-similar growth (**Fig. 2A; Fig. S3**). Cells therefore follow a position-dependent biphasic growth program: along the inner side, elongation switches from slow to fast, whereas along the outer side it shows the opposite transition, with the switch at maximal curvature (**Movie S3**).

**Figure 2.**
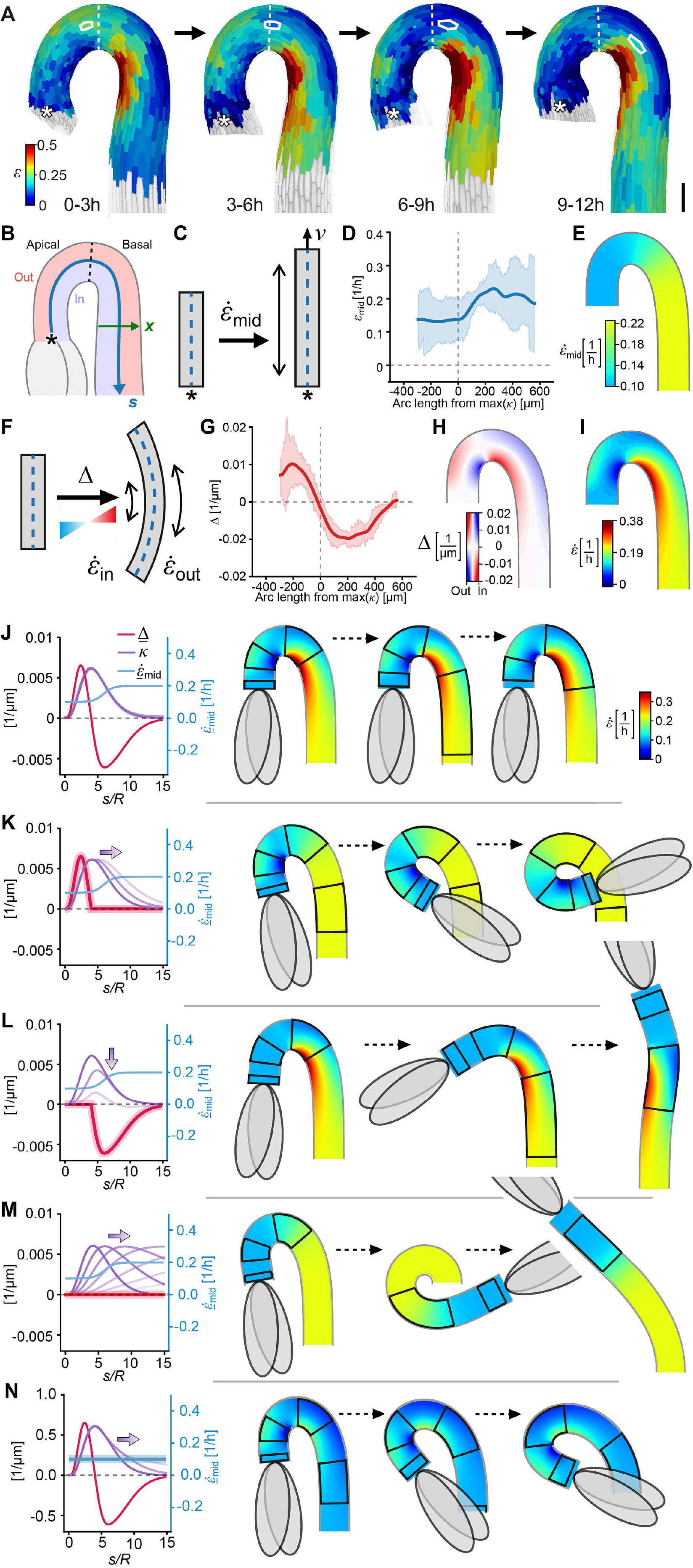
Hook maintenance requires precise coordination of local growth patterns. Heatmaps of incremental strain from onset of apical hook maintenance in WT (designated *t* = 0), depicting four consecutive 3h intervals. During the maintenance phase, the organ shape and patterns of growth are maintained while individual cells (white contours) pass through the hook. Dotted white lines indicate position of maximal curvature. Asterisks indicate hypocotyl apex. Scale bar, 100 μm. **(B-I)** We use a phenomenological kinematic model to decompose the observed pattern of cell elongation into two types of growth gradients: transverse and longitudinal. **(B)** Coordinate system for analysing hook growth: longitudinal axis along hook midline (blue arrow) defining the curvilinear position (*s*) from the apex. Dotted line indicates the position of maximal curvature, delimiting the apical and basal regions. Lateral position (*x*) is defined by a local axis perpendicular to the longitudinal curve (green arrow), with negative *x* towards the inner side of the hook, positive towards the outer side, and midline corresponding to *x*= 0. **(C)** Relative elongation rate of the hook midline, 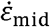, which causes the cells to move along the hook with respect to a reference fixed point (asterisk) with velocity *v*. **(D)** Measurements of 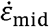 along the hook length in WT from onset of maintenance, over 12h, corresponding to **(A)**. **(E)** Idealized profile of 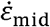 used in the kinematic model. **(F)** Transverse normalized gradient of elongation, 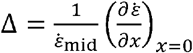, or relative difference in cell elongation between the outer and inner sides of the hook, which drives changes in curvature of material segments. Here we show a positive Δ, associated with an increase in signed curvature. **(G)** Measurements of Δ along the hook length in WT from onset of hook maintenance, over 12h, corresponding to **(A)**. Solid line: average curve of the 4 consecutive intervals, shaded area: confidence interval. **(H)** Idealized profile of Δ used in the kinematic model. Rather than assigning a uniform colour to each cross-section based on Δ(*s*), the local color represents 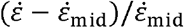 (see Eq.S13 in the SI), i.e., the relative difference between the local growth rate and the midline growth rate. The heatmap therefore visualises the normalised lateral growth gradient and its spatial pattern, with color intensity indicating the magnitude of Δ (positive or negative) as depicted by the colorbar. **(I)** Spatial distribution of local relative elongation rates 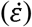 obtained by the idealized profiles of Δ in **(H)** and 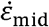 in **(E)**, based on Eq.S13 in the SI. **(J)** Kinematic model of WT maintenance: we assume a profile of 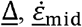 and initial shape *κ*that reproduces the hook shape, where underline symbols denoted time independent, self similar profiles. The model (Eqs.5,6) shows that inner/outer differential growth in the apical and basal regions must be exactly balanced with midline elongation rates to maintain hook shape. Left: profiles over arc length, normalized by the model organ’s radius (*R* = 100 μm). Right: three snapshots of the shape dynamics of a model organ, progressing in time from left to right. Black outlines on the organ indicate material segments which translate down the hook and elongate over time. See also supplementary movies 4-8. **(K)** In the absence of negative <u>Δ</u>, the hook shape cannot be maintained and progressively exaggerates. The curvature profile *κ* transitions from dark to light purple on the graph, corresponding to the shapes shown on the right. **(L)** In the absence of positive <u>Δ</u>, the hook shape cannot be maintained and the hook progressively opens. **(M)** When <u>Δ</u> = 0, the curvature *κ* propagates basipetally following the one-way wave equation, initially causing hyperbending followed by gradual opening. **(N)** Modification of the midline strain profile 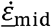 disrupts maintenance and results in hook exaggeration.

To understand this behavior, we developed a framework linking cellular growth to organ-scale curvature (**Fig. 2B–I**). Treating the hypocotyl as a growing rod, hook shape emerges from the interplay between axial elongation, which advects tissue, and differential growth between inner and outer sides, which generates bending. While this approach builds on established rod models of roots and shoots, the growth gradients driving curvature have not been experimentally resolved (Moulia et al. 2006; Bastien et al. 2013; Porat et al. 2020; Bastien et al. 2014; Fozard et al. 2016; Chelakkot and Mahadevan 2017; Moulton et al. 2020; Agostinelli et al. 2021; Porat, Rivière, et al. 2024; Porat, Tekinalp, et al. 2024; Oliveri et al. 2024).

Cell positions were described in a dynamic curvilinear coordinate system aligned with the organ midline (**Fig. 2B**), with a longitudinal coordinate *s* (*s*=0 at the hook apex) and lateral coordinate *x* (*x*>0 outer side, *x*< 0 inner side). The hook was further partitioned into apical and basal regions based on the position of maximal curvature.

To distinguish organ elongation from growth asymmetry, we decomposed the measured growth pattern into two fields, one driving axial tissue displacement and the other controlling curvature. The midline elongation field 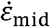 represents the longitudinal strain rate along the midline ( 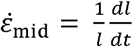, **Fig. 2C**). Although plant cells remain mechanically connected, cumulative elongation causes them to move away from the apex with velocity 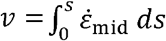. The second field, differential growth (Δ), captures the local growth asymmetry between the two sides of the organ (**Fig. 2F**). It is defined as the normalized lateral gradient of elongation: 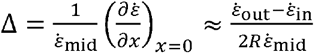, where 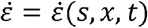 is the local elongation rate, 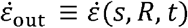 and 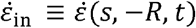 correspond to the outer and inner sides, and *R* is the hook radius.

Profiles of 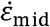 and Δ were extracted from cell growth data using local linear fits along the lateral coordinate (see **SI**). During maintenance, the elongation rate 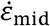 increases along the hook, with a sharp rise near the position of maximal curvature (**Fig. 2D; Fig. S4**), which we captured using an idealized spatial profile (**Fig. 2E**). Differential growth Δ delimits two domains: positive in the apical region and negative in the basal region, with a sign change at maximal curvature (**Fig. 2G,H; Fig. S4**). Combining idealized profiles of 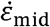 and Δ, the model accurately reproduces the growth patterns observed during hook maintenance (**Fig. 2I**).

Using this framework, we simulated hook dynamics at both the organ scale and the level of individual segments (**Fig. 2J–N, Movie S4–S8**). A segment initially located near the apex is displaced toward the base while its curvature *κ* evolves as:

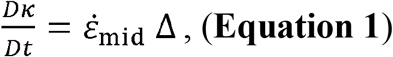

where *D*/*Dt* = ∂ / ∂*t* + *v*∂ / ∂*s* is the material derivative accounting for axial displacement (**see SI**). Thus, curvature dynamics in a segment depends both on differential growth Δ and midline elongation 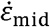.

We first modeled hook maintenance (**Fig. 2J**). During this phase, curvature changes of individual segments are coordinated with axial displacement, such that overall hook shape remains constant. In this regime, the system reaches a steady state of Eq. 1 in which curvature, differential growth, and elongation profiles are stationary and satisfy:

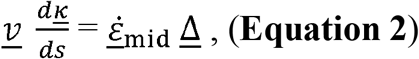

with all profiles maintained unchanged over time (denoted by an underline, see **SI**). Thus, bending and straightening are precisely balanced with axial displacement, allowing elongation while preserving the same curved shape. Consistent with Eq. 2, the position of maximal curvature coincides with the sign change of Δ, confirming self-similarity *in vivo* (**Fig. S5**). Using profiles of Δ and 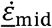 consistent with experimental data and Eq. 2, simulations reproduce curvature maintenance (**Fig. 2J**). As segments move along the hook, positive Δ in the apical region (faster outer-side growth) increases curvature, whereas negative Δ in the basal region (faster inner-side growth) drives straightening (**Movie S4**).

Next, we examined how perturbations of the growth fields affect hook maintenance (**Fig. 2K–N**). Removing the basal negative Δ led to progressive curvature increase, resembling hook exaggeration, whereas retaining only negative Δ caused progressive straightening and hook opening (**Fig. 2K–L, Movie S5-S6**). Setting Δ= 0 throughout the hook produced transient overbending followed by gradual straightening, as elongation advected curvature away from the hook region (**Fig. 2M, Movie S7**), consistent with a one-way wave equation governing curvature propagation (see **SI**). Finally, imposing a flat elongation profile while preserving Δ increased curvature (**Fig. 2N, Movie S8**), indicating that rapid basal elongation is also required to maintain hook shape.

We found that self-similar hook dynamics require tight coordination between axial elongation and differential growth. Outer-side growth in the apical region drives bending, while inner-side growth in the basal region drives straightening. At the segment level, these opposing dynamics must match axial displacement; disrupting this balance causes curvature runaway or premature opening. This framework links cellular growth to organ shape and provides a quantitative basis for phenotyping hook defects. Next, we applied this framework to mutants.

### High-Resolution Phenotyping Reveals Distinct Cell Growth Dynamics in Overbending Mutants

To test whether our framework can uncover the growth mechanisms underlying hook phenotypes, we analyzed two mutants with exaggerated curvature: *axr2-1* and *ctr1-1*, which affect auxin and ethylene signaling, respectively (Tiwari et al. 2001; Kieber et al. 1993) (**Fig. 3**). Despite targeting distinct pathways, both mutants reach substantially higher hook angles than WT before opening (**Fig. 3A,G**). This phenotypic similarity suggests a shared underlying growth mechanism.

**Figure 3.**
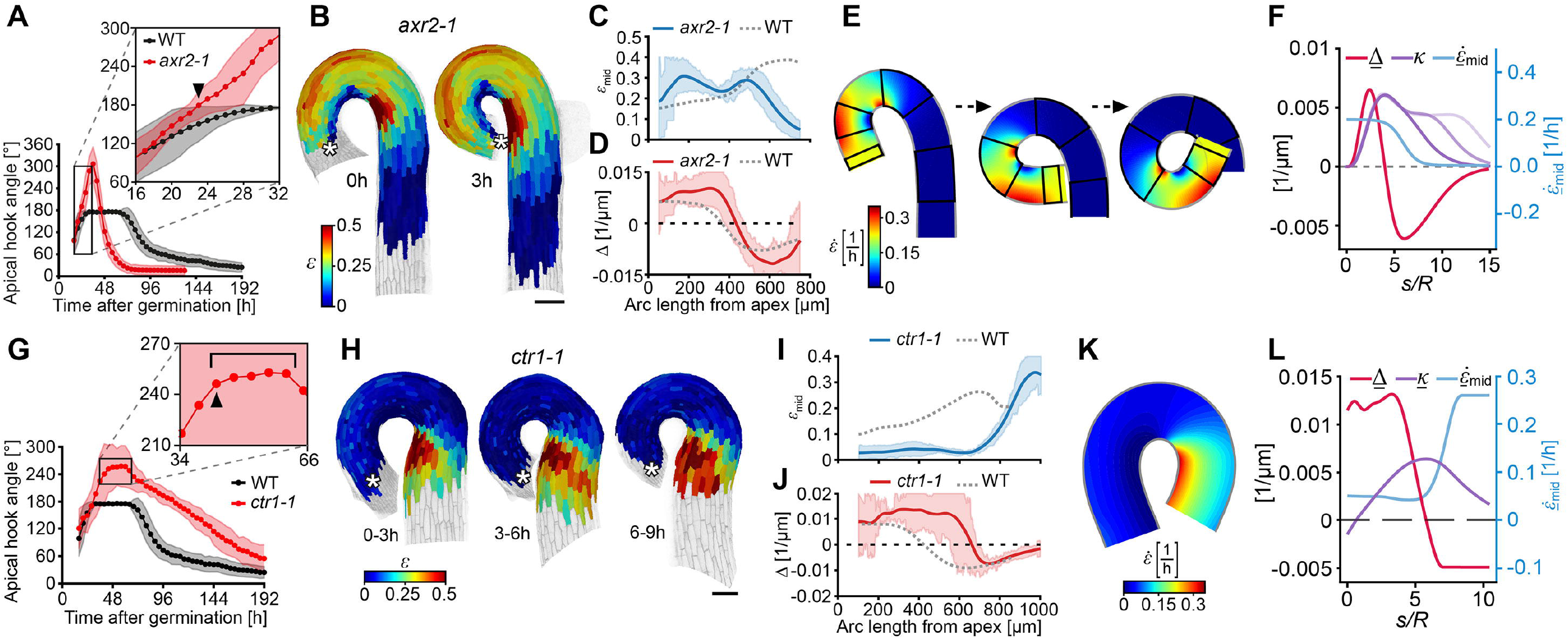
*axr2-1* and *ctr1-1* Exhibit Divergent Growth Dynamics Despite Similar Exaggerated Hook Angles. **(A)** Apical hook angle in WT and *axr2-1*. Inset highlights timepoint (arrowhead) where *axr2-1* angles deviate from WT. Shaded areas indicate mean ± SD. Measurements were performed at 3 h intervals. For inset interval, measurements were performed at 1 h intervals. *n* = 10. **(B)** Strain heatmaps of *axr2-1* during the 0–3 h interval corresponding to the onset of hook exaggeration (designated ***t***= **o**h) (arrowhead in **(A)** inset). The same growth pattern is projected onto both the 0 h and 3 h organ meshes for comparison with WT maintenance. Asterisks indicate the hypocotyl apex. Scale bar, 100 μm. **(C)** Average strain profile (*ε*_mid_) in WT and *axr2-1*, showing an apical-basal decline. *n* = WT 5, *axr2-1* 5. **(D)** Δ profile in *axr2-1*, which remains largely similar to WT. Dotted horizontal line indicates Δ= 0. *n* = WT 5, *axr2-1* 5. **(E-F)** Model simulation of the *axr2-1* phenotype, presented in a similar fashion to **Fig. 2J-N**. Here, the WT Δ profile is retained while the midline strain rate 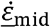 profile is inverted, leading to progressive hook exaggeration over time. **(G)** Apical hook angle in WT and *ctr1-1*. Inset highlights the timepoint where *ctr1-1* reaches a stable hook angle plateau (designated *t* = 0 h) (indicated by arrowhead). **(H)** Strain heatmaps of *ctr1-1* during three consecutive 3 h intervals after reaching a stable shape plateau, illustrating stable growth patterns. Asterisks indicate the hypocotyl apex. Scale bar, 100 μm. **(I)** Average strain (*ε*_mid_) profile in *ctr1-1* showing low strain in the apical region and a basal-ward displacement of the *ε*_mid_ acceleration relative to WT. *n* = WT 4, *axr2-1* 4. **(J)** Δ profile in *ctr1-1*, with the sign change (Δ= 0) displaced basally in coordination with the shift in *ε*_mid_. *n* = WT 4, *axr2-1* 4. **(K)** Model simulation reproducing the self-similar state of *ctr1-1*. **(L)** The modeling profiles used in **(K)**. Here, 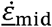 and Δ match the average experimental profiles from **(I-J)** and were used to derive the self similar curvature profile using our model. To reproduce the experimental shape, we set *R* = 150 μm, and added small initial values for the curvature (*R<u>κ</u>* (*s* = 0) = -0.15) and velocity 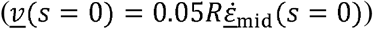.

First, we examined the *axr2-1* mutant (iaa7 gain-of-function), which exhibits exaggerated hook curvature (peak angle: 307° ± 47°, in *axr2-1*, n = 15 vs. 177° ± 0.54 in WT, n = 12) but fails to maintain a stable curvature plateau and opens rapidly (**Fig. 3A;** Ma et al. 2025; Vain et al. 2019). This allele stabilizes the Aux/IAA protein IAA7, rendering it resistant to auxin-induced degradation (Tiwari et al. 2001). We analysed *axr2-1* at the onset of divergence from WT, capturing the transition to exaggerated curvature (**Fig. 3A, inset; Fig. 3B**). Strikingly, while the differential growth profile (Δ) is similar to WT, the midline elongation profile 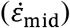 is inverted, with enhanced apical elongation and reduced basal elongation (**Fig. 3C– D**). Modeling identifies this inversion as sufficient to drive the phenotype: flipping 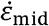 while preserving Δ breaks self-similarity and produces overbending (**Fig. 3E–F**). These results suggest that AXR2 primarily controls the apical–basal elongation profile during hook maintenance. Our interpretation is consistent with reported defects in hypocotyl elongation in *axr2-1* (Nagpal et al. 2000; Du et al. 2022), and contrasts with recent proposals that attribute the phenotype to altered inner–outer growth asymmetry (Ma et al. 2025).

Next, we analyzed the *ctr1-1* mutant, which carries a kinase□inactive allele of CTR1, a negative regulator of ethylene signaling (Kieber et al. 1993). Constitutive ethylene signaling in this mutant produces exaggerated hooks, similar to those induced by exogenous ethylene (Kieber et al. 1993; Vriezen et al. 2004). *ctr1*□*1* seedlings reached a peak hook angle of 258° ±45° (n = 12) compared to WT at 177.4° ± 0.6° (n= 10) (**Fig. 3G**). However, unlike *axr2-1, ctr1-1* maintains a stable—albeit exaggerated—curvature plateau before opening, indicating a modified but transient maintenance phase (plateau duration in *ctr1-1* : 15.3 ± 6.8 h, n = 12; in WT: 48.7 ± 8.0 h, n = 10). Cellular growth patterns during the *ctr1-1* maintenance phase differ markedly from both *axr2-1* and WT (**Fig. 3G inset; Fig. 3H**). Globally, the profiles of Δ and 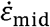 in *ctr1-1* resemble the WT self-similar organization (apical Δ> 0, basal Δ< 0, and 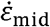 increasing basally). However, 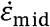 is reduced in the apical domain and both profiles are shifted ∼200 μm toward the base in *ctr1-1*, expanding the apical domain to roughly twice its WT length (**Fig. 3I–J**). Based on experimentally derived profiles, the model accurately reproduced the exaggerated yet stable curvature of *ctr1-1* (**Fig. 3K–L**). Thus, the phenotype arises from a spatial shift in the transition between bending and straightening domains.

Although *axr2-1* and *ctr1-1* produce similarly exaggerated hook angles, they arise from distinct growth mechanisms. Contrary to the prevailing view that both reflect enhanced auxin-driven inner–outer asymmetry (Vandenbussche et al. 2010; Žádníková et al. 2016; Ma et al. 2025), *axr2-1* displays an inversion of the apical–basal elongation profile that disrupts stable maintenance, whereas *ctr1-1* shows a basal shift of the bending–straightening boundary that enables transient maintenance of an exaggerated yet self-similar shape.

From a methodological standpoint, these results highlight the need for a new approach. Hook angle is a widely used readout of differential growth under genetic or environmental perturbations (Guzmán and Ecker 1990; Baral et al. 2021; Du et al. 2022; Zhang et al. 2014; Chen et al. 2024; Veen et al. 2025) and effectively captures maintenance dynamics (**Fig. 3A,G**). However, because it integrates curvature along the organ, the hook angle masks underlying growth patterns. Modeling demonstrates that any given hook angle, normal or exaggerated, can hypothetically arise from distinct configurations of Δ and 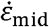, reflecting fundamentally different mechanisms (**Fig. S6**). Our framework resolves this ambiguity by analysing cell growth patterns. We next used it to test whether auxin distribution and signaling are sufficient to account for the coordinated growth dynamics underlying hook robustness.

### Auxin-Mediated Growth Repression Is Apically Restricted During Hook Maintenance

The apparent mismatch between the biphasic growth pattern and the unilateral auxin distribution indicated by DR5 during maintenance (**Fig. 1**) prompted us to examine earlier growth dynamics. We found that the maintenance pattern emerges progressively during formation (**Fig. 4A–B; Fig. S7**). Early on (0 h, ∼50–70°), Δ is positive along the entire hook, but a basal negative Δ region subsequently appears and advances toward the apex (3–9 h) as the hook acquires a self-similar shape. Thus, uniform auxin-mediated growth inhibition along the inner side applies only at early stages.

**Figure 4.**
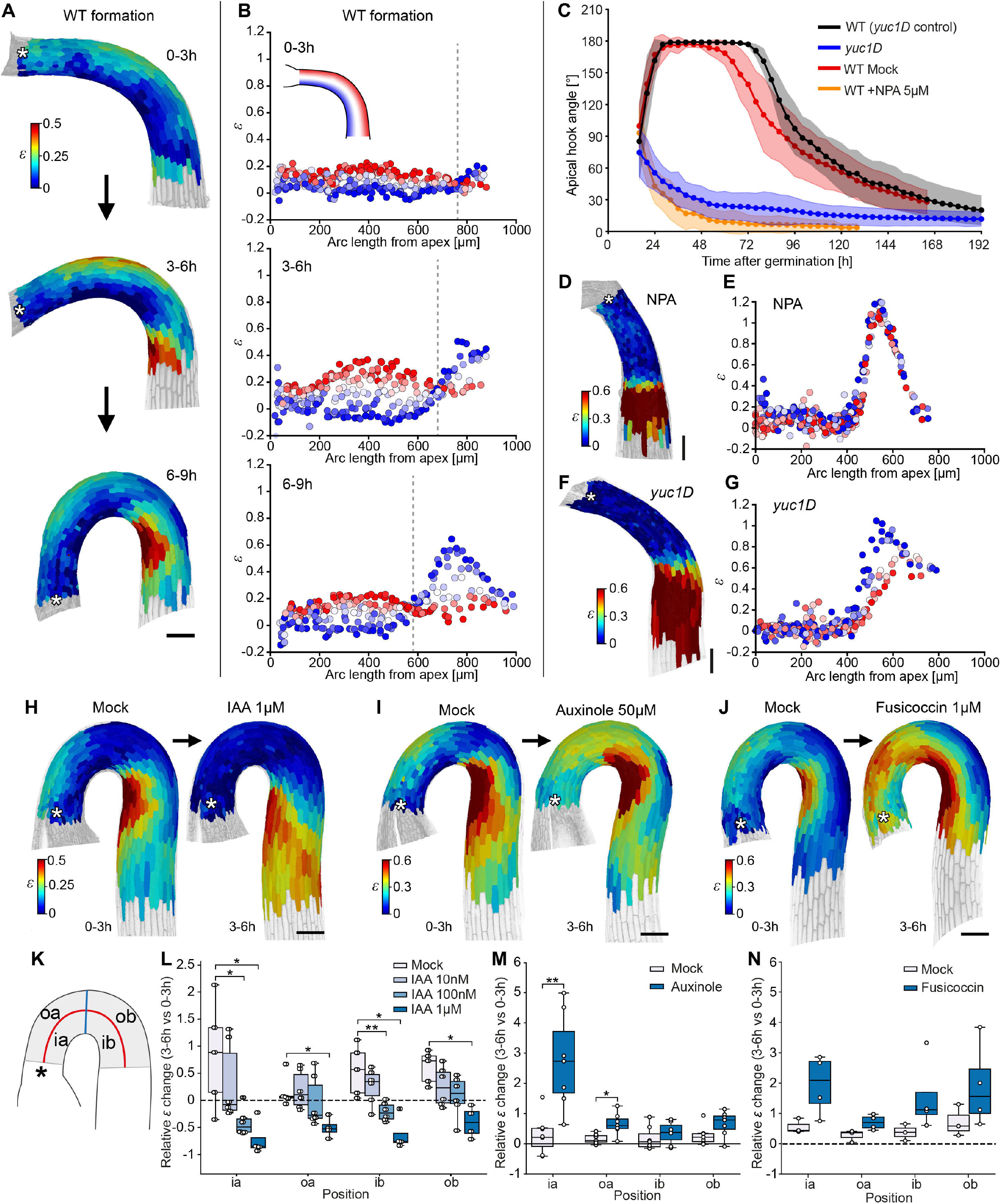
Growth Profiling and Auxin Perturbations Demonstrate Apically Confined Auxin Growth Repression During Hook Maintenance. **(A)** Heatmaps of strain (*ε*) in WT during apical hook formation, shown for three sequential 3 h timelapse intervals. **(B)** Individual cell strain patterns in WT corresponding to the intervals of the individual in **(A)**. Each point represents a single cell, color-coded by its position across the hypocotyl (blue = inner side, red = outer side). Inset: schematic of the apical hook illustrating the color-coding. Vertical dotted lines indicate Δ= 0. Additional biological replicates are shown in **Fig. S8** (*n* = 3). **(C)** Apical hook angle in *yuc1D* and WT ± NPA 5 μM treatment. Measurements were performed at 4 h intervals. n ≥ 8. Shaded areas indicate mean ±SD. **(D)** Heatmap of strain in WT germinated on NPA 5 μM during formation phase. **(E)** Individual cell strain patterns in WT germinated on NPA 5 μM during formation phase, corresponding to **(D)**, with individual dot color-coded as in **(B)**. Additional biological replicates are shown in **Fig. S9** (*n* = 3). **(F)** Heatmap of strain in *yuc1D* during formation phase. **(G)** Cell strain patterns in *yuc1D* during formation, corresponding to **(F)**, with individual dots color coded as in **(B)**. Additional biological replicates are shown in **Fig. S10** (*n* = 3). **(H)** Heatmaps of strain in WT upon sequential treatments of Mock (0-3 h interval) and IAA 1 μM (3–6 h interval) at onset of maintenance (*t* = 0 h). **(I)** Heatmaps of strain in WT upon sequential treatments of Mock (0–3 h interval) and Auxinole 50μM (3–6 h interval) at onset of maintenance (*t* = 0 h). **(J)** Heatmaps of strain in WT upon sequential treatments of Mock (0–3 h interval) and Fusicoccin 1μM (3–6 h interval) at onset of maintenance (*t* = 0 h). **(K)** Schematic representation of the apical hook, indicating regions examined in L-N; oa = outer apical, ob = outer basal, ia = inner apical and ib = inner basal. Apical and basal regions are divided by location of maximal curvature (blue line), each extending 200 μm in either direction. Inner and outer regions are divided by organ centerline (red line). **(L-N)** Relative change in strain between the 0–3 h and 3–6 h intervals in different regions of the apical hook. Experiments were initiated at the onset of hook maintenance (***t*** = oh). Seedlings were kept under mock conditions during the first interval (0–3 h). At 3 h, seedlings were transferred to fresh mock medium or to medium containing the chemical treatment. The relative strain change was calculated as 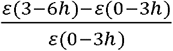. Boxplots show the median (line), interquartile range (box), whiskers (1.5 × IQR), and individual replicates (white circles). Asterisks indicate statistical significance (*p < 0.05, p < 0.005; two-sided Mann-Whitney U rank test). Regions examined as outlined in **(K)**. **(L)** IAA treatment (10nM, 100nM, 1μM). Individual replicate cell strain patterns are shown in **Fig. S11-14** (*n* = 5 mock, 6 10 nM, 6 100 nM, 4 1 μM). **(M)** Auxinole treatment (50 μM). Individual replicate cell strain patterns are shown in **Fig. S15-16** (*n* = 7 mock, 7 50 μM). **(N)** Fusicoccin treatment (1μM). Individual replicate cell strain patterns are shown in **Fig. S17** (*n* = 3 mock, 4 1 μM). All scale bars, 100 μm. Asterisks in A, D, F, H-K indicate hypocotyl apex.

To test whether auxin asymmetry is required to establish the biphasic pattern, we disrupted auxin distribution prior to hook formation, using complementary approaches (**Fig. 4C**). Inhibition of auxin polar transport with N-1-naphthylphthalamic acid (NPA) (Abas et al. 2021) prevented hook formation, as previously reported (Zádníková et al. 2010). The auxin-overproducing *yuc1D* mutant also failed to form a hook (Zhao et al. 2001; King et al. 1995; Jonsson et al. 2021), while displaying uniformly high levels of auxin signaling response (**Fig. S8**). Despite contrasting effects on auxin distribution, NPA-treated WT and *yuc1D* show strikingly similar growth patterns, with negligible inner–outer asymmetry and a basally increasing midline elongation rate 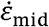 (**Fig. 4D–G; Fig. S9–S10**). These results indicate that, while auxin asymmetry drives the differential growth needed for hook formation, the axial elongation profile 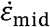 observed during maintenance emerges independently from auxin and from the hook shape.

Perturbing hook formation via genetics or long-term treatments prevents the onset of maintenance, obscuring auxin’s role in self-similar growth. Therefore, we applied short-term (3 h) treatments (IAA, auxinole, fusicoccin) during the maintenance, to test the role of auxin in the biphasic growth pattern (**Fig. 4H–N**). Hook growth post-treatment (3–6 h) was quantified relative to pre-treatment maintenance dynamics (0–3 h). The hook was partitioned into four domains (inner/outer × apical/basal) based on the midline and maximal curvature (**Fig. 4K**).

To test whether regional growth patterns reflect differences in auxin responsiveness, we performed dose–response experiments with exogenous IAA (10 nM–1 μM). Auxin reduced growth uniformly across the hook in a dose-dependent manner (**Fig. 4H, L; Fig. S11–S14**), indicating that spatial differences in responsiveness cannot explain the inner-side transition from slow to fast growth.

Blocking auxin perception with auxinole (Hayashi et al. 2012) caused strong growth de-repression, most pronounced in the inner apical region (**Fig. 4I, M; Fig. S15–S16**), indicating that endogenous auxin-mediated repression is localized to this domain. Fusicoccin, which maximizes acid-growth–driven expansion (Kiriyama et al. 2024; Lado et al. 1973; Adamowski et al. 2019; Fendrych et al. 2016), produced a similar de-repression pattern (**Fig. 4J, N; Fig. S17**). Therefore, auxin likely regulates hook development primarily by suppressing acid-growth machinery.

Given that the precise auxin distribution within the hook remains unresolved, it is commonly assumed that polar transport generates uniform accumulation along the inner side, repressing growth. Our results show that this model holds only during early formation. However, during maintenance, growth inhibition is strongest in the inner apical region, suggesting spatially restricted auxin activity. We therefore directly examined auxin distribution and signaling across hook domains.

### Auxin Accumulation and Signaling are Spatially Confined to the Inner Apical Domain During Maintenance

Our perturbation experiments show that auxin represses acid-mediated growth in the inner apical region, yet the rapid elongation observed basally remains unexplained. High-resolution tracking reveals a sharp, spatially fixed transition along the inner side, where cells switch from slow to rapid elongation as they move basally, increasing growth by over 700% within 3 h and reaching among the highest rates reported in the hypocotyl (Alimchandani et al. 2026) (**Fig 5A**). Although a strong basal acceleration of growth along the midline is present regardless of auxin distribution (**Fig. 4D,F**), we asked how auxin patterning relates to growth along the inner side.

**Figure 5.**
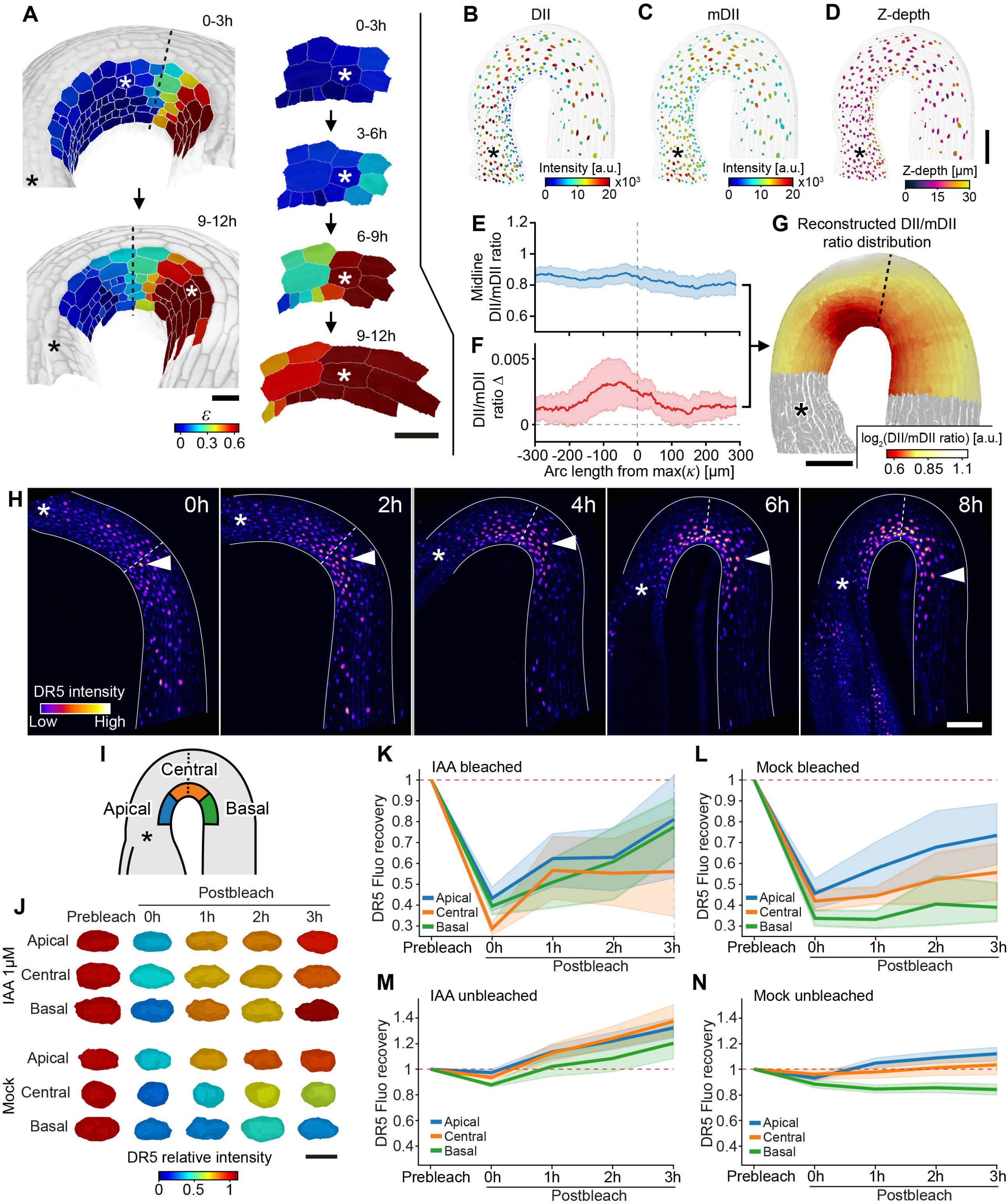
R2D2 and DR5 FRAP Reveal Apical-Restricted Auxin Abundance and Signaling During Maintenance. **(A)** During hook maintenance, a fast switch in cell growth on the inner side occurs at the location of maximal curvature and remains static while cells move through the hook structure. Left panel depicts overall cell positions progression over 9 h. Right panel depicts the same cells over 4 sequential 3 h intervals. Black asterisks indicate hypocotyl apex. White asterisks mark the same cell over time; its elongation rate increased from *ε* = 0.069 (3–6 h) to *ε* = 0.59 (6–9 h), corresponding to an ∼8.5-fold increase over 3 h. Dotted lines indicate position of maximal curvature. Scale bars, 40 μm. **(B–G)** Quantification of auxin distribution using the R2D2 ratiometric reporter at the onset of hook maintenance in WT. **(B)** Heatmap of DII nuclear signal projected onto segmented nuclei. **(C)** Heatmap of mDII nuclear signal projected onto segmented nuclei. **(D)** Heatmap of nuclear Z-depth projected onto segmented nuclei, used to correct for signal attenuation with imaging depth. Scale bar for **(B-D)** 100, μm. **(E)** Average DII/mDII ratio along the longitudinal axis of the hypocotyl after correction for nuclear depth. *n* = 15. **(F)** Lateral (inner/outer) gradient of the DII/mDII ratio along the longitudinal axis of the hypocotyl after correction for nuclear depth. *n* = 15. **(G)** Reconstructed DII/mDII ratio distribution projected onto the epidermal mesh surface, combining the longitudinal mean signal and lateral asymmetry. Values are displayed as log_2_-transformed DII/mDII ratios. The dotted line indicates the position of maximal curvature. Scale bar, 100 μm. **(H)** Confocal timelapse series of DR5::Venus during apical hook formation. DR5 distribution during maintenance indicates auxin accumulation on the inner side both in the apical and basal regions. However, examination of DR5 from formation to early maintenance shows that DR5::Venus-expressing cells quickly move from the apical to basal region as the hook elongates. White arrowheads indicate the same nucleus over time. Dotted lines indicate the point of maximal curvature. Asterisks indicate hypocotyl apex. Solid lines indicate hypocotyl outline. *n* = 6. Scale bar, 100 μm. **(I)** Schematic representation of the apical hook indicating the regions examined in DR5::Venus FRAP experiments; Central (within ±100 μm of point of maximal curvature), apical (<-100 μm), and basal (>100 μm). Dotted line indicates point of maximal curvature. Asterisk indicates hypocotyl apex. **(J)** Heatmaps of nuclei showing relative DR5 levels in WT from onset of hook maintenance, normalized to prebleach levels, during prebleach and sequential postbleach timepoints (0 min, 1 h, 2 h and 3 h) in regions as outlined in **(F)**. Scale bar, 4 μm. (For representative confocal images, see **Fig. S19a**, for DR5 analysis pipeline, see **Fig.S19b**). **(K–N)** DR5::Venus relative fluorescence recovery kinetics upon IAA 1μM **(K, M)** and mock **(L, N)** in either bleached cells **(K–L)** or unbleached neighbors **(M–N)**. After bleaching upon IAA, cells in all regions exhibited uniformly strong DR5 recovery **(K)**, while upon mock, recovery was strongest in the apical region and slowest in basal cells **(L)**. DR5 fluorescence intensity kinetics in unbleached neighboring nuclei exhibited similar trends as bleached nuclei, with unbleached nuclei in IAA-treated seedlings increased their fluorescence in all regions **(M)** while upon mock treatment, nuclei mildly increased fluorescence in the apical region while nuclei in the basal region reduced their fluorescence over time **(N)**. *n* = 5 Mock, 4 IAA.

To map auxin distribution, we used the R2D2 ratiometric sensor, in which auxin-dependent degradation of DII is normalized by a stable mDII reference (Brunoud et al. 2012; Janacek et al. 2024) (**Fig. 5B–G**). We quantified nuclear DII/mDII ratio using 3D segmentation, and extracted their longitudinal and lateral gradients to relate auxin distribution to the growth profiles 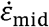 and Δ (**Fig. S18**). To correct for depth-dependent signal attenuation (Chudakov et al. 2010) we estimated the optical path length from each nucleus to the organ surface (**Fig. 5D**) and applied a weighted least-squares regression, prioritising nuclei with minimal signal distortion while extracting the gradients.

R2D2 analysis revealed a spatially restricted auxin pattern. Overall levels were uniform along the axis (Fig. 5E), but the lateral DII/mDII gradient peaked sharply in the apical region, upstream of maximal curvature (**Fig. 5F**). Reconstructed auxin levels were highest in the inner apical domain, intermediate basally, and lowest on the outer apical side (**Fig. 5G**). Auxin accumulation is therefore concentrated to the inner apical region during maintenance, while more evenly distributed between the inner and outer sides in the basal region. Previous DR5 and GUS reporter studies consistently suggest auxin accumulation along the inner side of the hook, including the basal region (H. Li et al. 2004; Gallego-Bartolomé et al. 2011; Zádníková et al. 2010). We hypothesized that this discrepancy with R2D2 arises from temporal “smearing” of transcriptional reporters. Signals from DR5::GUS and DR5::Venus lag behind auxin input due to transcription, translation, and chromophore maturation. Moreover, their proteins are highly stable—GUS can persist for >50 h, and Venus for several hours after auxin levels decline (X. Li et al. 1998; Jedličková et al. 2022; de Ruijter et al. 2003; Brunoud et al. 2012; Jefferson et al. 1987). We tracked DR5::Venus dynamics in individual nuclei from formation to early maintenance. During formation, DR5 marked auxin signaling all along the inner side (**Fig. 5H**, 0 h, 2 h). However, at the onset of maintenance, part of the basal DR5 signal originated from cells displaced from the apical region (**Fig. 5H**, arrow), indicating that basal signal might reflect reporter persistence to prior auxin exposure.

To test whether basal DR5 signal reflects active auxin signaling or residual reporter, we quantified DR5::Venus turnover using FRAP (Fluorescence Recovery After Photobleaching) (**Fig. 5I–N; Fig. S19**). Under exogenous IAA, photobleached nuclei recovered similarly across apical, central, and basal regions (**Fig. 5J–K**), indicating comparable signaling capacity. Under mock conditions however, recovery was strongly region-specific, fastest apically and slowest basally (**Fig. 5J,L**). In unbleached nuclei, IAA increased DR5 signal across all regions (**Fig. 5M**), whereas basal nuclei lost fluorescence over time under mock conditions, consistent with slow reporter decay (**Fig. 5N**). Together, these results uncover an apical-to-basal gradient of auxin signaling along the inner side, in agreement with the distribution revealed by R2D2, and show that static DR5 patterns are blurred by reporter persistence and cell displacement along the hook.

Auxin perturbations, R2D2, and DR5 FRAP consistently show that during maintenance, auxin activity peaks in the inner apical region and declines basally. Thus, auxin drives the positive apical Δ underlying bending but cannot explain the basal negative Δ required for straightening. To identify this mechanism, we turned to macroscopic rod models.

### Curvature-Dependent Feedback Confers Robustness to Shape Perturbations

To investigate hook dynamics beyond auxin, we next asked whether curvature itself could provide the missing regulatory signal. In rod models, this is captured by autotropic feedback, which links differential growth (Δ_auto_) to local curvature (**Fig. 6A**). Highly curved regions are locally straightened by promoting growth on the inner side (**Fig. 6B–C**). Although its molecular basis remains unclear, such curvature-dependent feedback is required to reproduce realistic bending dynamics (Bastien et al. 2013, 2014; Porat et al. 2024). In our model, Δ_auto_ offers a parsimonious explanation for the basal negative Δ.

**Figure 6.**
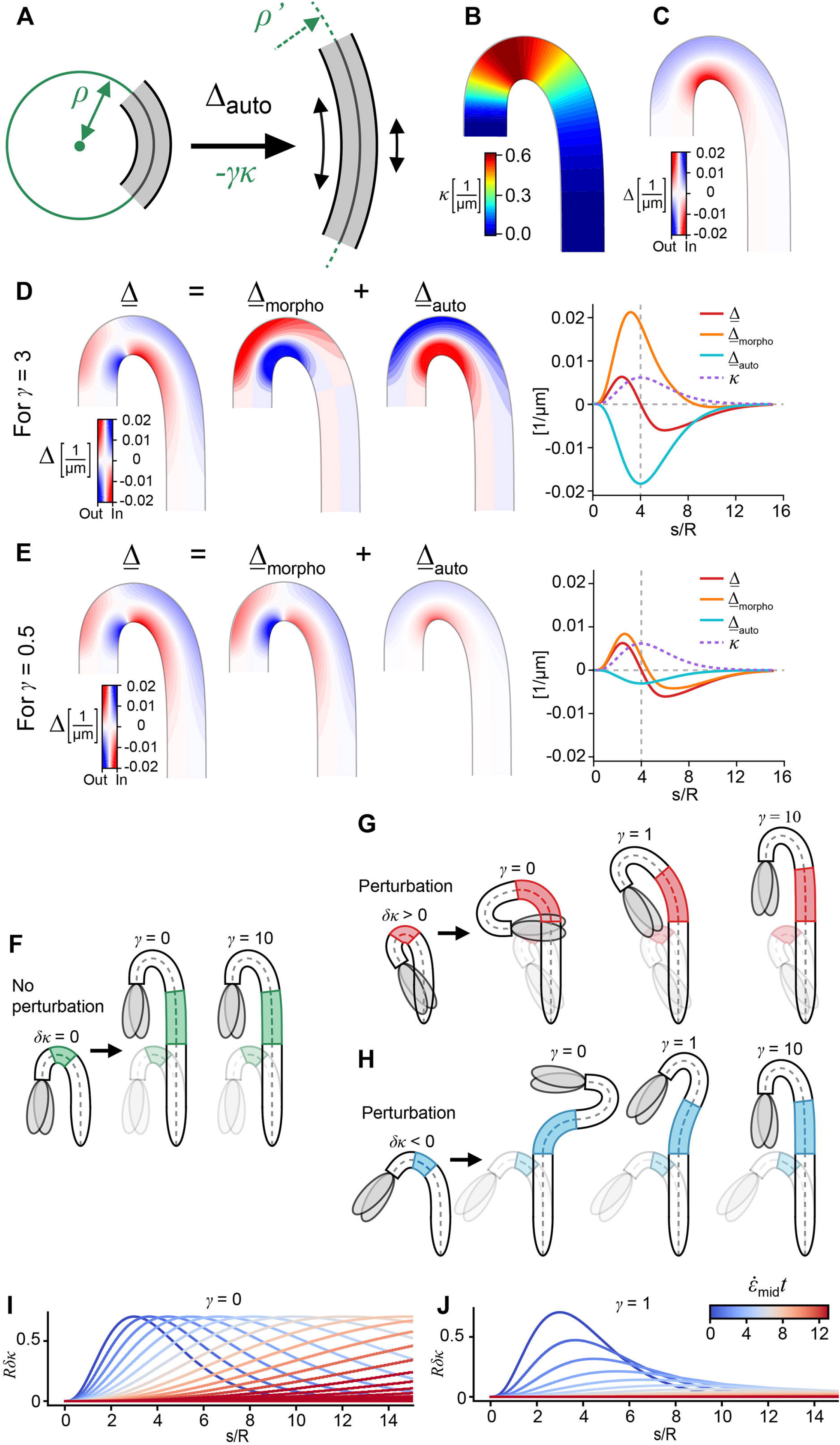
Modeling Suggests Autotropic Feedback May Complement Auxin and Confer Dynamical Stability. **(A–C)** Straightening (decrease in curvature) is controlled by Δ_auto_, corresponding to autotropism. **(A)** The higher the curvature, the faster autotropism locally straightens the material segment via Δ_auto_ = -*γκ*, where *γ* is the autotropic sensitivity (*γ* ≥ 0) and *κ* is the midline curvature. In the illustration, *ρ*= 1/*κ* is the initial radius of curvature and *ρ ′* is the radius of curvature after incremental growth. **(B)** Heatmap representing *κ* of the model hook. **(C)** Example for Δ_auto_ with *γ* = 1 on the model hook. In heatmaps depicting differential growth Δ and its additive decompositions, the local color represents 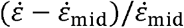 (as in **Fig. 2H**). The color intensity indicates the magnitude of Δ (positive or negative) as depicted by the colorbar. **(D–E)** The self similar differential growth Δ can be decomposed additively into two terms: Δ = Δ_morpho_ + Δ_auto_. The model profile of Δ shown corresponds to a self-similar growth pattern set by the given model *κ* and 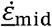. Using Δ_morpho_ = Δ + *γκ* then allows to decomposed Δ into pairs of curves Δ_morpho_, Δ_auto_} which depend on the value of *γ*. Left: The decomposition presented on the model hook; Right: The decomposition presented as a function of arc length *s* normalized by the cross section radius *R*. **(D)** For *γ*= 3, Δ_morpho_ is mostly positive and competes with strong autotropism in the apical part. **(E)** For *γ*= 1.5, Δ_morpho_ has both a positive and a negative part, meaning that the morphogen controls both bending and straightening. **(F–J)** To study the developmental robustness predicted by our model, we introduce a small initial curvature perturbation *δκ* over a small material segment at *t* = 0 and observe the resulting shape after growth. Assuming a constant relative growth rate 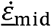 allows us to solve the spatiotemporal dynamics analytically. **(F)** The unperturbed shape dynamics is self-similar and does not depend on *γ*. **(G–H)** The shape dynamics after the perturbation *δκ* > 0 **(G)** or *δκ* < 0 **(H)** depends on *γ*: For low autotropism 0 ≤ *γ* < 1, the perturbation is amplified by the elongation of the perturbed segment; For *γ*= 1, the decay of *δκ* within the perturbed segment is perfectly balanced with the elongation of the segment, resulting in a constant excess angle and inclination of the model cotyledons; For high autotropism *γ*> 1, the perturbation decays at a rate which increases with *γ*. **(I-J)** The spatiotemporal dynamics of a perturbation *δκ* displayed as the progression of its spatial profiles at different times. Spatially, arc length *s* is normalized by the cross section radius *R*, and temporally, time *t* is normalized by the constant relative growth rate 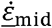. **(I)** For *γ*= 0, the perturbation spreads along the hook undisturbed following a one-way wave equation. **(J)** For *γ*= 1, the perturbation decays while spreading.

To test this hypothesis, we decomposed the lateral growth gradient into bending and straightening components (**Fig. 6D–E**):

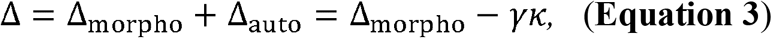

where *γ* is the autotropic feedback sensitivity. We assume that the bending contribution

Δ_morpho_ is set by auxin patterning, so the measured Δ reflects the sum of auxin-driven bending and curvature-dependent straightening. We therefore inferred the stationary field Δ_morpho_ from experimental Δ and *κ* maintenance profiles, across *γ* values. Δ_morpho_ profile depends strongly on r, which tunes the competition between the fields. High *γ* leads to excessive straightening that can only be compensated by unrealistically strong apical bending (Δ_morpho_), effectively implying negative growth on the inner side (**Fig. 6D**). Conversely, low *γ* fails to generate the observed basal negative Δ via autotropism only (**Fig. 6E**). Together, these constraints bound *γ* to low values apically (*γ* ≲ 3) and higher values basally (*γ* ≳ 3) (**Fig. S20**).

We next asked how the model explains the remarkable reproducibility of hook shape during maintenance (**Fig. 6F–J**). Since growth processes are inherently noisy (Hong et al. 2016; Raj and van Oudenaarden 2008), we can expect small fluctuations in Δ_morpho_, locally distorting curvature. As the organ elongates, local curvature defects (*δκ*) are moved along the hook and stretched, similarly to other fluctuations (Fruleux et al. 2024), while their amplitude evolves according to the strength of autotropic feedback *γ* (see SI):

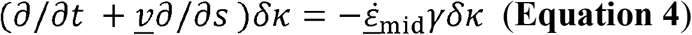

Without feedback (*γ* = 0), small perturbations are passively transported, while growth amplifies organ distortion over time (**Fig. 6G–I, Movie S9–S10**). With feedback (*γ* > 0), perturbations are damped and the original shape is restored (**Fig. 6G–H, J**). Complete stabilization occurs for *γ* > 1, consistent with theoretical predictions for posture control (Bastien et al. 2014; Oliveri et al. 2024). Given the drastic effect of small perturbations in the absence of feedback, observation of the maintenance phase suggests that *γ* > 1 is fulfilled.

Together, these results suggest that curvature-dependent feedback explains the negative Δ during maintenance. In this framework, auxin drives the positive Δ causing apical bending, while autotropism generates curvature-dependent straightening that stabilizes organ shape against growth noise. A sufficiently large autotropic sensitivity *γ* is required for robust maintenance and to infer a Δ_morpho_ pattern consistent with auxin patterns. However, large *γ* values also predict strong competition between bending and straightening in the apical region. To test model predictions, we next sought to quantify curvature-dependent growth responses *in vivo*.

### A Gradient of Autotropic Sensitivity Spatially Separates Curvature Feedback from Auxin-Driven Bending

In order to isolate curvature-dependent growth, we suppressed auxin-driven bending (Δ_morpho_) experimentally. Inhibiting auxin transport with NPA (5 µM) rapidly abolished the positive apical Δ within 3 h, while the basal negative Δ persisted (**Fig. 7A,B**), leading to hook opening upon prolonged treatment (**Fig. 7C**). Co-treatment with NPA and uniform auxin (NAA, 500 nM) produced the same response (**Fig. S21**), indicating that straightening does not depend on auxin gradients.

**Figure 7.**
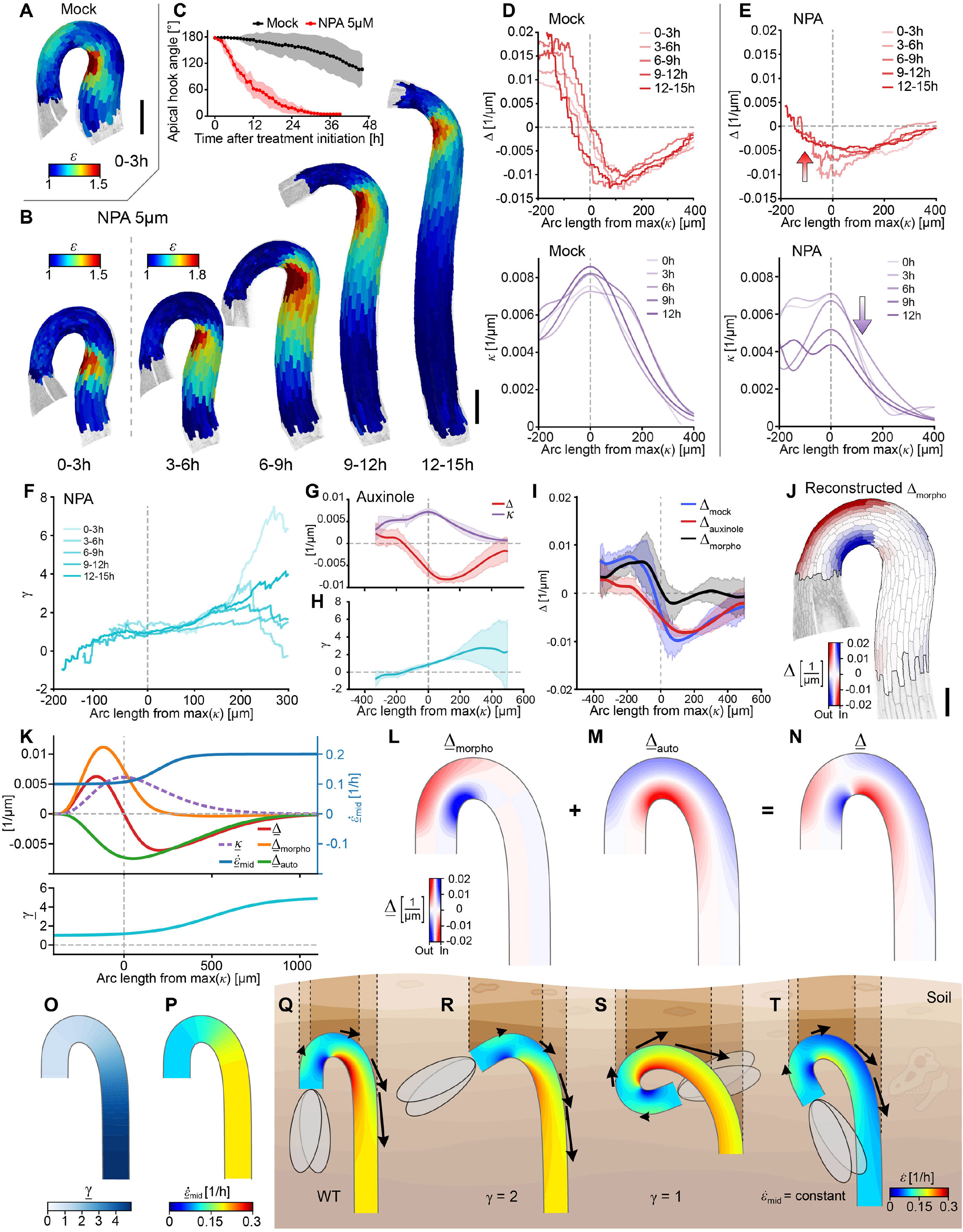
Experimental Decomposition of Growth Asymmetry Reveals Spatially Patterned Autotropism Optimizing Hook Shape. **(A)** Growth of WT hook during maintenance, under mock conditions. *n* = 2. Scale bar, 200 μm. **(B)** Growth of WT hook under long-term treatment with NPA (5 uM) initiated after the onset of hook maintenance (treatment start = 0 h). *n* = 3. Scale bar, 200 μm. **(C)** Hook angle measured from the onset of hook maintenance phase (0 h). At 0 h, seedlings were transferred to either mock and NPA 5uM conditions. *n* ≥ 8. **(D)** Profiles of Δ and *κ* over time during maintenance under mock conditions. All positions are aligned according to the maximal curvature. **(E)** Profile of Δ and *κ* over time during long-term NPA treatment for the sample shown in **(B)**. The amplitudes of Δ and *κ* decrease over time (arrows). **(F)** Inference of the autotropic sensitivity *γ* from the profiles of Δ and *κ* shown in **(E)**. **(G)** Profiles of Δ and *κ* after auxinole treatments (see Fig. 4). *n* = 7. **(H)** Inference of *γ* from Auxinole treatment. **(I)** Inference of Δ_morpho_ from auxinole treatment via Δ_morpho_ = Δ_mock_ - Δ_auxinole_. **(J)** Heatmap showing the Δ_morpho_ gradient on the actual hook geometry. In heatmaps depicting differential growth Δ and its additive decompositions, the local color represents 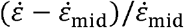 (as in **Fig.2H**). The color intensity indicates the magnitude of Δ (positive or negative) as depicted by the colorbar. Scale bar, 100 μm. **(K)** Model of hook maintenance, using idealized profiles of 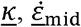 and *γ* as inputs. Δ_morpho_ is obtained as a model output. **(L)** Heatmap of the idealized <u>Δ</u>_morpho_ gradient obtained by the model. **(M)** Heatmap of the <u>Δ</u>_auto_ gradient used in the model. **(N)** Heatmap of the <u>Δ</u>, the sum of <u>Δ</u>_morpho_ and <u>Δ</u>_auto_. **(O)** Heatmap of the *<u>γ</u>* profile imposed in the maintenance model. **(P)** Heatmap of the midline strain profile 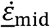 imposed in the maintenance model. **(Q)** Stationary growth pattern and shape correspond to the normal parameter profiles of maintenance. The resulting hook shape allows a small frontal contact with soil (in brown) when pushing upwards. Arrows indicate displacement speed *v* along the hook. Dotted vertical lines outline soil-contact zones: outer = full vertical hook–soil interface; inner = high-angle contact zone (45°–90° relative to vertical). **(R–T)** Stationary growth patterns and shapes obtained by flattening one graded profile, of either *γ* or 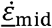, with respect to our maintenance mode. 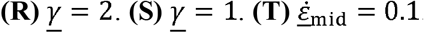.

To confirm that the treatment isolated the active Δ_auto_ by suppressing Δ_morpho_, we analyzed Δ profiles over time. During normal maintenance, the stable peaks of negative Δ and curvature are spatially offset (**Fig. 7D**). Under NPA, these peaks coincide, shift apically, and decay together (**Fig. 7E; Fig. S22–S23**), as expected for Δ_auto_, which scales with curvature. With Δ ≈ Δ_auto_, we estimated the autotropic sensitivity as *γ* = −Δ/*κ*. Profiles of *γ* were stable over time and revealed a pronounced gradient, increasing from *γ* ∼ 1 at the apex to *γ* ∼ 5 at the base (**Fig. 7F; Fig. S23**). These values satisfy the stability condition (*γ*> 1) (**Fig. 6I–J**) and are consistent with prior estimates in roots (*γ* ≈ 1.5; Porat et al., 2024). While spatial variation in autotropic sensitivity were previously unknown, these findings are consistent with our predicted bounds for apical (*γ* ≲ 3) and basal (*γ* ≳3) regions (**Fig. S20**). Auxinole treatment yielded comparable *γ* profiles (**Fig. 7G–H**), confirming that this gradient is auxin-independent. Together, these results establish a spatially graded, auxin-independent curvature feedback along the organ.

We next sought to isolate the bending component, Δ_morpho_. Although not directly measurable, it can be inferred by comparing Δ profiles before and after auxinole treatment (**Fig. 7I**). The inferred Δ_morpho_ closely matches the asymmetric auxin distribution measured by R2D2 (**Fig. 5G**) and the signaling pattern revealed by DR5 FRAP (**Fig. 5L**), supporting a causal link between auxin activity and bending.

To test consistency, we integrated all experimental profiles into a single model. Using a profile of *γ* increasing along the hook, we computed Δ_auto_ from the measured curvature during maintenance (**Fig. 7K–Q**), and combined it with Eqs. 2–3 to reconstruct Δ_morpho_. The resulting profile Δ_morpho_ closely matched the experimentally inferred one (**Fig. 7J,L**). Importantly, the increase in autotropic sensitivity along the hook minimizes interference between bending and straightening: low *γ* in the apex permits auxin-driven bending, while high basal *γ* ensures robust stabilization.

Together, these results show that hook maintenance emerges from the coordinated action of three spatial growth fields: auxin-driven Δ_morpho_ in the apex, curvature-dependent Δ_auto_ set by a gradient of *γ* independent of auxin (**Fig. 7O**), and 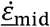 controlling elongation independently of auxin (**Fig. 7P**).

Lastly, we asked whether spatial gradients of autotropic sensitivity and axial growth fine-tune hook shape for its protective function (**Fig. 7O–T**). During germination, rapid hypocotyl elongation pushes the hook through the soil. Optimal soil penetration requires to reduce the hook mechanical resistance, hence minimizing the projected frontal area: sharp geometries concentrate stress at the tip and limit distributed contact with the surrounding medium.

Using the inferred Δ_morpho_ profile, we compared WT geometry with shapes generated under uniform *γ* and 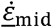 profiles. The experimental profiles reproduced WT dynamics while minimizing frontal contact (**Fig. 7Q**). By contrast, uniformly high *γ* produced a shallow hook with increased frontal area, whereas low *γ* exaggerated curvature (**Fig. 7R–S**), both predicted to elevate compressive loading and promote buckling. Finally, imposing a constant 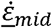 equal to the apical WT value slowed cell displacement, prolonging exposure to compressive stress (**Fig. 7T**). Together, spatial profiles of *γ* and 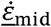 optimize hook geometry during emergence by maintaining robust curvature while minimizing both the magnitude and duration of mechanical stress experienced by cells passing through the frontal contact zone. As all cells in the hook ultimately contribute to hypocotyl elongation, this protection is likely critical for seedling survival.

## Discussion

Stable curvature in the growing *Arabidopsis thaliana* apical hook emerges from the coupling of local morphogen activity and global geometric feedback. Auxin asymmetry drives bending but is insufficient to maintain shape. Instead, curvature is stabilized by a closed-loop system combining apical auxin-mediated growth repression with basal straightening mediated by graded autotropic feedback. This division of labor separates pattern generation from structural stabilization, enabling robust shape maintenance during growth and optimizing the hook for efficient soil penetration. This framework raises two key mechanistic questions.

What confines auxin activity to the inner apical domain during maintenance? PIN-dependent auxin transport is sensitive to organ geometry, becoming disorganized in hookless conditions and restored by mechanical bending (Zádníková et al. 2010; Baral et al. 2021). We observed the auxin maximum is increasingly restricted spatially to the apical inner side during formation, suggesting that curvature may bias auxin flux within the bending zone. This supports a feedback loop in which auxin transport drives curvature, while curvature in turn stabilizes the transport configuration that sustains the apical auxin domain, reminiscent of the mechanism proposed in the meristem (Heisler et al. 2010).

What generates the graded autotropic feedback that limits bending? One possibility is a dedicated curvature-sensing system, as proposed in proprioceptive frameworks (Moulia et al. 2019), although no molecular correlate has been identified in the hook. Alternatively, autotropism may arise from mechanosensitive pathways, in which curvature-induced stress patterns are transduced into growth regulation through calcium signaling, cell wall integrity pathways, or cytoskeletal reorganization (Moulia et al. 2021; Tran et al. 2017; Park and Shin 2022; Feng et al. 2018; Bacete and Hamann 2020, Chebli et al. 2021; Walia et al. 2024). Finally, straightening may emerge from passive mechanical properties without explicit sensing (Porat et al. 2025). Discriminating among these mechanisms will require integrating mechanical perturbations, genetics, and quantitative modeling.

Together, our results define a general principle: morphogen-driven local growth asymmetry generates global form, while geometry-dependent feedback stabilizes the shape via a superimposed growth pattern. This dual architecture provides a robust strategy for shaping growing organs.

## Materials and Methods

*Arabidopsis thaliana* lines used in this study included *yuc1D* (Zhao et al. 2001), *ctr1-1* (Kieber et al. 1993), *axr2-1* (Timpte et al. 1994), pUBQ10::PM-tdTomato (Segonzac et al. 2012), DR5::Venus (Heisler et al. 2005), UBQ10::H2B-mCherry (Marquès-Bueno et al. 2016), R2D2 (Janacek et al. 2024).

Chemicals used in this study included Naphthylphthalamic acid (NPA) (Duchefa, N0926), Indole-3-acetic acid (IAA) (Sigma-Aldrich, 1003530010), 1-Naphthaleneacetic acid (NAA) (Sigma-Aldrich, N0640), Auxinole (MedChemExpress, HY-111444), and Fusicoccin (MedChemExpress, HY-122815).

### Growth conditions

Seedlings were grown on square Petri dishes supplied with 1/2 MS (2.2 g/l Murashige & Skoog nutrient mix (Duchefa), 0.8% (w/v) plant agar (Duchefa), 0.5% (w/v) sucrose, 2.5 mM 2-Morpholinoethanesulfonic acid (MES) (Sigma-Aldrich) buffered at pH 5.8 with KOH. For all experiments, seeds were stratified for 2 days at 4°C, exposed to 6h of light treatment and then grown vertically in darkness at 21°C for the required time length.

### Kinematic analysis of hook angle and hypocotyl length

For time-lapse analysis of apical hook development and hypocotyl length, seedlings grown in darkness at 21°C on vertical Petri dishes were imaged at 1 h, 3 h, or 4 h intervals using a Canon EOS D50 camera with a Godox Thinklite speedlite TT520 flash equipped with a green filter and diffusion sheets. For pharmacological treatments, seedlings were germinated on agar plates supplemented with the respective chemical or solvent control. For hook maintenance transfer experiments, seedlings were grown on control plates until early hook maintenance, and then transferred to agar plates supplemented with the respective chemical or solvent control. Hook angle was quantified in Fiji (Schindelin et al. 2012) using the Angle tool, measuring the angle formed between the hypocotyl axes above and below the hook (see inset of **Fig. 1B**). Hypocotyl length was measured in Fiji using the Segmented Line tool. ≥8 seedlings analyzed for all experiments.

### Confocal microscopy

Confocal laser scanning microscopy (CLSM) was performed using either a Zeiss LSM800 or LSM880 microscope, as specified in each subsection. Seedlings were typically mounted on agar blocks without coverslip, unless otherwise stated. Imaging was carried out under dark conditions by covering the microscope with a textile blackout curtain. To locate seedlings through the oculars, a dim external green light source was used instead of the microscope’s transmitted light illumination. Seedlings were maintained vertically on agar blocks between imaging intervals inside custom 3D-printed dark chambers. Images were acquired using Zeiss ZEN software. All experiments were performed at room temperature.

### Time-lapse imaging of cell elongation

Time-lapse imaging by CLSM was performed on a Zeiss LSM800 with a 10×/0.45 NA Plan-Apochromat dry objective. Seedlings were mounted on agar blocks without coverslip and imaged using tdTomato as a fluorescent marker (excitation 561 nm; emission collected between 580–620 nm). Laser intensity was adjusted dynamically across the z-stack (1–15%) to compensate for signal attenuation at deeper focal planes. Acquisition cubic voxel size ranged 0.64 µm and 1 µm, but was kept constant within individual experiments. Scans were acquired in bidirectional mode without averaging. Between imaging time points, seedlings were incubated vertically on agar blocks in 3D-printed dark chambers.

For control imaging conditions, seedlings germinated on control medium were transferred to control agar blocks either at the stage of hook formation (∼16 h after germination) or at onset of hook maintenance (∼24 h after germination). For experiments where seedlings were germinated on chemical treatment medium, they were transferred at ∼24 h after germination to agar blocks containing the same chemical or solvent control. For experiments where treatment was induced at the onset of hook maintenance, seedlings were germinated on control medium, transferred to control agar blocks at ∼24 h, and subsequently shifted to chemical treatment agar blocks at a defined time point during the time-lapse. In these induction experiments, a droplet of liquid medium containing the respective chemical or solvent control was applied at the onset of treatment and replenished after each imaging interval.

### R2D2 imaging

R2D2 imaging was performed on a Zeiss LSM880 using a 40×/1.2 NA water-immersion objective. Seedlings at onset of hook maintenance (∼24 h after germination) were mounted on agar blocks containing a 200 µm-deep rectangular well to minimize compression by the coverslip during imaging, supplemented with liquid medium, and covered with a coverslip. DII-Venus was excited at 488 nm and emission collected at 517–544 nm; mDII-tdTomato was excited at 561 nm and emission collected at 575–618 nm. Z-stacks were acquired at 0.54 × 0.54 × 0.54 µm voxel resolution using bidirectional scanning. Laser intensity and master gain were adjusted individually for each seedling to avoid signal saturation.

### DR5 imaging

DR5 imaging was performed on a Zeiss LSM880 equipped with a 20×/0.8 NA water-dipping Plan-Apochromat objective. Seedlings co-expressing DR5-Venus and tdTomato were mounted on agar blocks containing a 200 µm-deep rectangular well, supplemented with liquid medium, and covered with a coverslip. DR5-Venus was excited at 488 nm and emission collected at 517–552 nm; tdTomato was excited at 561 nm and emission collected at 574–618 nm. Identical laser intensity (16% for 488 nm; 7.5% for 561 nm) and master gain (817 for 488 nm; 725 for 561 nm) were used for all replicates. Z-stacks were collected at 0.88 × 0.88 × 1 µm voxel size.

### Macro-confocal time-lapse imaging of DR5::Venus

Imaging was conducted during the hook formation phase (∼16 h after germination). Seedlings were imaged on vertically positioned agar plates using a Nikon AZ-C2 macro-confocal microscope fitted with a 5×/0.5 NA 15 mm macro-objective at 2 h intervals in a dark room. Excitation was performed at 488 nm, with emission collected using a spectral detector. Between laser scans, Petri dishes were wrapped in aluminum foil to limit light exposure.

### FRAP experiment

Fluorescence recovery after photobleaching (FRAP) was performed on a Zeiss LSM800 with a 40×/1.0 NA water-dipping Plan-Apochromat objective. Seedlings at onset of hook maintenance (∼24 h after germination) co-expressing DR5-Venus, H2B-mCherry, and tdTomato were mounted on agar blocks containing a 200 µm-deep rectangular well, supplemented with liquid medium, and covered with a coverslip. DR5-Venus was excited at 488 nm and emission collected at 400–534 nm; H2B-mCherry and tdTomato were excited at 561 nm and emission collected at 587–700 nm. A pre-bleach Z-stack was acquired. Circular regions of interest (ROIs) were then drawn over nuclei and photobleached with 488 nm and 561 nm lasers at 100% power until DR5-Venus and H2B-mCherry nuclear fluorescence was strongly reduced. Immediately afterward, a post-bleach Z-stack was acquired. The coverslip was then removed, and seedlings were incubated vertically on agar blocks inside 3D-printed dark chambers. Recovery was monitored by acquiring Z-stacks of the same region at 1 h intervals over 3 h. Identical acquisition settings were used for pre- and post-bleach imaging to allow direct comparison of fluorescence recovery. Different laser intensities were used between replicates to avoid signal saturation. Voxel size was 0.62 × 0.62 × 0.62 µm.

### Image analysis

#### Time-lapse imaging of cell elongation

Confocal stacks were processed and analyzed using MorphoGraphX software (Barbier de Reuille et al. 2015; Strauss et al. 2022). Stacks were manually stitched into a single file using the “combine stack” tool. Stacks were blurred using one iteration of Gaussian blur with XYZ values between 0.5–2. Surface detection was done with the “edge detect” tool using a threshold from 8000–10000. Meshes were created with an initial cube size of 5□µm subdivided 2 times before projecting membrane signal (5–10 □µm from the mesh). Segmentation was performed by manual seeding. Parent relations between cells of successive time points were attributed manually.

#### Growth analysis

An organ-based coordinate system was created using a custom Bézier line that closely followed the centerline of the organ, starting from the hypocotyl apex. The Bézier line was generated either manually or using an automated MorphoGraphX plugin that fits a curve through a selected longitudinal cell file. Cells were assigned longitudinal (*s*) and radial (*x*) coordinates relative to the Bézier line, such that *x*< 0 corresponded to cells on the inner side, and *x*> 0 the outer. Cell time-lapse tracking (“Parent labels”) were used to compute cell growth. In order to quantify cell longitudinal strain *ε* (*s, x*), principal strains and their associated orientations (PDGs) were extracted and used to evaluate the directional strain along the local longitudinal direction defined by the Bézier curve.

Midline elongation (*ε*_mid_) and differential growth (Δ) were computed along the Bézier curve using a single procedure. For a given longitudinal position (*s*), longitudinal strain *ε*(*s, x*) were extracted for all cells within a window spanning 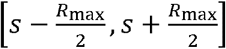, where *R*_max_ is the maximal lateral extent of the sample. Within this window, a linear fit of elongation rate as a function of radial position was performed, *f*(*x*) = *ax* + *b*, where *f* is the elongation rate and *x* the radial coordinate. The midline strain *ε*_mid_ was estimated from the intercept *b*, and differential growth Δ was quantified as the normalized lateral gradient *a*/*b*. (see Eqs.S11,S14 in the **SI**).

#### Curvature analysis

Organ curvature was measured using the Bézier line following the organ midline. Bézier control points were used to reconstruct the Bézier line in Python, using the *Bezier* package, for which the 3D curvature was found using 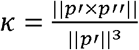, where the curve is parametrised in 3D Cartesian coordinates, *p*(*t*) = (*x*(*t*), *y*(*t*), *z*(*t*)). By approximating the curve as planar, we treated the curvature as the signed curvature, set to be positive along the WT hook. That is, while traveling from the hook apex to the base (increasing arc length *s*), *κ* > 0 corresponds to a curve turning towards the right.

#### Auxin related treatment analysis

Samples treated with IAA, auxinole, fusicoccin, and their respective mocks were analysed for their change in elongation rate over time. Per time point average longitudinal elongation rate and curvature were found as described above. Regions of interest on the organ were then defined using the custom Bézier coordinates. First, cells were divided into “inner” and “outer” as described above. Based on the point of maximum curvature (*s*= 0), cells with - 200 μm < *s* < 0 μm were classed as “apical”, and cells with 0 μm < *s* < 200 μm were classed as “basal”. The difference in average longitudinal elongation rate between two successive time points (mock and treated) was calculated for each of the four regions: inner apical (ia), outer apical (oa), inner basal (ib), outer basal (ob). Each treatment at each concentration was tested against their mock using a two-sided Mann-Whitney U rank test.

#### R2D2 analysis

Confocal stacks were processed and analyzed using MorphoGraphX software (Barbier de Reuille et al. 2015). mD2-mCherry signal was used for nuclear volume extraction. The stacks were filtered with Gaussian blur (sigma = 0.3), followed by Median filter (size 1) and a binarization (threshold 5000). Meshes were created using Marching Cubes Surface (cube size 1□µm), and subsequently smoothed 2 passes. Nuclei mesh were used to compute the total fluorescent signal of DII-Venus and mDII-tdTomato contained in each nucleus.

The organ surface was extracted from the DII-Venus signal. The stack was blurred using Gaussian filter (sigma = 0.5), followed by Brighten Darken function (value 16), Edge Detect (threshold = 10000) and Edge Detect Angle. The surface mesh was created with the Marching Cubes Surface (cube size = 1 µm), followed by 5 passes of smoothing. To exclusively quantify nuclei from epidermal cells, nuclei were selected based on their perpendicular distance to the outer epidermal surface, using an approximate 20 µm cutoff followed by manual verification. The surface was auto-segmented (auto seed radius 3 µm) to delimit small cell-like regions. The vertical distance from each nucleus to the organ surface was computed by matching the (x,y) position between a nucleus and a cell-like region on the surface.

Based on nuclei DII/mDII signal ratio, we computed the profiles of midline R2D2 ratio and the outer/inner R2D2 ratio along the hook, using a similar method as for computing 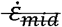 and *Δ* (see Growth Analysis section). The regression of DII/mDII signal ratio along the *x* direction was obtained with weighted least squares. Regression weights were computed based on the vertical distance between each nucleus and the organ surface, according to 1- (depth/30 μm), such that nuclei at the epidermis surface have a value of 1 and those at 30 μm or deeper have a value of 0.

#### FRAP analysis

Confocal stacks of the H2B-mCherry signal were used for surface cell segmentation, parent lineage tracking, and curvature measurement as described above.

DR5::Venus signal was used for nuclear volume extraction. The stack was processed with CNN using the NucleiUNet prediction (Vijayan et al. 2024) with a resample factor of 2, followed by Trimming (threshold = 28,000) and Morphological Closing (radius = 2) with a rounded neighborhood. Processed stacks were binarized with a threshold of 28,000 and manually cleaned. Next, nuclei surface were extracted with Marching Cubes 3D (size = 1), and smoothed (1 pass). Nuclei were assigned matching labels with cells segmented on the surface mesh, allowing to track their movements.

The signal intensity per unit volume was measured per nucleus for both stacks (DR5::Venus and H2B-mCherry). For unbleached nuclei, volumetric intensity H2B-mCherry served as a control to correct for variations in intensity of DR5::Venus which would be independent from auxin (e.g. caused by different light absorption by surrounding tissue). At each timepoint DR5::Venus signal was normalized by the H2B-mCherry from the same nucleus.

Both DR5::Venus and H2B-mCherry signals strongly decreased after bleaching, which prevented applying the same procedure. Therefore, for every bleached nucleus, the signal used for DR5::Venus normalization was obtained by computing the average H2B-mCherry volumetric intensity of its closest neighbours (within 50 μm in longitudinal position). The corrected DR5::Venus signal at Prebleach was taken as the reference to measure relative changes in intensity.

All nuclei were assigned a region based on their position at the Prebleach time point relative to the position of maximum curvature (*s*= 0). Those with an arc length position of less than -100 μm were classed as apical, those with an arc length position of more than 100 μm were classed as basal, and those in between were classed as central. Plots were generated in Python using *lineplot* from the *seaborn* package.

### Numerical integration, plotting and statistical analysis

The dynamics of the rod model were numerically integrated using the method described in (Porat et al. 2020). Plots were created in Python using the *matplotlib* and *seaborn* packages. Statistical tests were performed in Python using the *scipy* package.

### Visualization

Image processing for visualization was performed using Fiji. Interpolated morphing movies were generated using Abrosoft Fantamorph (Abrosoft Co.) to bridge consecutive imaging time points; these visualizations were used for illustrative purposes only and were not included in quantitative analyses. Figures were assembled using Adobe Illustrator, and image cropping and layout adjustments were performed in Adobe Photoshop.

## Acknowledgments

We thank Rishikesh Bhalerao for helpful discussions and comments. This work was supported by the Åforsk Repatriation Grant (24-749) and the Vetenskapsrådet International Postdoctoral Grant (2020-06442) to K.J., the Gatsby Charitable Foundation (GAT3395/PR4B) to A.P and H.J, The Herchel Smith Postdoctoral Fellowship to A.P., the Israel Science Foundation Research Grant (ISF) no. 2307/22, and ERC grant GROWsmart 101165101 to Y.M and CRC, Discovery, and HFSP to A.-L.R.-K.

## Resource availability

### Lead contact

Further information and requests for resources and reagents should be directed to and will be fulfilled by the lead contact, Anne-Lise Routier-Kierzkowska

### Materials availability

Plant lines used in this study are available from the lead contact upon request, subject to standard material transfer agreements where applicable.

### Data and code availability

Microscopy datasets generated in this study are available from the lead contact upon reasonable request.

Custom analysis scripts used for curvature measurements, growth analysis, and data processing are available from the lead contact upon request.

The code implementing the numerical integration of the shape dynamics is available at https://github.com/poratamir/Hook_animations.

This study did not generate large-scale sequencing or proteomic datasets.

## Author Contributions

Conceptualization, K.J., A.P., and A.-L.R.-K.; methodology, K.J., A.P., V.A., S.H., and A.- L.R.-K.; software, A.P., V.A., and S.H.; investigation, K.J., A.P., and V.A.; formal analysis, K.J., A.P., V.A. and A.-L.R.-K.; data curation, V.A.; visualization, K.J., A.P., V.A., and A.- L.R.-K.; supervision, H.J., Y.M. and A.-L.R.-K.; writing – original draft, K.J., A.P., and A.- L.R.-K.; writing – review and editing, K.J., A.P., V.A., H.J. and Y.M.; and funding acquisition, K.J., A.P., H.J, Y.M., and A.-L.R.-K.

## Declaration of generative AI and AI-assisted technologies in the manuscript preparation process

During the preparation of this work, the authors used ChatGPT, Grok, and Claude to assist with language editing and clarity. After using these tools, the authors have reviewed and edited the content as needed and take full responsibility for the content of the published article.

## Supplementary figure legends

**Figure S1.**
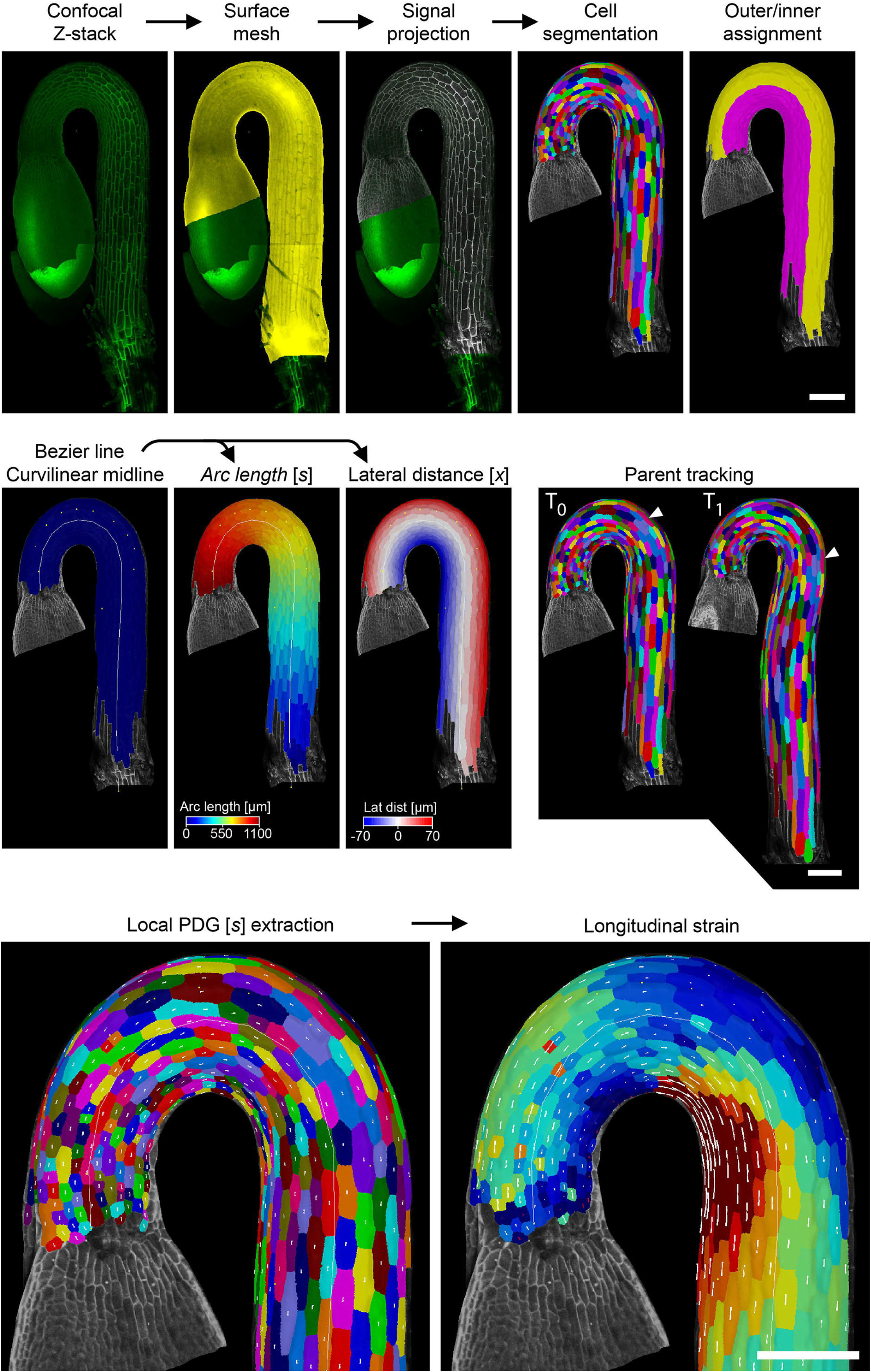
MorphographX growth analysis pipeline. Confocal Z-stacks served as templates for generating surface meshes. Fluorescence signal was projected onto the mesh surfaces to obtain cell outlines, followed by manual seeding for cell segmentation. Inner and outer lateral sides of the hook were assigned. A Bezier curve was fitted along the organ midline to establish a curvilinear coordinate system, from which we obtained arc length position (*s*) and lateral position (*x*). Cells were assigned persistent identities (parented) to enable tracking of individual cells across time points. Local principal directions of growth (PDGs) were computed relative to the Bezier curve, from which longitudinal strain rates were extracted. For detailed information, see methods section

**Figure S2.**
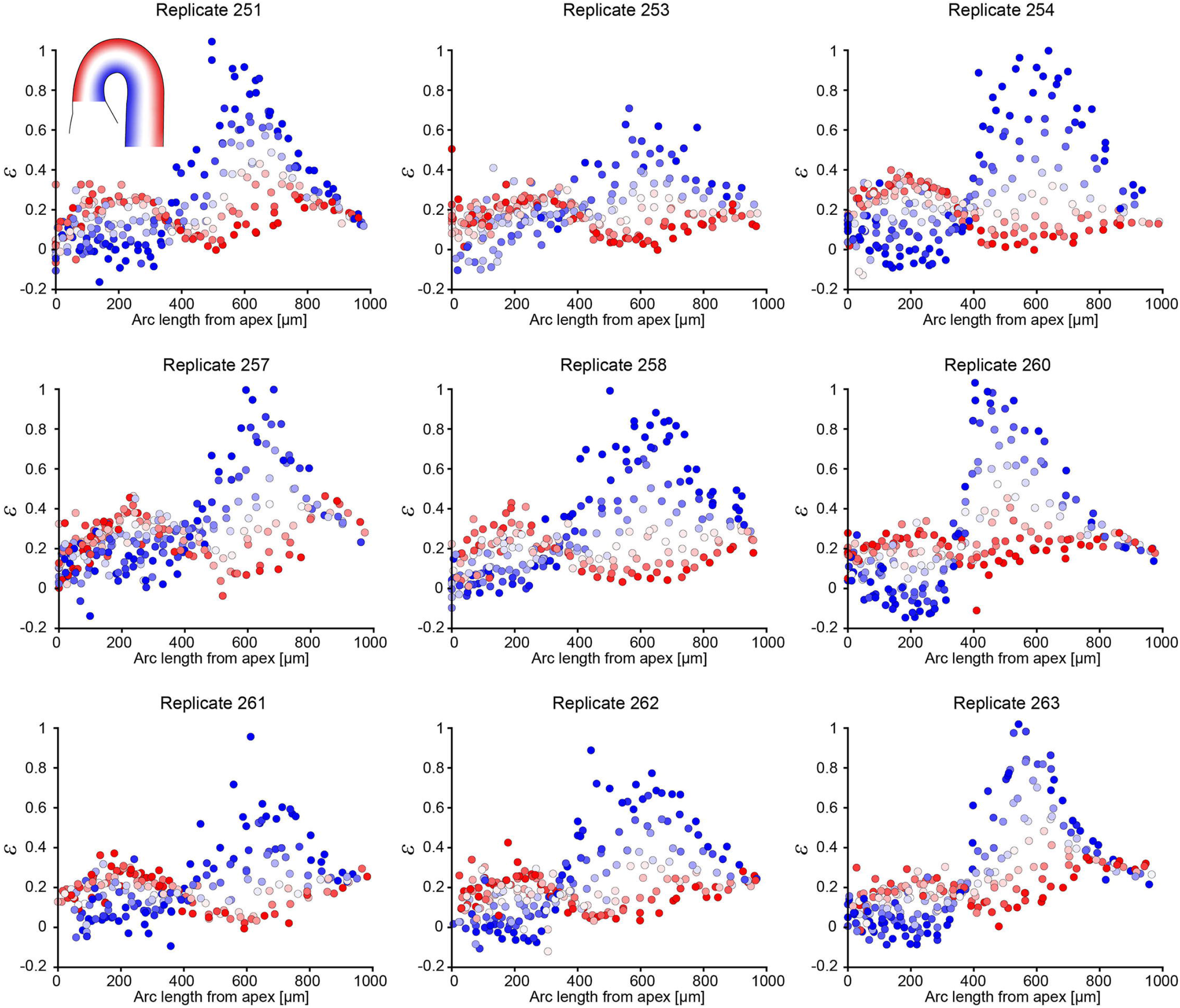
Single-cell strain patterns at onset of hook maintenance in WT (replicate datasets underlying Fig. 1F). Single-cell strain (*ε*) patterns from onset of hook maintenance (∼24 h after germination) in WT. Each point represents a single cell, color-coded by radial position within the hypocotyl tissue (Indicated by inset; blue: inner; red: outer).

**Figure S3.**
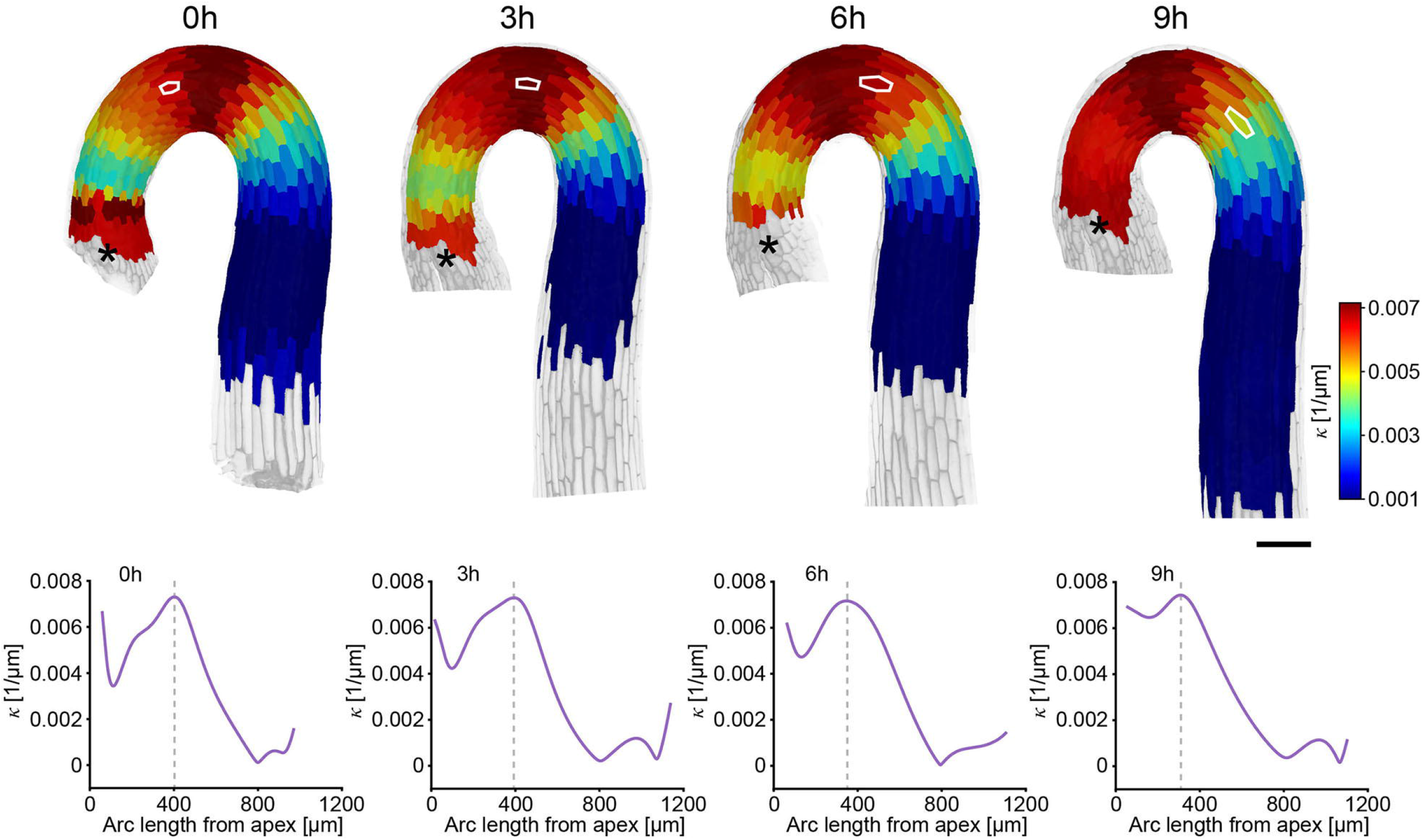
Temporal stability of the curvature profile in WT during early maintenance. Top: Representative time-lapse series showing heatmaps of midline curvature at 3 h intervals of a single WT seedling starting from onset of the hook maintenance phase (denoted *t* = 0 h). The spatial distribution of curvature remains largely unchanged over time, while tissues elongate and are displaced along the organ. White contours indicate the same cell over time. Asterisks indicate hypocotyl apex. Scale bar, 100 µm Bottom: Curvature plots corresponding to heatmaps. Vertical dotted lines indicate point of maximal curvature

**Figure S4.**
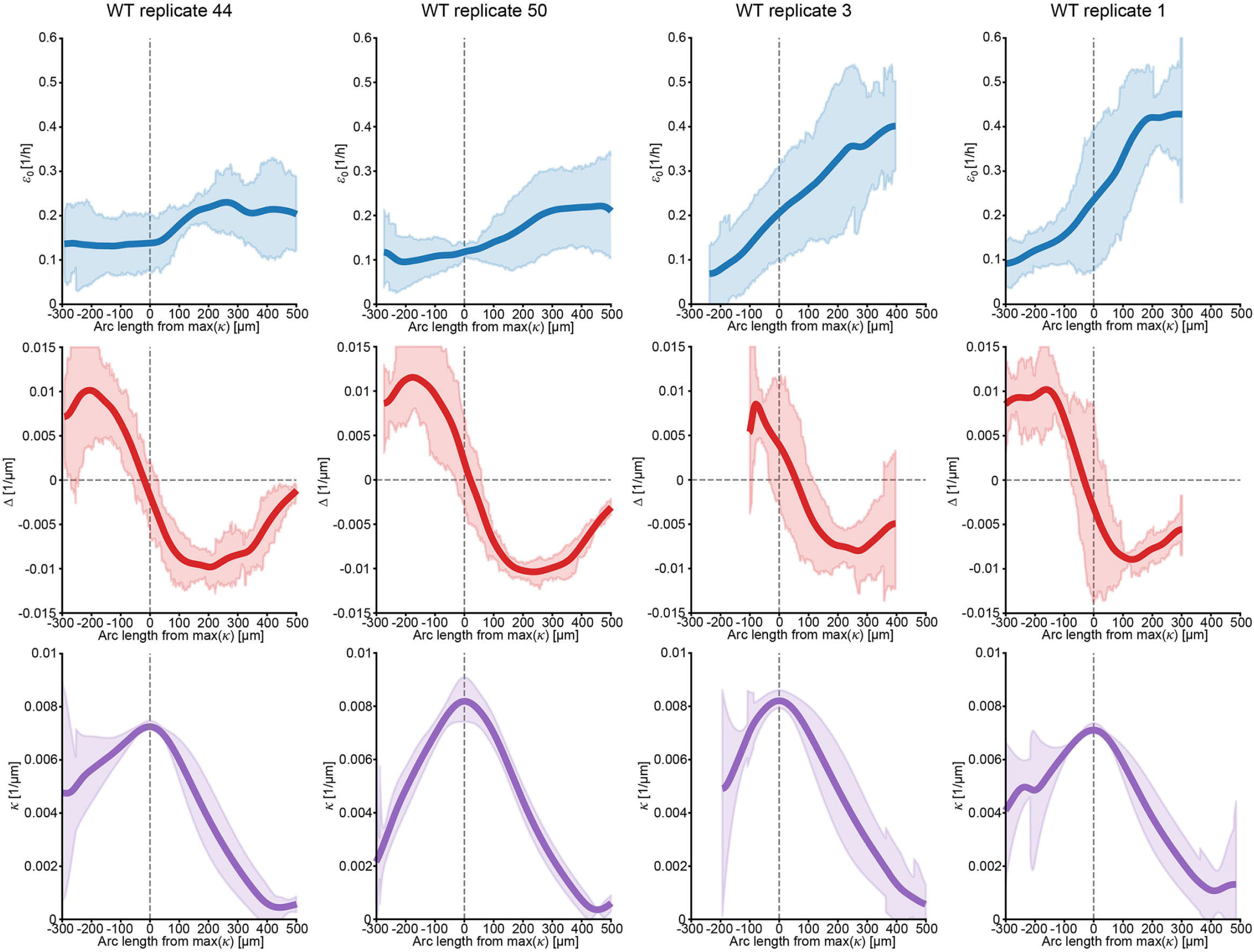
Patterns of *ε*_mid_, Δ and *κ* during hook maintenance in WT. *ε*_mid_, Δ and *κ* patterns during the maintenance phase of apical hook development. Data for each individual seedling are averaged over four sequential 3 h time steps, with x-axis positions centered on the position of maximal curvature. Vertical dotted lines indicate position of maximal curvature for each replicate. Horizontal dotted lines indicate Δ= 0. *n* = 4.

**Figure S5.**
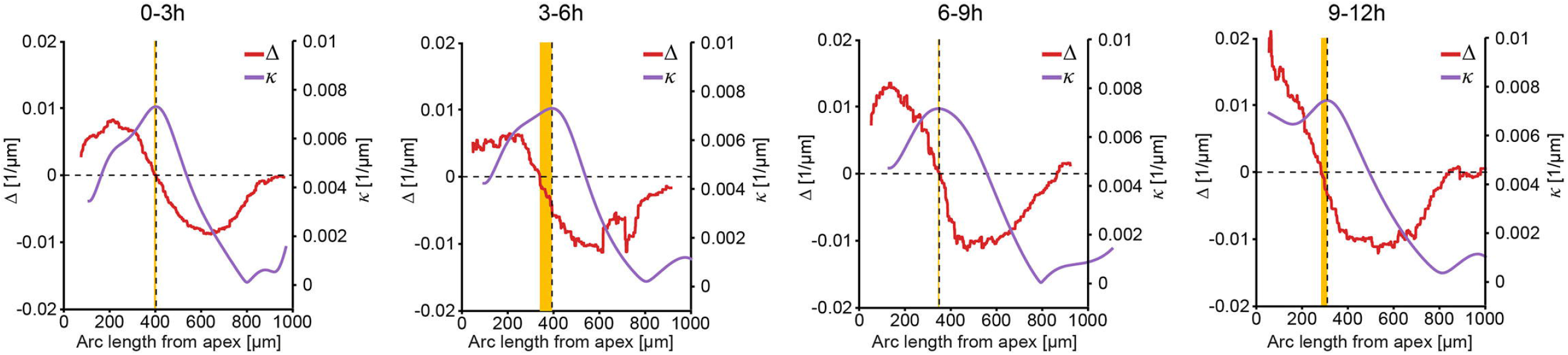
Temporal alignment between maximal curvature and Δ sign change from onset of hook maintenance in WT Curvature. *κ* and lateral growth gradient Δ profiles extracted from four consecutive 3h intervals shown in Fig. 2A. The vertical dotted line indicates the position of maximal curvature; the horizontal dotted line indicates Δ= 0. The orange shaded area represents the midline axial offset between these positions. Under self-similar stationary growth (Eq. 6), the position of maximal curvature and Δ=0 are predicted to coincide. The consistently narrow shaded areas across intervals indicate stable alignment, supporting the predicted growth–curvature coupling.

**Figure S6.**
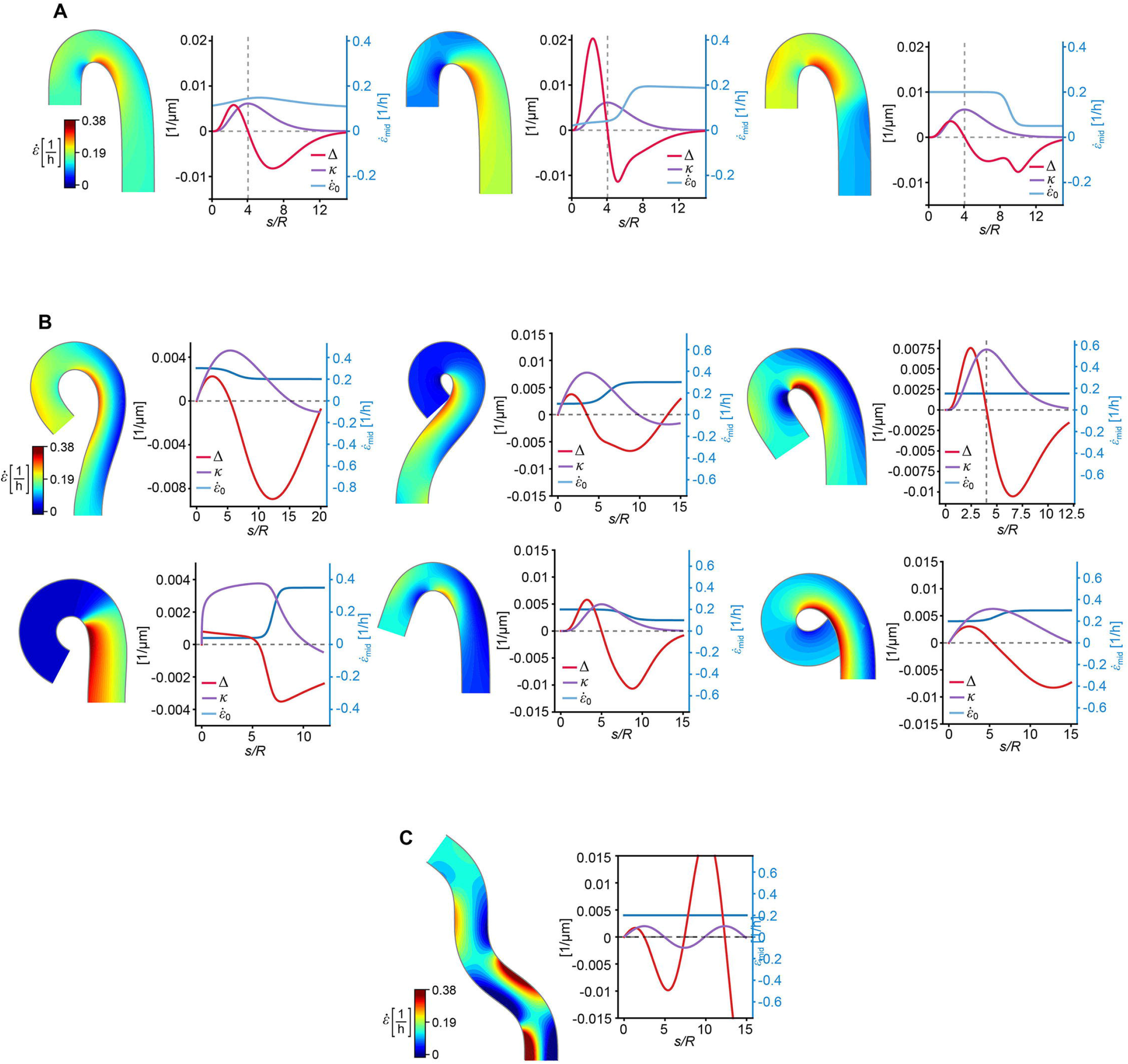
Constraints and degeneracy of growth patterns compatible with self-similar hook shapes. Using our modelling framework (Eq. 2), we fix the WT curvature profile *κ* and ask which growth fields 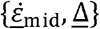 are compatible with maintaining this shape under self-similar growth. We found two main constraints on the these fields: First, as for strictly growing organs 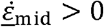 and |*R*Δ|≤ 2 (see SI, Eq. S19), Eq.2 shows that self-similar maintenance imposes a lower bound on the axial growth rate via 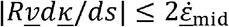. Second, the maximum of the curvature profile (*dκ*/*ds* = 0) necessarily corresponds to Δ crossing the *s* axis, changing sign from positive to negative. Despite these constraints, the space of admissible growth patterns remains infinite. **(A)** To illustrate that the WT curvature profile can arise from multiple coordinated 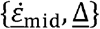 combinations, we sample this space of growth patterns. This demonstrates that hook shape—even the full *κ* profile—does not uniquely determine the underlying growth dynamics. Instead, the system admits multiple self-similar solutions, with stable shapes extending well beyond the WT parameter set. **(B)** More generally, any self-similar hook shape is admissible provided |*Rκ*|< 1 and self-intersections are avoided. Here, examples of self-similar growth patterns generating non-WT shapes with a range of hook angles are shown. **(C)** Our model predicts that a self-similar shape with constant 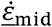 and oscillating curvature produces an oscillatory Δ with linearly increasing amplitude. Following Eq.2 in the main text, this amplification arises from the linear increase in the velocity of material segments as they advect through the shape. This edge case both resembles and explains the asymmetric Δ profile observed during WT maintenance, in which the basal lateral gradient is higher than the apical lateral gradient.

**Figure S7.**
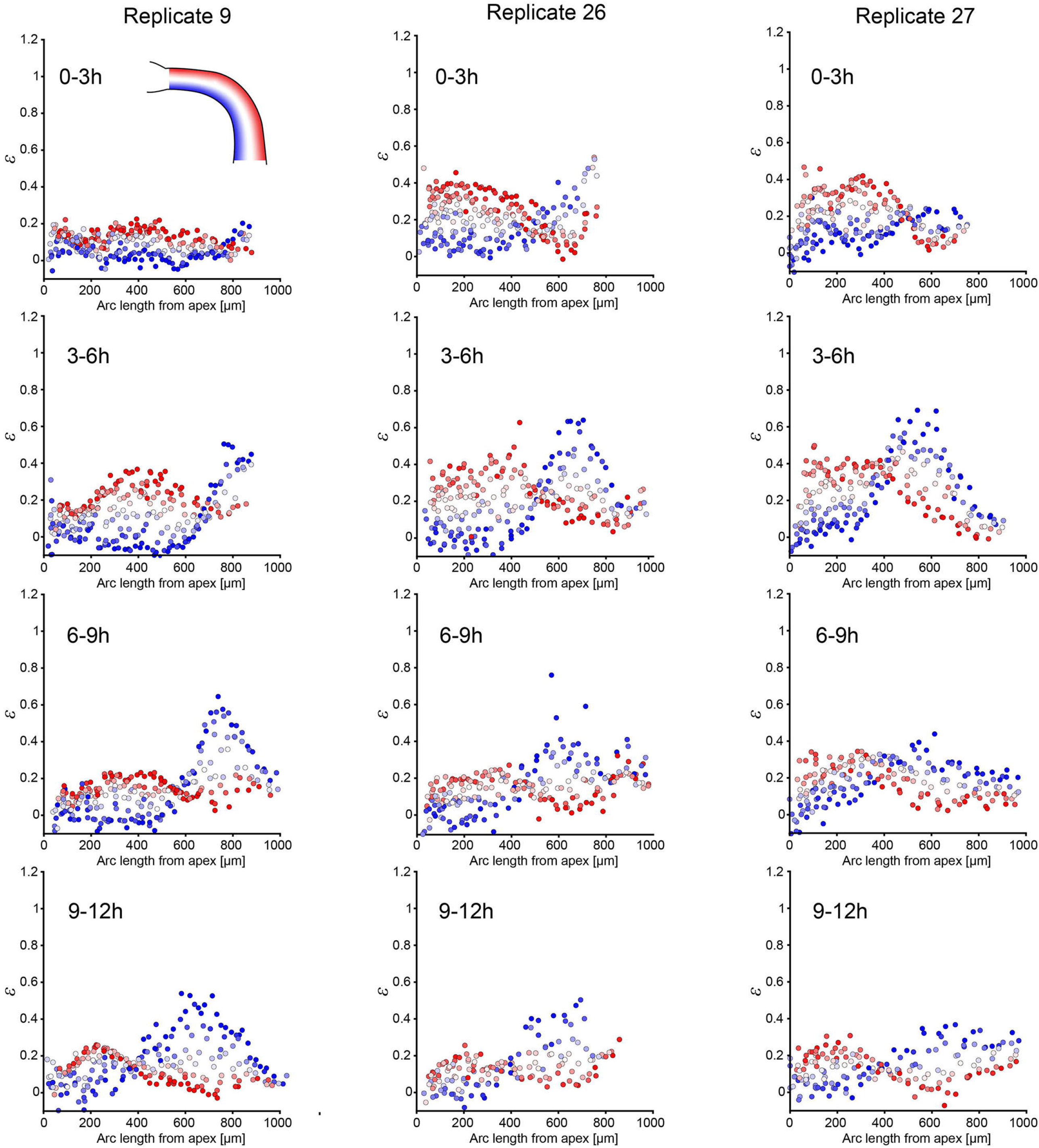
Single-cell strain patterns during WT hook formation (replicate datasets underlying Fig. 4B) Single-cell strain (*ε*) patterns over four successive 3 h intervals during the formation phase of hook development in WT under mock conditions. Each point represents a single cell, color-coded by radial position within the hypocotyl tissue (Indicated by inset; blue: inner; red: outer).

**Figure S8.**
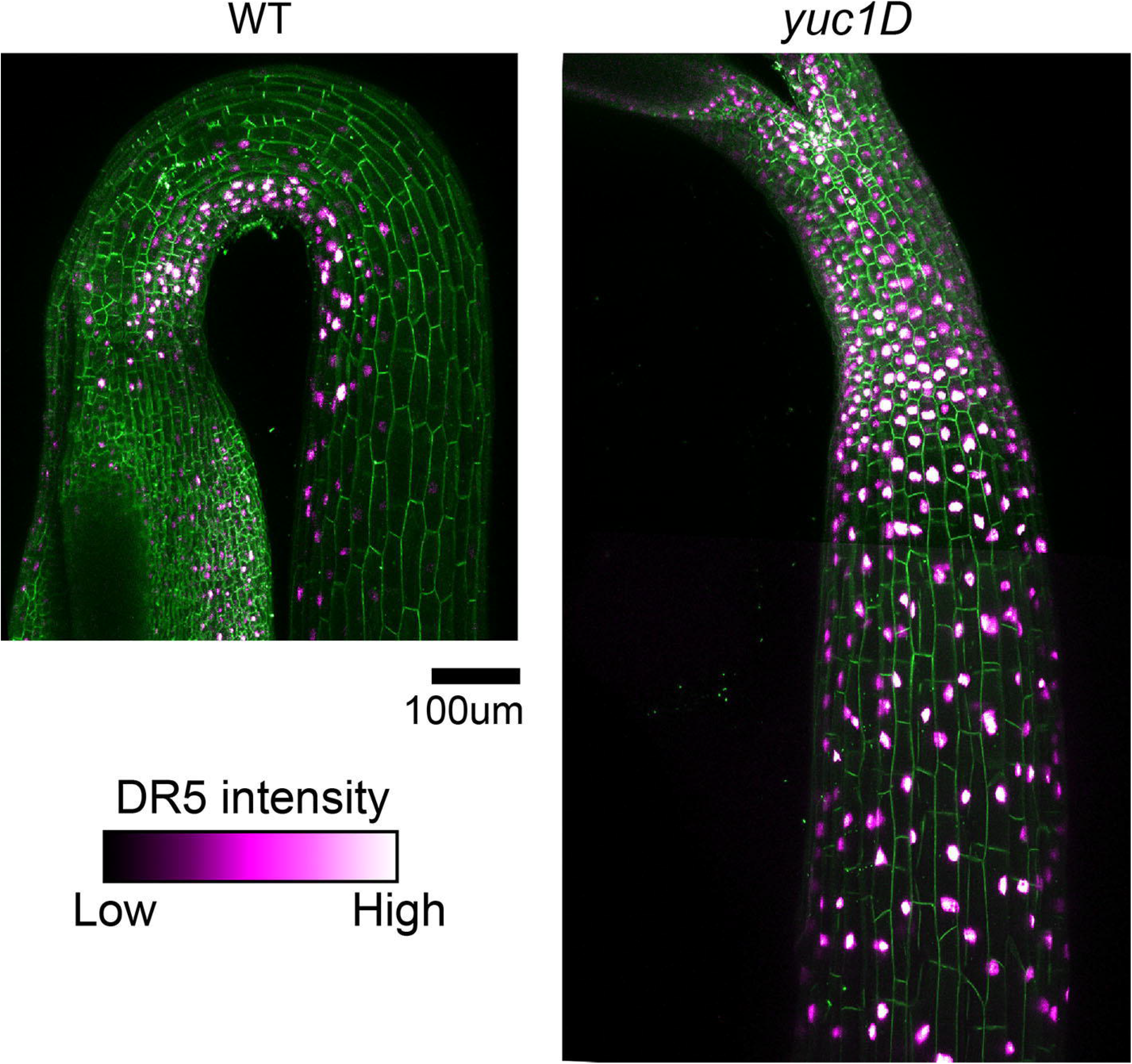
DR5 is expressed ubiquitously in *yuc1D*. Representative confocal images of DR5 fluorescence in WT and *yuc1D* at onset of hook maintenance phase (∼30 h after germination). While signal is confined to the inner side in WT, DR5::Venus is expressed ubiquitously throughout the hook region in *yuc1D*. (*n* = WT 16; *yuc1D* 10). Magenta: DR5::Venus; Green: PM-tdTomato. Scale bar, 100 μM

**Figure S9.**
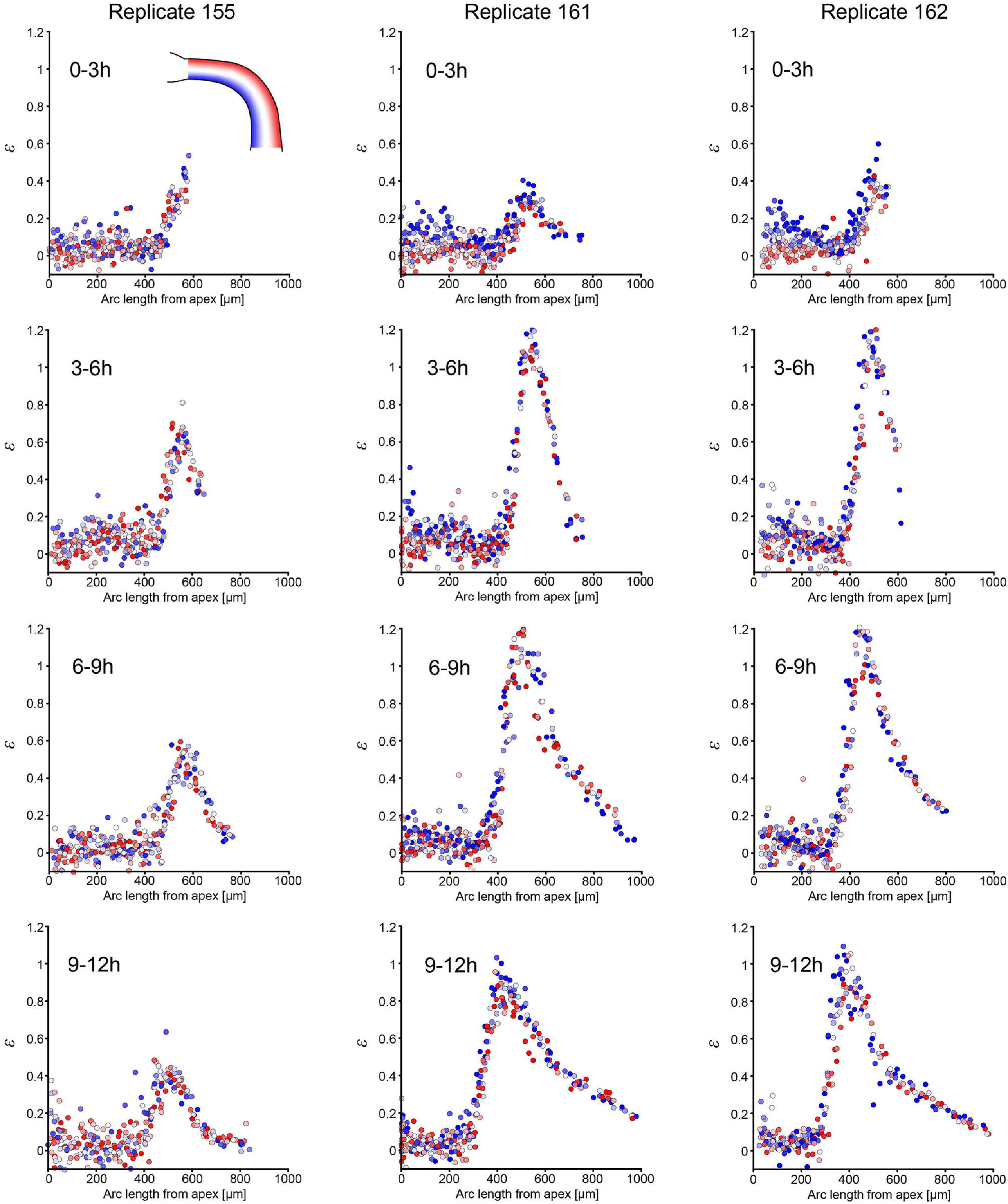
Single-cell strain patterns during hook formation in WT treated with NPA (replicate datasets underlying Fig. 4E) Single-cell strain (*ε*) patterns over four successive 3 h intervals during the formation phase of hook development in WT germinated on NPA 5μM. Each point represents a single cell, color-coded by radial position within the hypocotyl tissue (Indicated by inset; blue: inner; red: outer).

**Figure S10.**
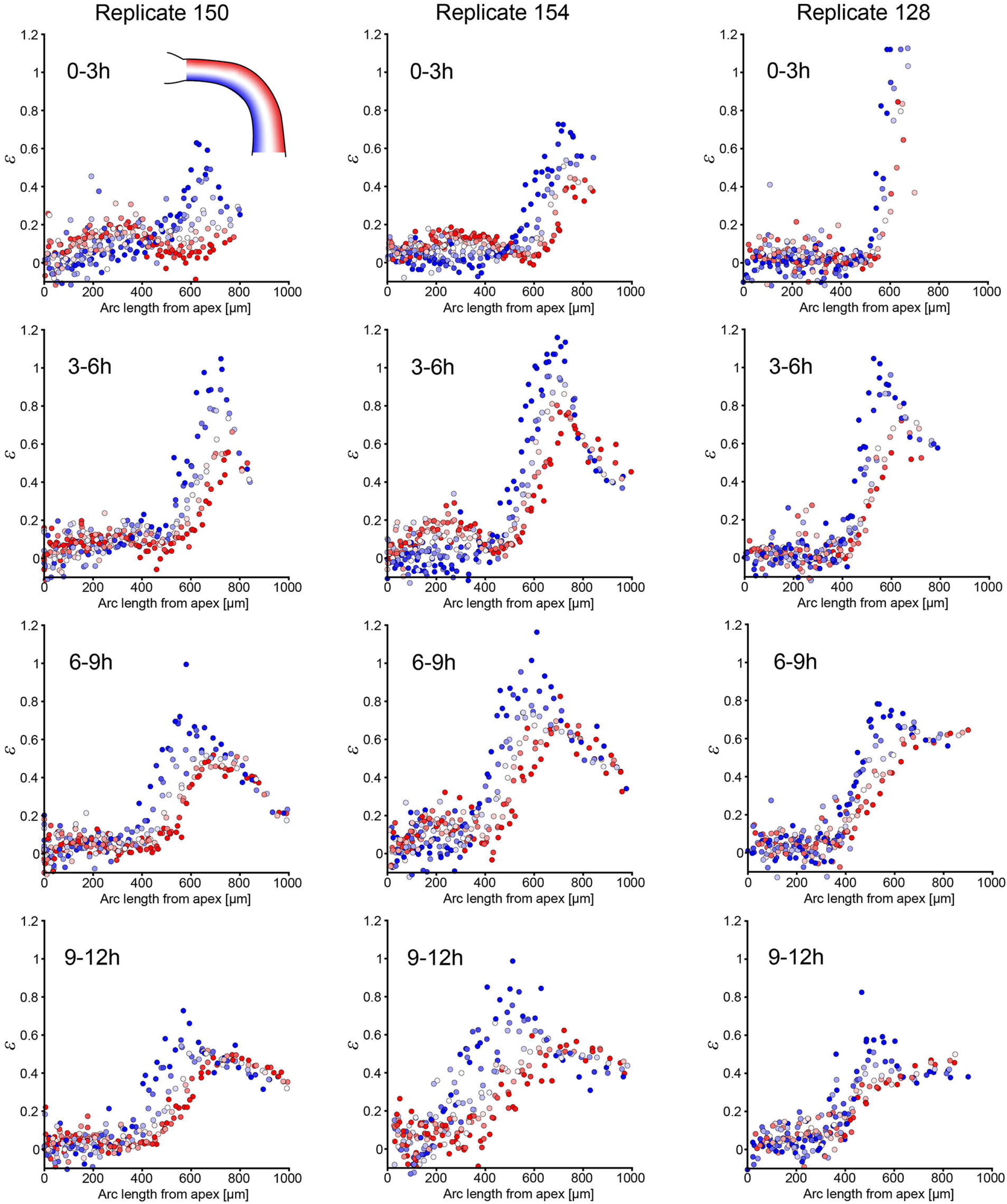
Single-cell strain patterns during hook formation in *yuc1D* (replicate datasets underlying Fig. 4G) Single-cell strain (*ε*) patterns over four successive 3 h intervals during the formation phase of hook development in *yuc1D*. Each point represents a single cell, color-coded by radial position within the hypocotyl tissue (Indicated by inset; blue: inner; red: outer).

**Figure S11.**
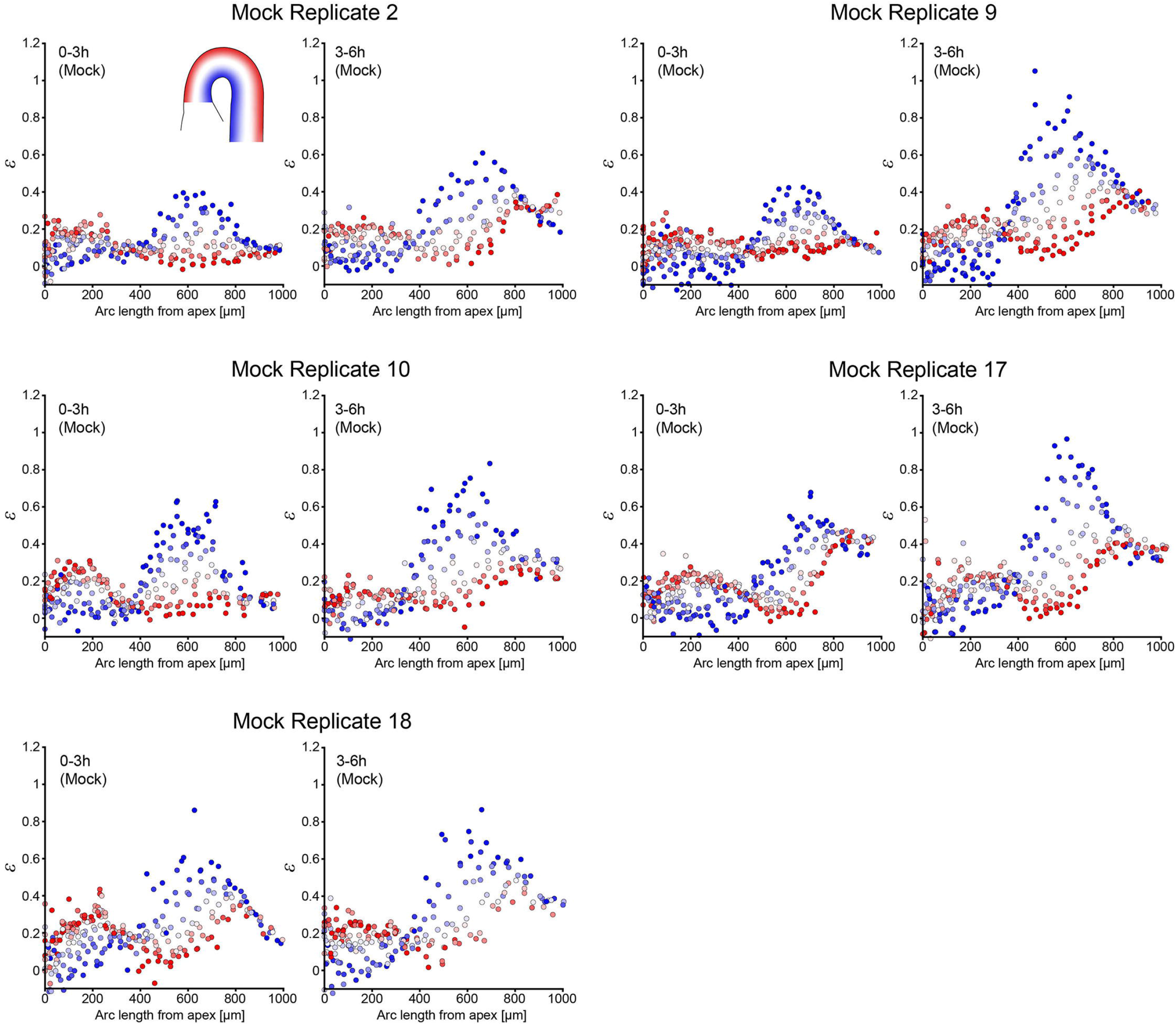
Single-cell strain patterns during WT hook maintenance under mock conditions (replicate datasets underlying Fig. 4H and 4L) Single-cell strain (*ε*) patterns from onset of hook maintenance (designated *t* = 0 h) in WT under mock conditions. Two successive 3 h intervals are shown (0–3 h and 3–6 h), serving as a control for the IAA treatment time course (see **Fig. S12-14**). Each point represents a single cell, color-coded by radial position within the hypocotyl tissue (Indicated by inset; blue: inner; red: outer).

**Figure S12.**
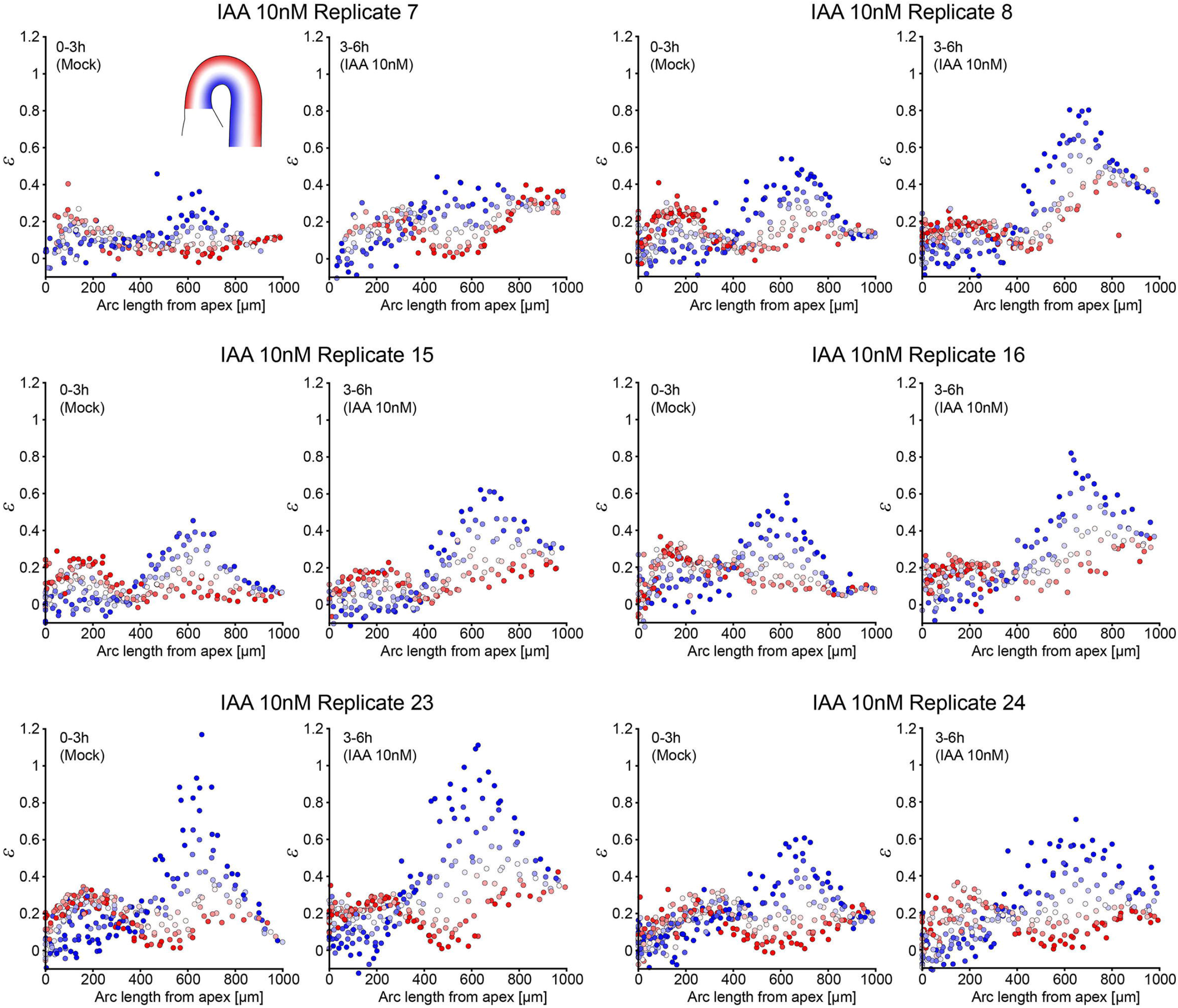
Single-cell strain response to 10 nM IAA during hook maintenance (replicate datasets underlying Fig. 4H and 4L) Single-cell strain (*ε*) patterns from the onset of hook maintenance (designated *t* = 0 h) in WT. Successive 3 h intervals of mock (0–3 h) and IAA (10 nM; 3–6 h) are shown. Each point represents a single cell, color-coded by radial position within the hypocotyl (Indicated by inset; blue: inner; red: outer).

**Figure S13.**
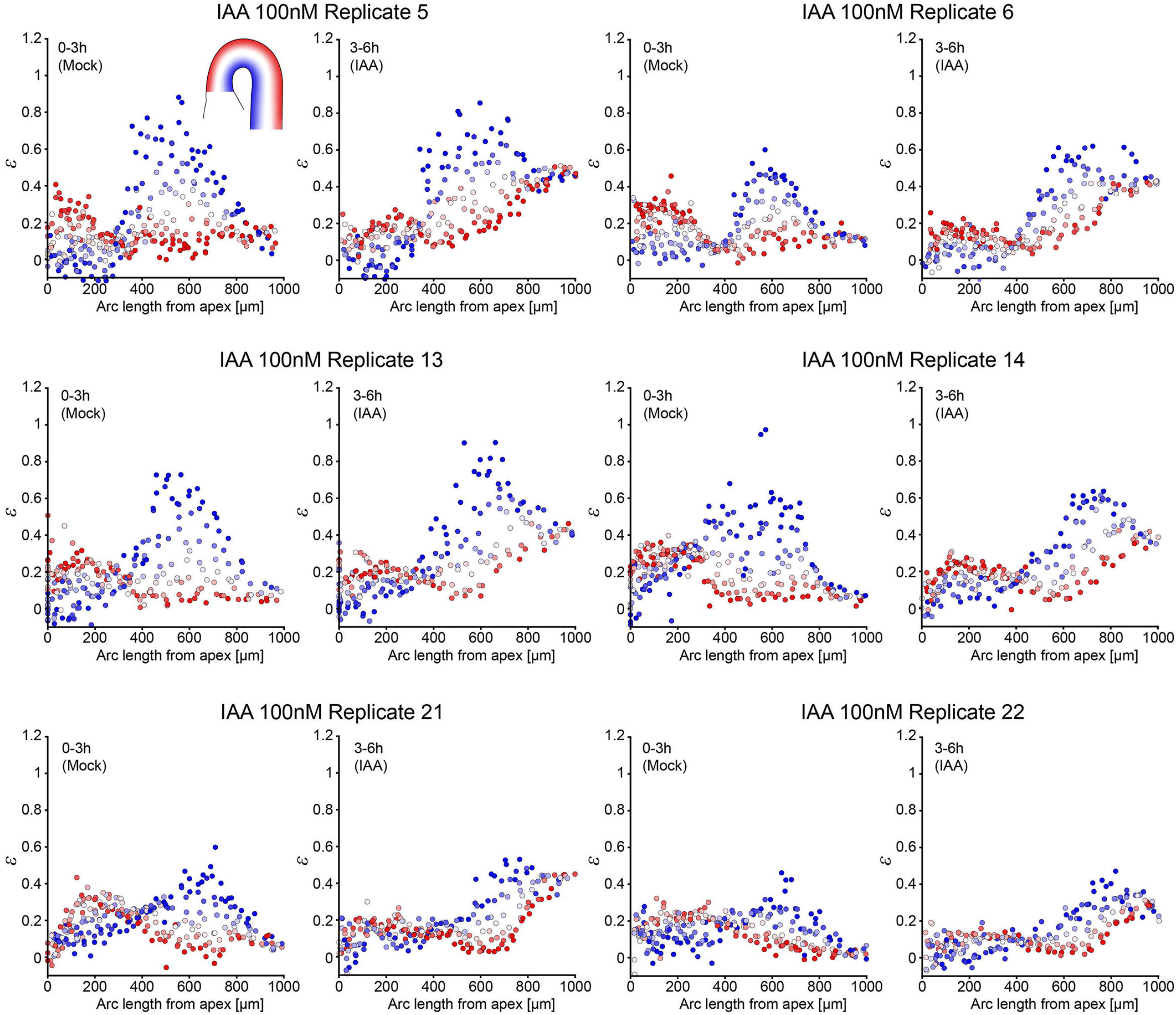
Single-cell strain response to 100 nM IAA during hook maintenance (replicate datasets underlying Fig. 4H and 4L) Single-cell strain (*ε*) patterns from the onset of hook maintenance (designated *t* = 0 h) in WT. Successive 3 h intervals of mock (0–3 h) and IAA (100 nM; 3–6 h) are shown. Each point represents a single cell, color-coded by radial position within the hypocotyl (Indicated by inset; blue: inner; red: outer).

**Figure S14.**
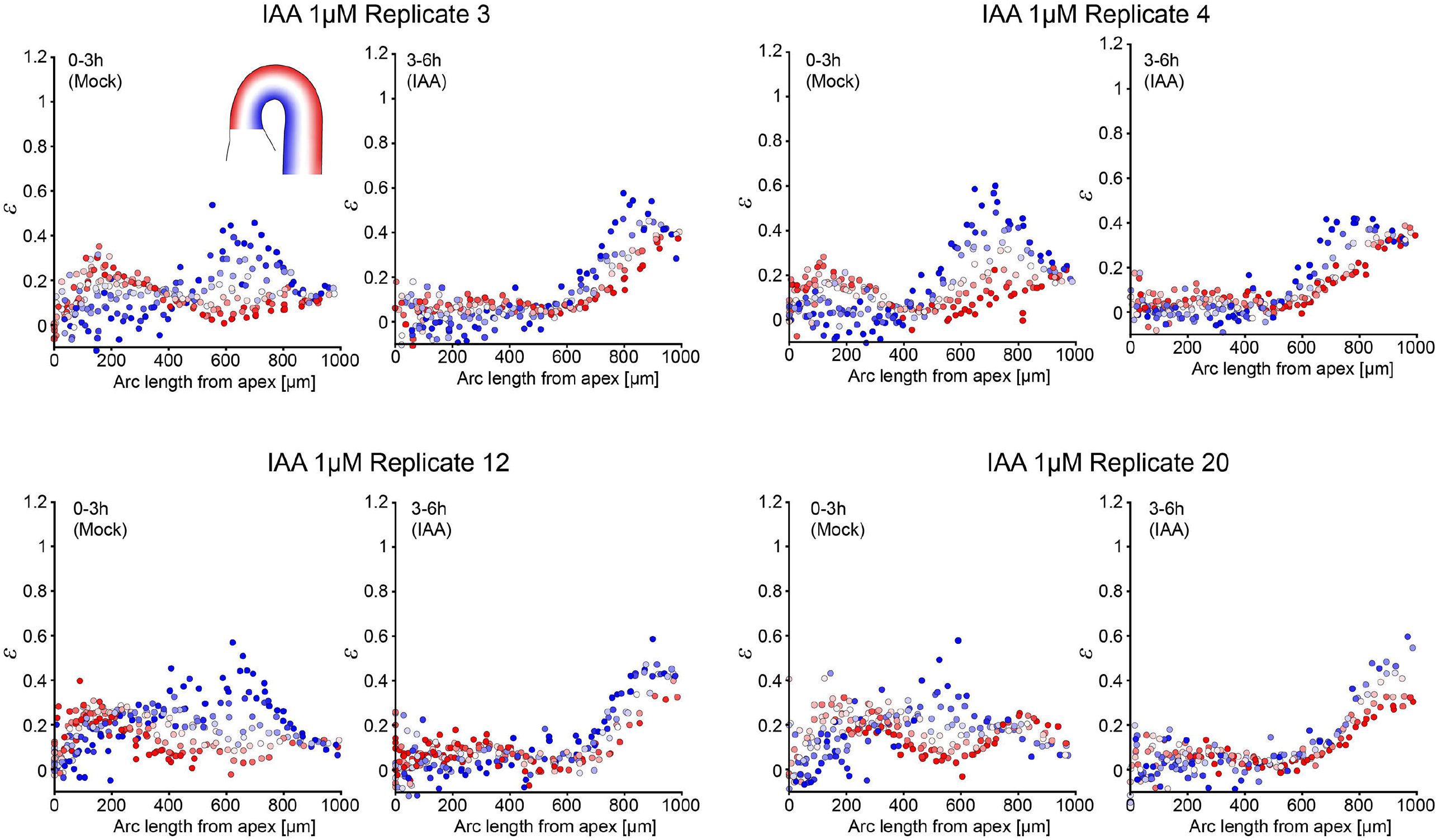
Single-cell strain response to 1 μM IAA during hook maintenance (replicate datasets underlying Fig. 4H and 4L) Single-cell strain (*ε*) patterns from the onset of hook maintenance (designated *t* = 0 h) in WT. Successive 3 h intervals of mock (0–3 h) and IAA (1 μM; 3–6 h) are shown. Each point represents a single cell, color-coded by radial position within the hypocotyl (Indicated by inset; blue: inner; red: outer).

**Figure S15.**
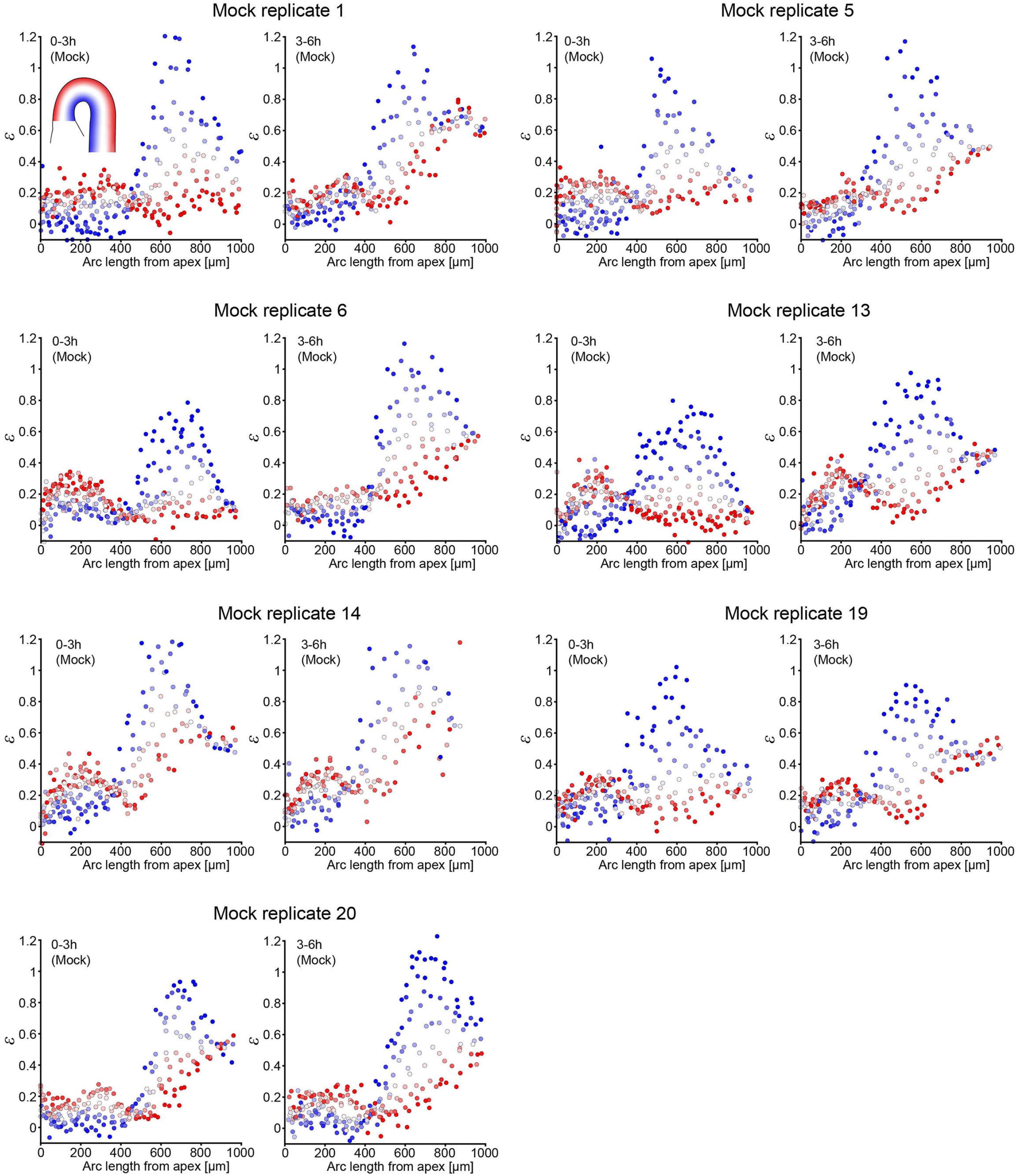
Single-cell strain patterns during WT hook maintenance under mock conditions (replicate datasets underlying Fig. 4H and 4L) Single-cell strain (*ε*) patterns from onset of hook maintenance (designated *t* = 0 h) in WT under mock conditions. Two successive 3 h intervals are shown (0–3 h and 3–6 h), serving as a control for the Auxinole treatment time course (see **Fig. S16**). Each point represents a single cell, color-coded by radial position within the hypocotyl tissue (Indicated by inset; blue: inner; red: outer).

**Figure S16.**
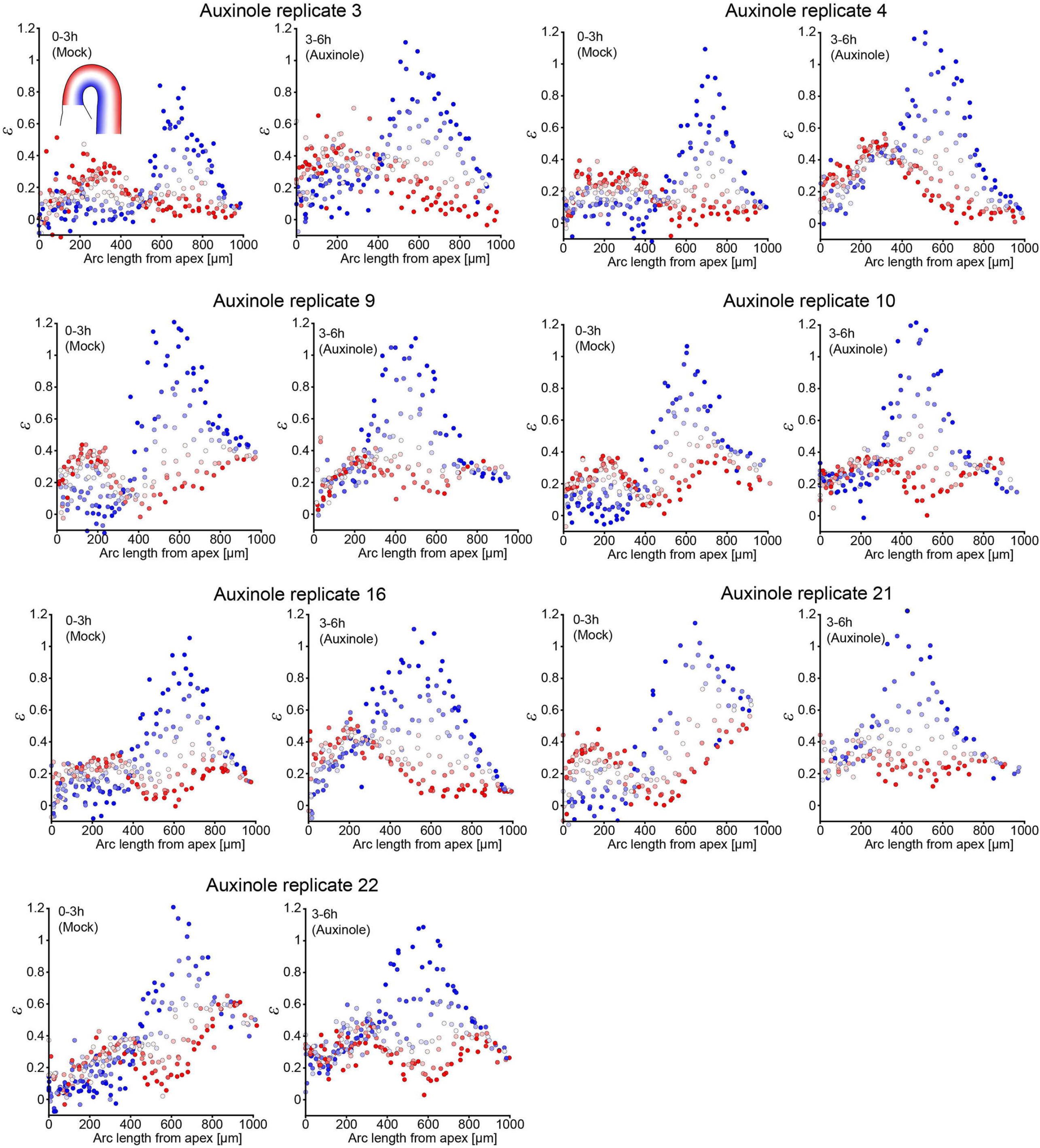
Single-cell strain response to 50 μM Auxinole during hook maintenance (replicate datasets underlying Fig. 4I and 4M) Single-cell strain (*ε*) patterns from the onset of hook maintenance (designated *t* = 0 h) in WT. Successive 3 h intervals of mock (0–3 h) and Auxinole (50 μM; 3–6 h) are shown. Each point represents a single cell, color-coded by radial position within the hypocotyl (Indicated by inset; blue: inner; red: outer).

**Figure S17.**
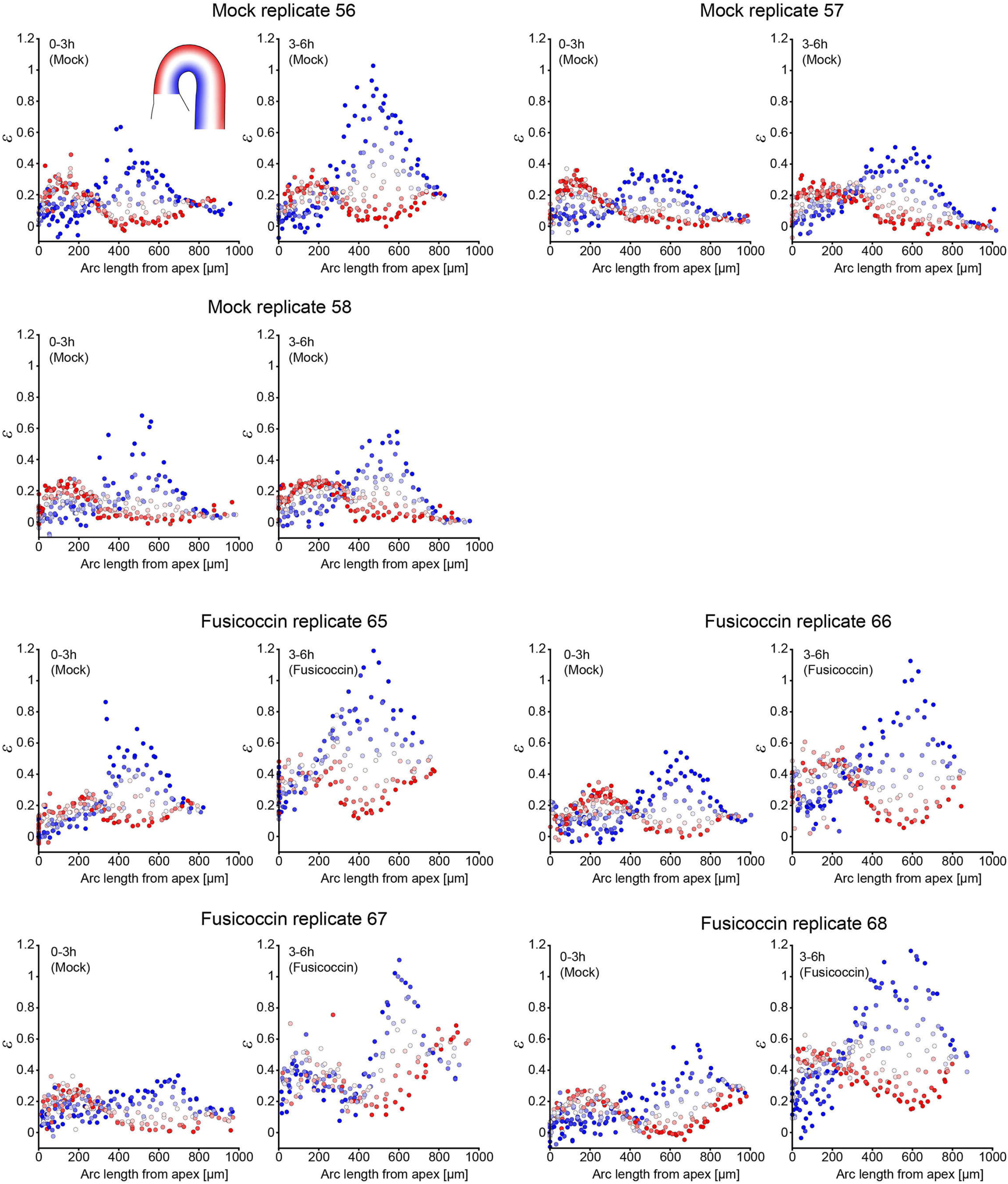
Single-cell strain response to 1 μM Fusicoccin during hook maintenance (replicate datasets underlying Fig. 4J and 4N) Single-cell strain (*ε*) patterns from the onset of hook maintenance (designated *t* = 0 h) in WT upon Fusicoccin (1 μM) treatment. Successive 3 h intervals are shown for both mock-only replicates (0–3 h and 3–6 h) and treatment replicates consisting of a mock interval (0–3 h) followed by Fusicoccin (1 μM; 3–6 h). Each point represents a single cell, color-coded by radial position within the hypocotyl (Indicated by inset; blue: inner; red: outer).

**Figure S18.**
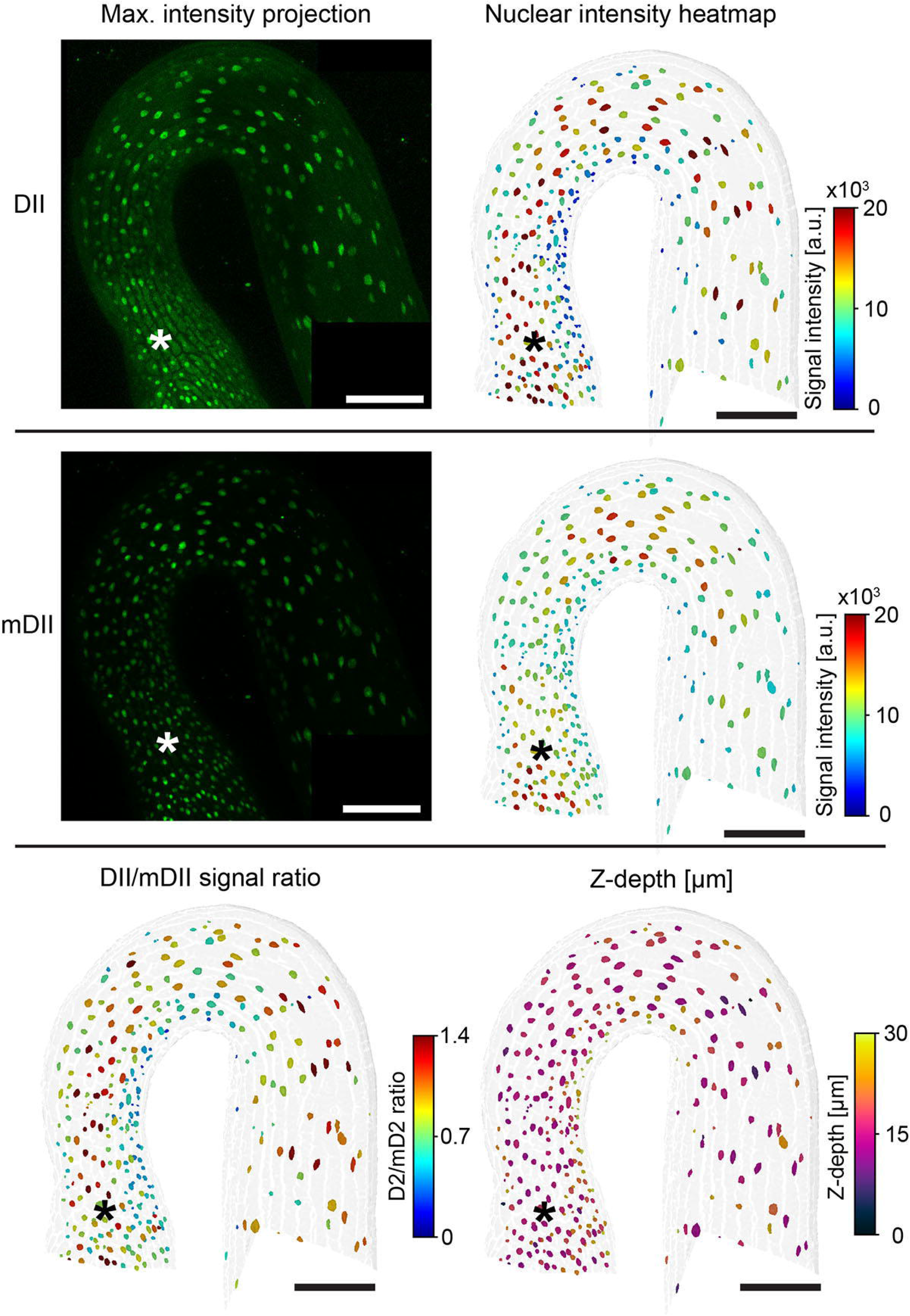
R2D2 signal and nuclear segmentation components used for auxin ratio analysis Maximum-projection views and nuclear segmentation underlying the R2D2 analysis. **Top row:** maximum projection of the DII–Venus channel and the corresponding segmented nuclei colored according to nuclear DII fluorescence intensity. **Middle row:** maximum projection of the mDII–tdTomato channel and segmented nuclei colored according to nuclear mDII fluorescence intensity. **Bottom row:** heatmap of the nuclear DII/mDII signal intensity ratio prior to Z-depth correction (raw ratio), alongside a heatmap indicating nuclear Z-depth within the organ used for subsequent optical path–length adjustment. Asterisks indicate the hypocotyl apex. All scale bars, 100 µm.

**Figure S19.**
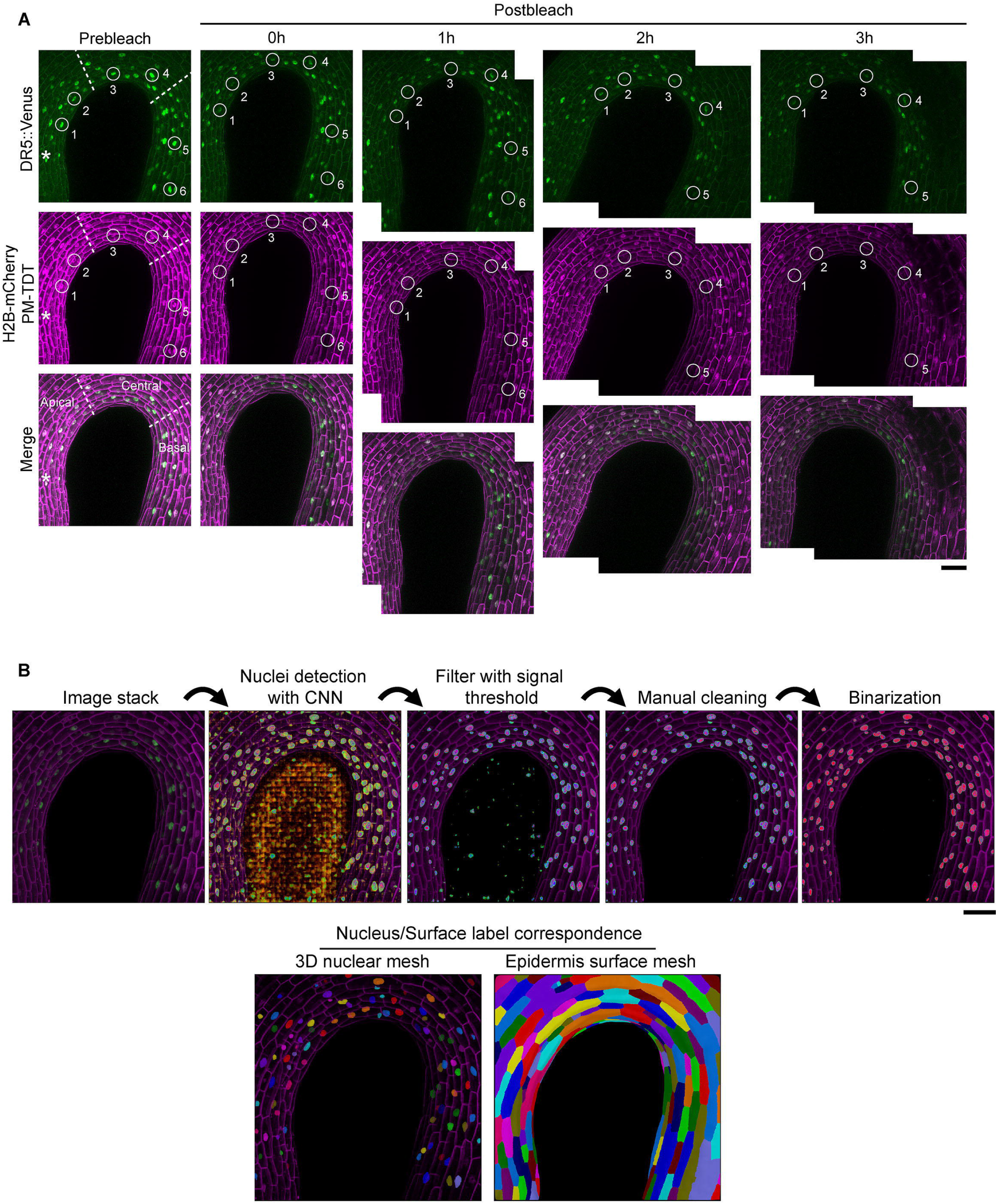
Fluorescence Recovery After Photobleaching (FRAP) analysis of DR5-Venus in the apical hook. **(A)** Confocal time-lapse images of a representative seedling (mock), depicting input data used for the recovery rate quantification presented in Figure 5K-N. Images show prebleach and postbleach fluorescence at indicated time points following photobleaching of DR5::Venus and H2B-mCherry. Top row depicts DR5::Venus, middle row depicts H2B-mCherry (nuclear) and PM-tdTomato (plasma membrane), and bottom row depicts merged channels. White circles indicate bleached nuclei, with numbers highlighting individual nuclei tracked during recovery. Asterisk indicates hypocotyl apex. Dotted lines demarcate the borders between the apical, central, and basal regions examined for recovery rates. Scale bar, 50 µm. **(B)** Workflow for nuclei segmentation and 3D mesh generation used to quantify recovery rates. Starting from the image stack, nuclei were detected using a convolutional neural network (CNN) for the DR5::Venus channel, filtered by signal threshold, manually cleaned, and binarized. The PM-tdTomato channel was used to extract epidermal cell surfaces, enabling assignment of nuclei to their respective cells for tracking over time. For detailed information, see methods section. Scale bar, 50 µm.

**Figure S20.**
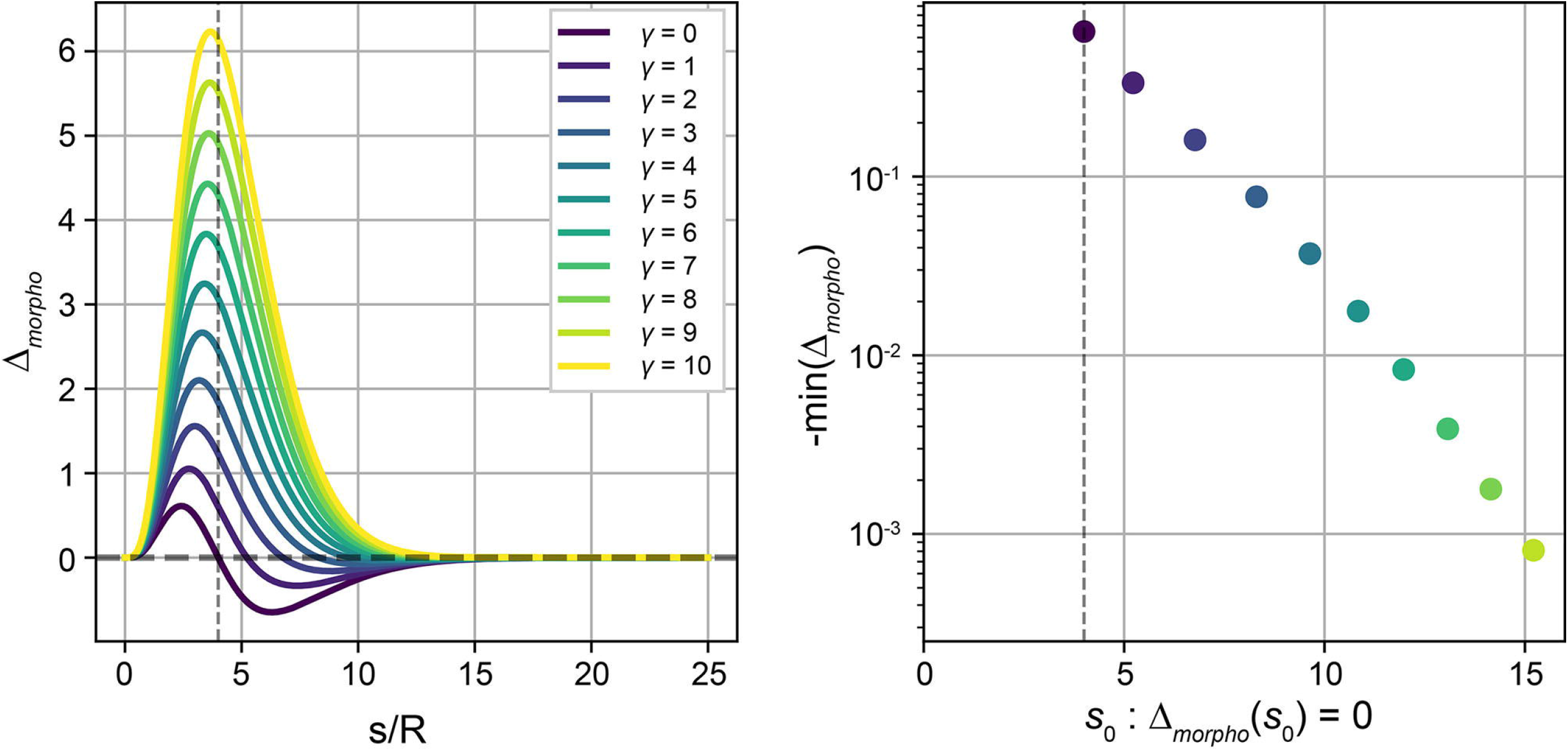
Characterization of Δ_morpho_ for various values of *γ*. **Left:** Profiles of <u>Δ</u>_morpho_ extracted from Eq. 3 of the main text, assuming <u>Δ</u>_morpho_ = Δ + *γκ*, for various values of the autotropic sensitivity *γ*. We note that here, for clarity and without loss of generality, we take *R* = 1 μm such that *R*<u>Δ</u>_morpho_ = <u>Δ</u>_morpho_. The longitudinal position of maximal curvature (*s*/*R* = 4) is indicated by a dashed vertical line. In the apical region (*s*/*R* < 4), strong competition between autotropism and <u>Δ</u>_morpho_ leads to large values of <u>Δ</u>_morpho_. Assuming <u>Δ</u>_morpho_ does not cause longitudinal cell shrinkage constrains this value to *R*<u>Δ</u>_morpho_ <2 (see SI, Eq.S19). This condition is violated for *γ* ≳ 3, setting an upper bound on *γ*. In the basal region (*s*/*R* > 4), <u>Δ</u>_morpho_ systematically changes sign from positive to negative, and therefore cannot remain positive along the full length of the organ. **Right:** For each *γ*, we plot the magnitude of the negative peak in the basal region of <u>Δ</u>_morpho_ against the longitudinal position *s*_0_ at which <u>Δ</u>_morpho_ = 0, and its sign changes. As *γ* increases, *s*_0_ shifts basipetally and the magnitude of the negative peak decreases. To allow <u>Δ</u>_morpho_ to remain positive within the hook, the sign change must occur below the hook in the hypocotyl (*s*_0_/*R* > 8). This condition sets a lower bound of *γ* ≳ 3, complementary to the upper bound from the apical region.

**Figure S21.**
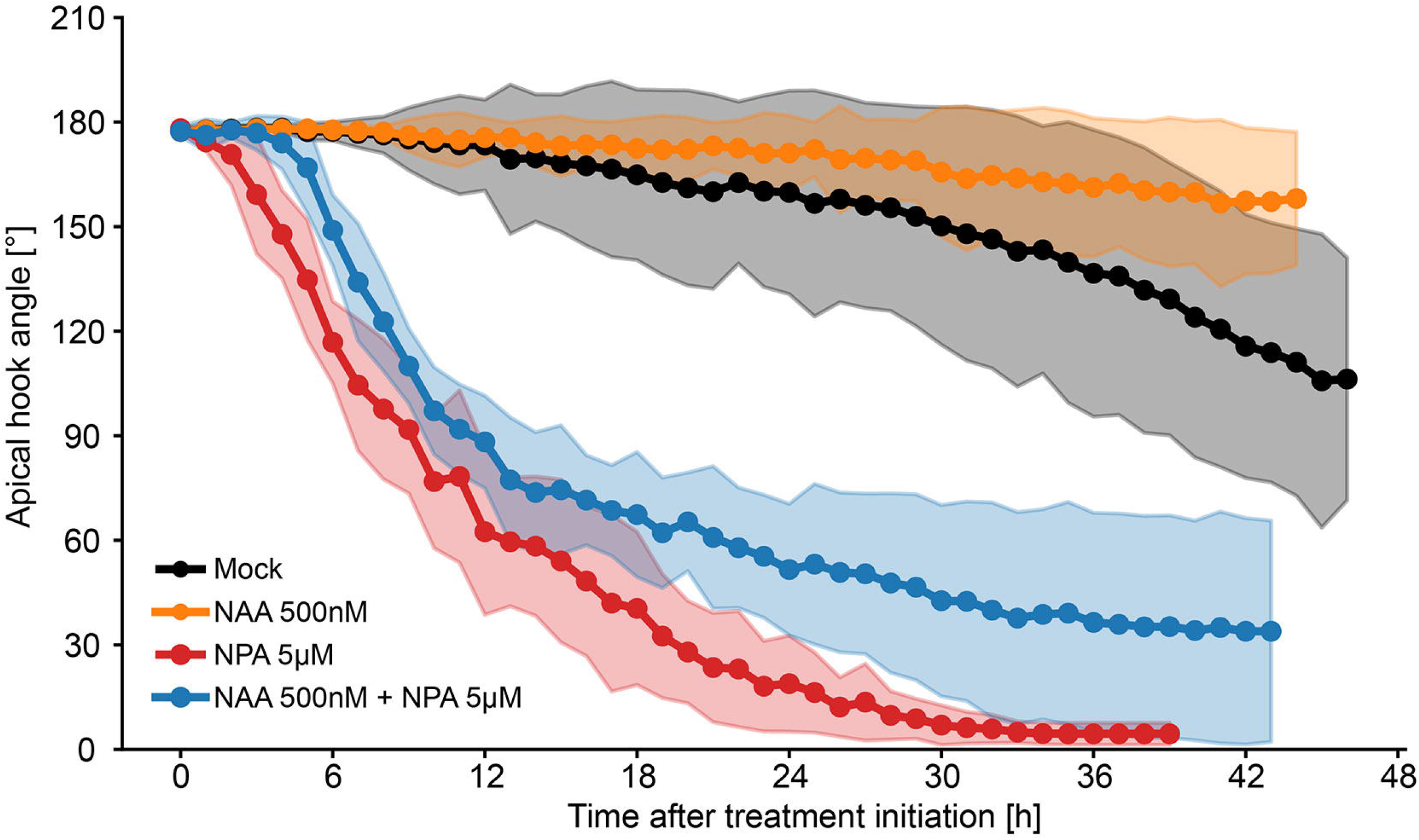
Hook opening induced by NPA persists upon co-treatment with NAA. Hook angle measured from the onset of the maintenance phase (designated *t* = 0 h). At 0 h, seedlings were transferred to mock, NAA (500 nM), NPA (5 µM), or combined NPA (5 µM) + NAA (500 nM) conditions. Curves show the temporal evolution of hook angle under each treatment. Shaded areas indicate mean ±SD. *n* ≥ 8

**Figure S22.**
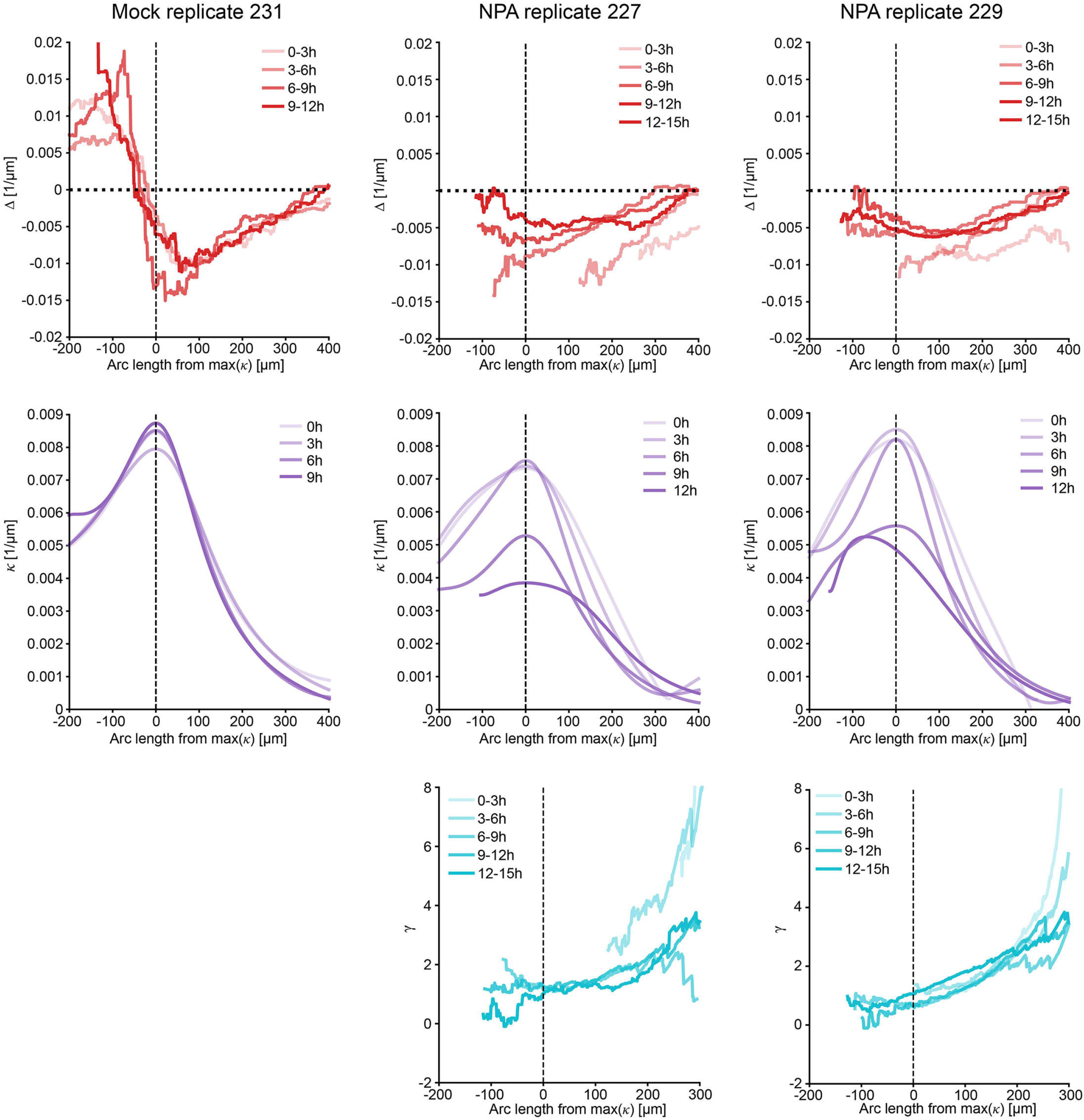
Replicate profiles of Δ, *κ* and y during mock and NPA treatments. Supplementary replicates corresponding to Fig. 7D–F. Profiles of Δ and *κ* over time during maintenance under mock and NPA 5 µM conditions are shown, with positions aligned to the point of maximal curvature, as in Fig. 7D-E. Corresponding *γ* profiles inferred from the Δ and *κ* data for each additional NPA-treated replicate are shown, as described for Fig. 7F.

**Figure S23.**
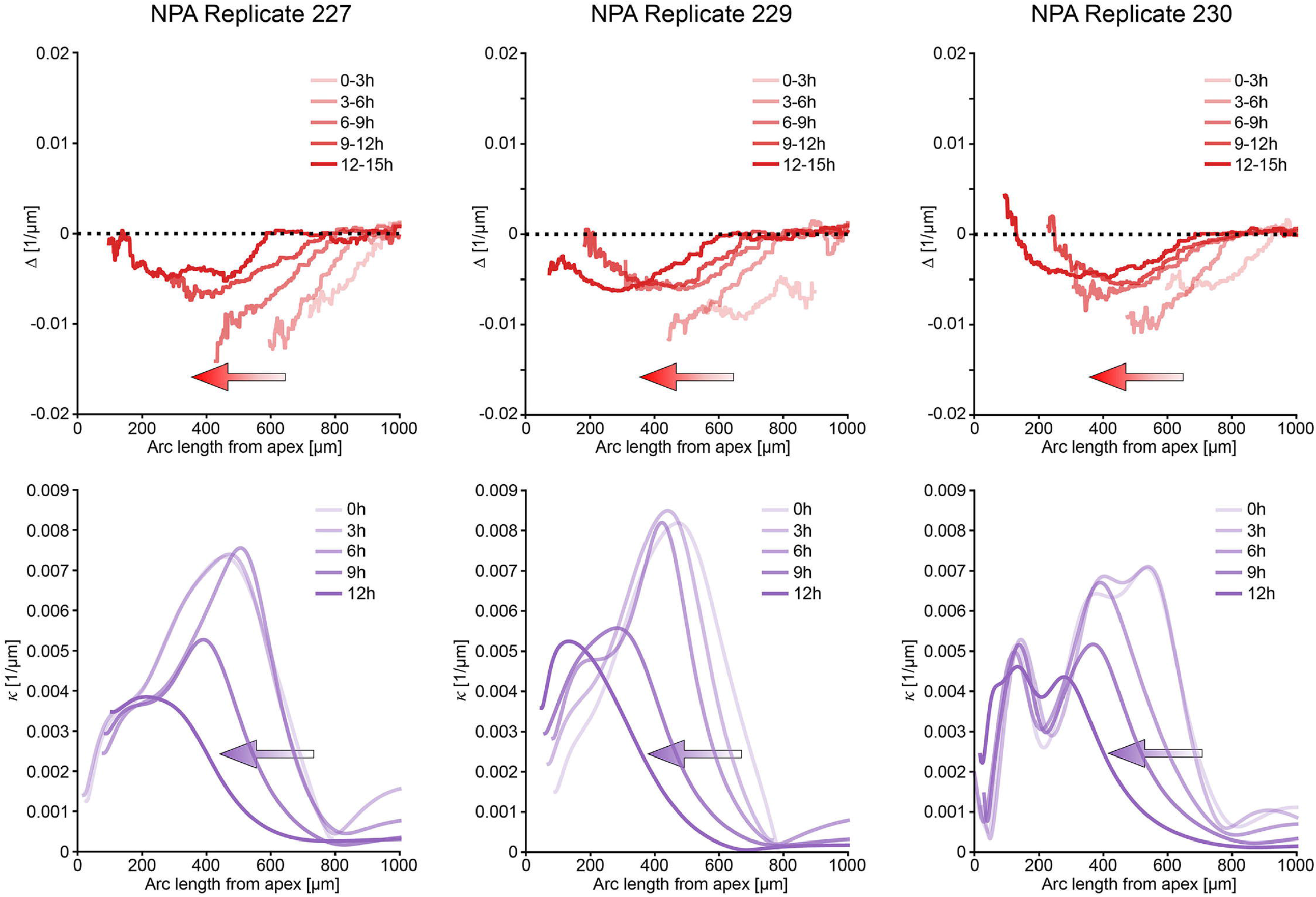
Basipetal displacement of Δ and *κ* profiles during NPA-induced hook opening. Profiles of Δ and *κ* over time during long-term NPA treatment (5 µM) for three independent biological replicates. In contrast to Fig. 7E, profiles are aligned with the hypocotyl apex set as *x*= 0, allowing visualization of spatial displacement along the organ. Over time, both the peak of negative Δ and the position of maximal curvature shift toward the apex. Temporal progression is indicated by color coding (Δ: early time points in light red, later time points in darker red; *κ*: early time points in light purple, later time points in darker purple). Arrows indicate the direction of profile displacement toward the apex.

## Supplementary movie legends

**Movie S1. Cell-level tracking during hook maintenance**

Morphing visualization of the same seedling shown in Fig. 1G, tracking individual cells over 12 h from the onset of hook maintenance (t = 0 h). The movie interpolates between consecutive 3 h intervals. Scale bar, 100 μm.

**Movie S2. Organ-region tracking during hook maintenance**

Morphing visualization of sequential images from Fig. 1G, showing kinematic tracking of organ regions over 12 h from the onset of hook maintenance (t = 0 h). The movie interpolates between consecutive 3 h intervals. Scale bar, 100 μm.

**Movie S3. Cell strain dynamics during hook maintenance**

Morphing visualization of sequential strain heatmaps from Fig. 2A, showing the same WT seedling over four consecutive 3 h intervals during apical hook maintenance. The movie interpolates between consecutive 3 h intervals. Scale bar, 100 μm.

**Movie S4. Kinematic model of WT maintenance**

Using stationary profiles of 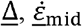 and initial shape *κ* that reproduces the hook shape, we integrate the shape dynamics via Eq.1. Left: profiles over arc length, normalized by the model organ’s radius (R = 100 μm). Right: hook shape dynamics. Black outlines on the organ indicate material segments which translate down the hook and elongate over time. Here, the profiles 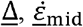 match the stationary shape *κ* via Eq.2, resulting in self similar dynamics as shown in Fig.2J. To prevent curvature drift at long times due to numerical errors, we applied autotropic error correction and used Eq.3 with *γ*= 5.

**Movie S5. Perturbation of kinematic model of WT maintenance (no negative** Δ **component)**

As in Movie S4, with *γ*= 0. Here, we retain only the positive part of Δ, which leads to overbending, as shown in Fig. 2K.

**Movie S6. Perturbation of kinematic model of WT maintenance (no positive** Δ = 0 **component)**

As in Movie S4, with *γ*= 0. Here, as shown in Fig. 2L, we retain only the negative part of Δ, which leads to opening followed by bending in the opposite direction.

**Movie S7. Perturbation of kinematic model of WT maintenance (**Δ = o)

As in Movie S4, with *γ* = 0. Here, as shown in Fig. 2M, setting Δ =0 reduces Eq.1 to a one-way wave equation for the curvature. This is illustrated on the Left graph by the propagation of the initial curvature profile.

**Movie S8. Perturbation of kinematic model of WT maintenance** 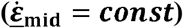

As in Movie S4, with *γ*= 0. Here, as shown in Fig. 2L, we keep the WT Δ and vary the midline strain rate to a constant value of 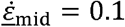 1/h along the organ. This leads to overbending.

**Movie S9. Curvature perturbation dynamics (***δκ* > 0)

As shown in Fig.6G, starting from the WT shape and Assuming a constant relative growth rate 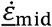 allows to solve Eq.4 and study the developmental robustness of the hook shape for different values of *γ*. Here, we show *γ* ∈ {0,1,10}, set a perturbation *Rδκ* = 0.3 over a small material segment, and display the shape dynamics in non dimensional time 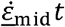. As the entire organ grows axially, the material segment is moving basipetally relative to the apex and acropetally relative to the base. For *γ*= 0, the perturbation is amplified by the elongation of the perturbed segment; For *γ*= 1, the decay of *κ* within the perturbed segment is perfectly balanced with the elongation of the segment, resulting in a constant excess angle and inclination of the model cotyledons; For *γ*= 10, the perturbation decay at a rate which increases with *γ*.

**Movie S10**.

As in Movie S9, but with a negative perturbation *Rδκ* = -0.3.

