## Supplementary material for "Robust Organ Shape During Growth Requires Local Morphogen Signaling and Global Curvature Feedback": SI

### – Supplementary Information: Mathematical Model –

### 1. Curvature dynamics

**A. A growing centerline.** In our modeling the hypocotyl geometry is described using its centerline $\boldsymbol{r}(s,t)$, characterized by its arc-length $s$ and time $t$, as well as a local material frame $\{\mathbf{d}_1(s,t), \mathbf{d}_2(s,t), \mathbf{d}_3(s,t)\}$ (1, 2). The curvature vector of the centerline $\boldsymbol{\kappa}(s,t)$ propagates the material frame in arc length via $\partial \mathbf{d}_i(s,t)/\partial s = \boldsymbol{\kappa}(s,t) \times \mathbf{d}_i(s,t)$ for $i \in \{1,2,3\}$. Assuming the organ is twistless and shearless allows us to identify the tangent to the centerline $\partial \boldsymbol{r}(s,t)/\partial s = \hat{\mathbf{T}}(s,t)$ as a material frame vector $\hat{\mathbf{T}}(s,t) = \mathbf{d}_3(s,t)$ and the curvature vector as $\boldsymbol{\kappa}(s,t) = \hat{\mathbf{T}}(s,t) \times \partial \hat{\mathbf{T}}(s,t)/\partial s$. For simplicity, we assume the shape dynamics are planar such that $\mathbf{d}_2$ is constant along the centerline and that the curvature vector can be expressed via the signed curvature $\kappa(s,t)$ using $\boldsymbol{\kappa}(s,t) = -\kappa(s,t)\mathbf{d}_2$.

To let the hypocotyl grow axially, we assume that for each time $t > 0$, the current arc length $s$ is related to an initial reference arc length $S_0$ by a smooth, strictly monotone map $s = s(S_0,t)$ with $\partial s/\partial S_0 \geq 1$, such that $s(S_0,0) = S_0$. This guarantees a bijective correspondence with a well defined inverse relation $S_0 = S_0(s,t)$. The coordinate $S_0$ thus acts as a material coordinate of the growing hypocotyl, and the axial stretch with respect to the reference configuration can be defined as $\lambda_0(S_0,t) \equiv \partial s(S_0,t)/\partial S_0$. The growth driven velocity of material points along the centerline can then be expressed by the dynamics of the axial stretch:

$$v(s,t) - v(0,t) \equiv \frac{ds(S_0,t)}{dt} = \frac{d}{dt}\int_0^{S_0} \lambda_0(S_0',t)dS_0' = \int_0^s \frac{d\lambda_0(S_0'(s',t),t)}{dt}\frac{dS_0'}{ds'}ds' = \int_0^s \frac{d}{dt}\ln\left(\lambda_0(S_0'(s',t),t)\right)ds' \quad \text{[S1]}$$

where in the third equality we change the integration variable from $S_0'$ to $s'$. Assuming $s = 0$ is a fixed point with $v(0,t) = 0$ and using the logarithmic strain measure $\varepsilon_{\mathrm{mid}}(s,t) \equiv \ln\left(\lambda(S_0(s,t),t)\right)$ simplifies Eq. S1 to:

$$v(s,t) = \int_0^s \dot{\varepsilon}_{\mathrm{mid}}(s',t)ds' \quad \text{[S2]}$$

where $\dot{\varepsilon}_{\mathrm{mid}}(s,t) \equiv d\varepsilon_{\mathrm{mid}}(s,t)/dt$ is the axial strain rate of the centerline.

**B. Differential axial growth.** We now wish to relate the distribution of axial growth across the cross section of the hypocotyl and the dynamics of curvature. For this, we express an explicit embedding of the material points in the volume of the hypocotyl using the centerline and the local material frame:

$$\boldsymbol{p}(s,x,y,t) = \boldsymbol{r}(s,t) + x\mathbf{d}_1(s,t) + y\mathbf{d}_2 \quad \text{[S3]}$$

where as before the centerline $\boldsymbol{r}(s,t)$ is constrained to the plane normal to $\mathbf{d}_2$. Since the cross section coordinates $x$ and $y$ do not depend on time, in Eq. S3 we implicitly assume that the cross section has a constant shape. For simplicity, we assume the shape is circular with radius $R$. The coordinates $(s,x,y)$ then act as dynamic curvilinear orthogonal coordinates that span the tubular volume of the hypocotyl at each time $t$. In these coordinates, the scale factor of arc length coordiante $s$ describes the local axial stretch with respect to the centerline:

$$\lambda(s,x,y,t) = \left|\frac{\partial \boldsymbol{p}(s,x,y,t)}{\partial s}\right| = 1 + x\kappa(s,t) > 0 \quad \text{[S4]}$$

where in the last equality we assume that the signed curvature is small such that $|R\kappa(s,t)| < 1$. The local total stretch with respect to the reference coordinate $S_0$ can then be found using the chain rule:

$$\lambda_{\mathrm{tot}}(s,x,y,t) = \left|\frac{\partial \boldsymbol{p}(s(S_0,t),x,y,t)}{\partial S_0}\right| = \frac{\partial s(S_0,t)}{\partial S_0}\left|\frac{\partial \boldsymbol{p}(s,x,y,t)}{\partial s}\right| = \lambda_0(S_0,t)\lambda(s,x,y,t) = \lambda_0(S_0,t)(1 + x\kappa(s,t)) \quad \text{[S5]}$$

where in the third equality we use Eq. S4. To decouple the curvature dynamics from the growth rate of the centerline, we introduce once more the logarithmic strain:

$$\varepsilon_{\mathrm{tot}}(s,x,y,t) \equiv \ln\left(\lambda_{\mathrm{tot}}(s,x,y,t)\right) = \ln\left(\lambda_0(S_0,t)\right) + \ln\left(1 + x\kappa(s,t)\right) \quad \text{[S6]}$$

Taking a time derivative of the strain then gives the local axial strain rate at each point in the volume of the hypocotyl:

$$\dot{\varepsilon}_{\mathrm{tot}}(s,x,y,t) = \dot{\varepsilon}_{\mathrm{mid}}(s,t) + \frac{x\dot{\kappa}(s,t)}{1 + x\kappa(s,t)} \quad \text{[S7]}$$

where we denote $\dot{X} \equiv dX/dt$ and use the axial strain rate of the centerline.

Following Eq. S7, the varying curvature can be related to the differential axial growth, namely, the gradient of the the axial strain rate across the cross section in the plane of bending $\partial \dot{\varepsilon}_{\mathrm{tot}}(s,x,y,t)/\partial x$. As this gradient does not depend on the perpendicular coordinate $y$, we can directly estimate it from experimental cellular measurements on the epidermis of a growing hypocotyl. In our analysis, we use the average gradient of strain rates on an experimental cross section, which can be related to our model by:

$$\left\langle \frac{\partial \dot{\varepsilon}_{\mathrm{tot}}}{\partial x} \right\rangle (s,t) = \frac{1}{2R}\int_{-R}^{R} \frac{\partial \dot{\varepsilon}_{\mathrm{tot}}(s,x',y,t)}{\partial x}dx = \frac{\dot{\varepsilon}_{\mathrm{tot}}(s,R,y,t) - \dot{\varepsilon}_{\mathrm{tot}}(s,-R,y,t)}{2R} = \frac{\dot{\kappa}(s,t)}{1 - (R\kappa(s,t))^2} \quad \text{[S8]}$$

where in the third equality we use Eq. S7. Rearranging Eq. S8 gives the curvature dynamics:

$$\dot{\kappa}(s,t) = \left(1 - (R\kappa(s,t))^2\right) \left\langle \frac{\partial \dot{\varepsilon}_{\text{tot}}}{\partial x} \right\rangle (s,t) \tag{S9}$$

We note that the time derivative of the signed curvature appearing in Eq. S9 is a material time derivative, such that:

$$\dot{\kappa}(s,t) = \frac{\partial \kappa(s,t)}{\partial t} + \frac{ds(S_0,t)}{dt}\frac{\partial \kappa(s,t)}{\partial s} = \frac{\partial \kappa(s,t)}{\partial t} + v(s,t)\frac{\partial \kappa(s,t)}{\partial s} \tag{S10}$$

where in the last equality we used Eqs. S1,S2.

**C. The normalized lateral gradient.** Lastly, we define the normalized lateral gradient of the axial strain rate, $\Delta(s,t)$, as a measure of diffenrtial axial growth:

$$\Delta(s,t) \equiv \frac{\dot{\kappa}(s,t)}{\dot{\varepsilon}_{\text{mid}}(s,t)} = \frac{1-(R\kappa(s,t))^2}{\dot{\varepsilon}_{\text{mid}}(s,t)} \left\langle \frac{\partial \dot{\varepsilon}_{\text{tot}}}{\partial x} \right\rangle (s,t) = \frac{1}{\dot{\varepsilon}_{\text{mid}}(s,t)} \left( \frac{\partial \dot{\varepsilon}_{\text{tot}}}{\partial x} \right)_{x=0} (s,t) \tag{S11}$$

where in the last equality we use Eq. S7. We then express the curvature dynamics via:

$$\dot{\kappa}(s,t) = \frac{\partial \kappa(s,t)}{\partial t} + v(s,t)\frac{\partial \kappa(s,t)}{\partial s} = \dot{\varepsilon}(s,t)\Delta(s,t) \tag{S12}$$

and the axial strain rate using:

$$\dot{\varepsilon}_{\text{tot}}(s,x,y,t) = \dot{\varepsilon}_{\text{mid}}(s,t)\left(1 + \frac{x\Delta(s,t)}{1+x\kappa(s,t)}\right) \tag{S13}$$

We note Eq. S12 gives Eq.1 in the main text. To extract $\dot{\varepsilon}(s,t)$ and $\Delta(s,t)$ from experimental data, we assume we are in the small curvature regime ($|R\kappa(s,t)| \ll 1$), and use Eqs. S11,S13 to obtain:

$$\dot{\varepsilon}_{\text{tot}}(s,x,y,t) \approx \langle \dot{\varepsilon}_{\text{tot}} \rangle (s,t) + x \cdot \left\langle \frac{\partial \dot{\varepsilon}_{\text{tot}}}{\partial x} \right\rangle (s,t) \approx \langle \dot{\varepsilon}_{\text{tot}} \rangle (s,t) \cdot (1 + x\Delta(s,t)) \tag{S14}$$

The normalized lateral gradient is commonly used in models of tropic movements (3, 4), where the additive decomposition of the differential growth term is assumed. Here, we decompose the normalized lateral gradient into two main contributions:

$$\Delta(s,t) = \Delta_{\text{morpho}}(s,t) - \gamma(s,t)\kappa(s,t) \tag{S15}$$

where $\Delta_{\text{morpho}}(s,t)$ is the local response to inhomogeneous concentrations of a morphogen, and $\gamma(s,t)$ is the local autotropic sensitivity which straightens the organ proportionally to its local curvature, as in Eq.3 in the main text. We note that the additive decomposition of the differential growth term can be related to a multi variable growth law. The axial growth rate of cells may be a function of various local properties $\{\mu_i\}$, such that $\dot{\varepsilon}_{\text{tot}}(\mu_1,...,\mu_N)$, where each property $\mu_i = \mu_i(s,x,y,t)$ is allowed to vary spatially in the organ. The properties $\{\mu_i\}$ may symbolize any field that affects the local growth rate, such as distributions of various morphogens or physical properties such as mechanical stress, Turgor pressure or cell wall properties. Following Eq. S11 and the chain rule, these dependencies give (omitting the $(s,t)$ dependencies for brevity):

$$\Delta \propto \left\langle \frac{\partial \dot{\varepsilon}_{\text{tot}}(\mu_1,...,\mu_N)}{\partial x} \right\rangle = \left\langle \sum_{i=1}^{N} \frac{\partial \dot{\varepsilon}_{\text{tot}}}{\partial \mu_i}\frac{\partial \mu_i}{\partial x} \right\rangle = \sum_{i=1}^{N} \left\langle \frac{\partial \dot{\varepsilon}_{\text{tot}}}{\partial \mu_i}\frac{\partial \mu_i}{\partial x} \right\rangle \tag{S16}$$

which effectively decomposes $\Delta$ into additive terms.
In addition, we note that in a strictly growing organ, the magnitude of the differential growth term is bounded. To show this, we set the axial strain rate in Eq. S13 as non-negative everywhere:

$$0 \le \dot{\varepsilon}_{\text{tot}}(s,x,y,t) = \dot{\varepsilon}_{\text{mid}}(s,t)\left(1 + \frac{x\Delta(s,t)}{1+x\kappa(s,t)}\right) \tag{S17}$$

Since $\dot{\varepsilon}(s,t) \ge 0$ we obtain:

$$\frac{x\Delta(s,t)}{1+x\kappa(s,t)} \ge -1 \tag{S18}$$

Since $x \in [-R,R]$ and $1 + x\kappa(s,t) > 0$ (as in Eq. S4) this inequality translates to:

$$0 \le |\Delta(s,t) + \kappa(s,t)| \le \frac{1}{R} \tag{S19}$$

such that $|R\Delta(s,t)| \le 2$, which we use in Figs.S6,S20.

### 2. Self similar growing hook

**A. Model assumptions.** We now turn to solve the curvature dynamics. For this, we place the origin $s = 0$ on the apex, and assume the growth rate depends only on the distance from the apex, denoting $\dot{\varepsilon}_{\text{mid}}(s,t) \equiv \dot{\varepsilon}_{\text{mid}}(s) = dv(s)/ds$ where $v(0) = 0$ (Eq. S2). We note that unlike in the main text, we do not always employ the underbar notation for time-independent profiles; instead, we explicitly indicate functional dependencies. We further assume the differential growth profile along the hook is defined by:

$$\Delta(s,t) = \Delta_{\text{morpho}}(s) - \gamma(s)\kappa(s,t) \tag{S20}$$

Here, we assume the spatial response to the morphogen $\Delta_{\text{morpho}}$ and the autotropic sensitivity $\gamma$ are functions of the distance from the apex alone. Substituting Eq. S20 in Eq. S12 while using $\dot{\varepsilon}(s)$ and $v(s)$ gives a linear PDE for the curvature $\kappa(s,t)$:

$$\frac{\partial\kappa(s,t)}{\partial t} + v(s)\frac{\partial\kappa(s,t)}{\partial s} = \dot{\varepsilon}_{\text{mid}}(s)\Delta_{\text{morpho}}(s) - \gamma(s)\dot{\varepsilon}_{\text{mid}}(s)\kappa(s,t) \tag{S21}$$

We note that this model includes the edge case $\gamma = 0$, for which $\Delta(s,t) = \Delta_{\text{morpho}}(s)$, reflecting the study cases presented in Fig.2 in the main text.

**B. Solution of the curvature dynamics.** We solve Eq. S21 by assuming the organ reaches a self similar shape with a stationary axial flow of material. In this non equilibrium steady state, the curvature profile remains constant in time even though the hypocotyl grows axially. We denote the self similar curvature profile as $\underline{\kappa}(s)$ and substitute it in Eq. S21. Since the partial time derivative disappears, Eq. S21 turns into a first order linear ODE:

$$v(s)\frac{d\underline{\kappa}(s)}{ds} = \dot{\varepsilon}_{\text{mid}}(s)\Delta_{\text{morpho}}(s) - \gamma(s)\dot{\varepsilon}_{\text{mid}}(s)\underline{\kappa}(s) \tag{S22}$$

The solution can be expressed using the integration factor:

$$\mu(s) = \exp\left(\int^{s} \frac{\gamma(s')\dot{\varepsilon}_{\text{mid}}(s')}{v(s')}ds'\right) = \exp\left(\int^{s} \gamma(s')\frac{d}{ds'}\ln\left(v(s')\right)ds'\right) \tag{S23}$$

Such that:

$$\underline{\kappa}(s) = \frac{1}{\mu(s)}\left(\int_0^{s} \frac{\dot{\varepsilon}_{\text{mid}}(s')\Delta_{\text{morpho}}(s')}{v(s')}\mu(s')ds' + c\right) \tag{S24}$$

where $c$ is an integration constant that can be found using an initial condition $\underline{\kappa}(0)$. We note that for $\gamma = const$ the integration factor (Eq. S23) simplifies to:

$$\mu_{\gamma=\text{const}}(s) = (v(s))^{\gamma} \tag{S25}$$

Assuming that the integral in Eq. S24 is well defined and the solution $\underline{\kappa}(s)$ exists allows us to solve the full PDE in Eq. S21. Substituting $\Delta_{\text{morpho}}(s)$ from Eq. S22 into Eq. S21 gives:

$$\frac{\partial\kappa(s,t)}{\partial t} + v(s)\frac{\partial}{\partial s}\left(\kappa(s,t) - \underline{\kappa}(s)\right) = -\gamma(s)\dot{\varepsilon}_{\text{mid}}(s)\left(\kappa(s,t) - \underline{\kappa}(s)\right) \tag{S26}$$

To continue, we denote $\delta\kappa(s,t) = \kappa(s,t) - \underline{\kappa}(s)$ as the difference between the current curvature and the steady state curvature. Since $\partial\delta\kappa(s,t)/\partial t = \partial\kappa(s,t)/\partial t$, Eq. S26 can be written as:

$$\frac{\partial\delta\kappa(s,t)}{\partial t} + v(s)\frac{\partial\delta\kappa(s,t)}{\partial s} = -\gamma(s)\dot{\varepsilon}_{\text{mid}}(s)\delta\kappa(s,t) \tag{S27}$$

Giving Eq.4 in the main text. Multiplying Eq. S27 by $\mu(s)$ (Eq. S23) and rearranging terms gives:

$$\left(\frac{\partial}{\partial t} + v(s)\frac{\partial}{\partial s}\right)\left(\mu(s)\delta\kappa(s,t)\right) = 0 \tag{S28}$$

which under the change of variable $\tau(s) = \int \frac{ds}{v(s)}$ is the one-way wave equation:

$$\left(\frac{\partial}{\partial t} + \frac{\partial}{\partial \tau}\right)\left(\mu(\tau)\delta\kappa(\tau,t)\right) = 0 \tag{S29}$$

The solution for $\delta\kappa(s,t)$ is then expressed by:

$$\delta\kappa(s,t) = \frac{1}{\mu(s)}F\left(\tau(s) - t\right) = \frac{1}{\mu(s)}F\left(\int\frac{ds}{v(s)} - t\right) \tag{S30}$$

where $F$ is a general function that depends on the initial condition, and $\int \frac{ds}{v(s)} - t = \text{const}$ are the characteristic curves of the PDE, tracing the axial movement of material points along the organ. The case with no autotropism $\gamma = 0$, such that $\mu(s) = 1$, depicts an organ for which the excess curvature $\delta\kappa$ is stretched along the arc length of the organ without change. With autotropism, since $\gamma, \dot{\varepsilon}_{\text{mid}}, v > 0$ for $s > 0$, Eq. S23 gives $d\ln(\mu(s))/ds > 0$, such that the excess curvature depicted in Eq. S30 decays with arc length. In the following section, we demonstrate this property using the simple case in which $\gamma$ is piecewise constant and the growth rate is constant. Lastly, we note that solving $\kappa(s,t)$ allows to expressed the hook angle trajectory analytically via:

$$\theta_{\text{hook}}(t) = \int_0^{L_{\text{hook}}} \kappa(s,t)ds \tag{S31}$$

where $L_{\text{hook}}$ is the length of the hook.

**C. Example: solutions with a constant growth rate.** If the growth rate is constant $\dot{\varepsilon}_{\mathrm{mid}}(s) = \dot{\varepsilon}_0$, the velocity is $ds/dt = v(s) = s\dot{\varepsilon}_0$, and the characteristic curves are given by $s(S_0, t) = S_0 e^{\dot{\varepsilon}_0 t}$. We can then use the material coordinate $S_0 = se^{-\dot{\varepsilon}_0 t}$ as the argument of $F$ in Eq. S30. Using Eqs. S25 and S30, for an initial shape of constant curvature $\kappa_0$ (i.e., a circular arc of radius $1/\kappa_0$) with constant $\gamma$, the initial condition for $\delta\kappa(s,t)$ is:

$$\delta\kappa(s,0) = \kappa_0 - \underline{\kappa}(s) = (s\dot{\varepsilon}_0)^{-\gamma} F(s) \tag{S32}$$

Since $s(S_0, t=0) = S_0$, this condition sets $F(S_0) = (S_0\dot{\varepsilon}_0)^{\gamma}(\kappa_0 - \underline{\kappa}(S_0))$. Substituting this expression and $S_0 = se^{-\dot{\varepsilon}_0 t}$ in Eq. S32 then allows to find the curvature for all times $t > 0$:

$$\kappa(s,t) = \underline{\kappa}(s) + \delta\kappa(s,t) = \underline{\kappa}(s) + e^{-\gamma\dot{\varepsilon}_0 t}(\kappa_0 - \underline{\kappa}(se^{-\dot{\varepsilon}_0 t})) \tag{S33}$$

showing that the local curvature decays exponentially in time to its steady state value for $\gamma > 0$. For a piecewise constant profile, where $\gamma(s) = 0$ for $s < s_0$ and $\gamma(s) = \gamma_0 > 0$ for $s \geq s_0$, we obtain $\mu(s) = (v(s)/v(s_0))^{\gamma(s)}$, and the excess curvature decays only for $s \geq s_0$. However, since the velocity is strictly positive, all material points will eventually surpass $s_0$ and have their excess curvature damped.

**D. Stability against perturbations.** Using our model of the curvature dynamics with a constant growth rate, we can ask how robust the self similar shape is to perturbations. We assume that the hypocotyl is presenting a self similar hook shape, and that at time $t = 0$ the shape is perturbed over a finite segment, such that $\delta\kappa(s,0)$ is non-zero only over the segment $[S_1, S_2]$ with $0 < S_1 < S_2$. We denote $\eta(s) \equiv \delta\kappa(s,0)$ and refer to $\eta(s)$ as the initial error in curvature. Such local curvature variations can occur, for example, following a noisy spatial morphogen pattern, a noisy axial growth rate profile or an external mechanical force that deforms the hook plastically. In this case, Eqs. S30,S32 show that $F(s) = (s\dot{\varepsilon}_0)^{\gamma}\eta(s)$ and that $\delta\kappa(s,t)$ can be expressed by:

$$\delta\kappa(s,t) = (s\dot{\varepsilon}_0)^{-\gamma} F(se^{-\dot{\varepsilon}_0 t}) = e^{-\gamma\dot{\varepsilon} t}\eta\left(se^{-\dot{\varepsilon}_0 t}\right) \tag{S34}$$

Since $\eta(s)$ is non zero over the segment $[S_1, S_2]$ at $t = 0$, $\delta\kappa(s,t)$ is non zero over the segment $[S_1 e^{\dot{\varepsilon}_0 t}, S_2 e^{\dot{\varepsilon}_0 t}]$ for $t > 0$, and the length over which the error propagates is growing exponentially following $e^{\dot{\varepsilon}_0 t}(S_2 - S_1)$. Focusing on the developing hook shape after the perturbation, the propagating error in curvature can be expressed by the total excess angle added to the shape:

$$\delta\theta(t) = \int_{S_1 e^{\dot{\varepsilon}_0 t}}^{S_2 e^{\dot{\varepsilon}_0 t}} \delta\kappa(s,t) ds = e^{-\gamma\dot{\varepsilon}_0 t} \int_{S_1 e^{\dot{\varepsilon}_0 t}}^{S_2 e^{\dot{\varepsilon}_0 t}} \eta\left(se^{-\dot{\varepsilon}_0 t}\right) ds = e^{-(\gamma-1)\dot{\varepsilon}_0 t} \int_{S_1}^{S_2} \eta\left(S_0\right) dS_0 \tag{S35}$$

where in the second equality we use Eq. S34 and in the last equality we changed the integration variable to $S_0 = se^{-\dot{\varepsilon}_0 t}$. This result can be rewritten via the initial excess angle:

$$\delta\theta(t) = e^{-(\gamma-1)\dot{\varepsilon}_0 t}\delta\theta(0) \tag{S36}$$

We find that the excess angle either diverges or decays over time depending on the value of $\gamma$, the autotropic sensitivity. If $\gamma < 1$, the error diverges. This means that models with low autotropic coefficients are unstable to perturbations. If $\gamma = 1$, the excess angle remains constant as the decay in curvature is perfectly balanced with the elongation of the curved segment. If $\gamma > 1$, the perturbation disappears with the decay time:

$$\tau_{decay} = \frac{1}{(\gamma - 1)\dot{\varepsilon}_0} \tag{S37}$$

This stability criterion of self similar curvature dynamics reproduces the stability criterion of gravitropic models against passive orientation drift detailed in (3). The decay time in Eq. S37 can also characterize the transition time between two self similar profiles if $\Delta_{\mathrm{morpho}}(s,t)$ is allowed to change in time.
